# Genetic association of *TMPRSS2* rs2070788 polymorphism with COVID-19 Case Fatality Rate among Indian populations

**DOI:** 10.1101/2021.10.04.463014

**Authors:** Rudra Kumar Pandey, Anshika Srivastava, Prajjval Pratap Singh, Gyaneshwer Chaubey

## Abstract

SARS-CoV2, the causative agent for COVID-19, an ongoing pandemic, engages the ACE2 receptor to enter the host cell through S protein priming by a serine protease, TMPRSS2. Variation in the TMPRSS2 gene may account for the difference in population disease susceptibility. The haplotype-based genetic sharing and structure of *TMPRSS2* among global populations have not been studied so far. Therefore, in the present work, we used this approach with a focus on South Asia to study the haplotypes and their sharing among various populations worldwide. We have used next-generation sequencing data of 393 individuals and analysed the TMPRSS2 gene. Our analysis of genetic relatedness for this gene showed a closer affinity of South Asians with the West Eurasian populations therefore, host disease susceptibility and severity particularly in the context of *TMPRSS2* will be more akin to West Eurasian instead of East Eurasian. This is in contrast to our prior study on ACE2 gene which shows South Asian haplotypes have a strong affinity towards West Eurasians. Thus ACE2 and TMPRSS2 have an antagonistic genetic relatedness among South Asians. We have also tested the SNP’s frequencies of this gene among various Indian state populations with respect to the case fatality rate. Interestingly, we found a significant positive association between the rs2070788 SNP (G Allele) and the case fatality rate in India. It has been shown that the GG genotype of rs2070788 allele tends to have a higher expression of TMPRSS2 in the lung compared to the AG and AA genotypes, thus it might play a vital part in determining differential disease vulnerability. We trust that this information will be useful in underscoring the role of the TMPRSS2 variant in COVID-19 susceptibility and using it as a biomarker may help to predict populations at risk.

## 1. Introduction

COVID-19 is an ongoing pandemic that has cost millions of lives worldwide, caused by the SARS-CoV2 virus of the Beta Family. Along with ACE2 (Angiotensin-converting enzyme 2) which acts as a receptor, TMPRSS2 (Transmembrane protease, serine 2), a serine protease, is also involved in virus entry the host cell through S Protein priming (1,2). Along with SARS-CoV-2, the Influenza virus, as well as the various human coronaviruses such as HCoV-229E, MERS-CoV, and SARS-CoV, have been identified to utilize this protein for cell entrance (3). Serine proteases have been linked to a variety of physiological and pathological processes. Androgenic hormones were shown to upregulate this gene in prostate cancer cells, while androgen-independent prostate cancer tissue was found to downregulate it (4). Northern blots analysis has revealed that in mice TMPRSS2 is mainly expressed in the kidney and prostate, whereas in humans, TMPRSS2 is largely expressed in the prostate, salivary gland, stomach and colon (5). TMPRSS2 is also expressed in the epithelia of the respiratory, urogenital and gastrointestinal tracts according to in-situ hybridization investigations performed on mice embryos and adult tissues (5).

The impact of the COVID-19 crisis is not uniform across ethnic groups. Patients from different ethnic backgrounds suffer disproportionately (6). Discrepancies in infection as well as case fatality rates (CFR) could be due to multiple reasons e.g., differences in quarantine and social distancing policies, access to medical care, reliability & coverage of epidemiological data, and population age structure, which shows that mortality is greater among the elderly and those with comorbidity (7,8). However, many young and healthy people have also lost their lives due to rapid cytokine storms (9). It is important to note that these factors do not appear to account for all the disparities noticed among groups, and there are significant gaps that require the scientific community’s attention to propose and test theories that will assist us in better understanding the disease etiology. This is even more important, keeping in mind that the number of cases and deaths may be poorly reported in some populations however, countries with strict standards for the collection and presentation of epidemiological data suggest that human variation in genetic makeup may account for differential susceptibility and severity in disease outcomes among different populations (10). There is evidence that supports the role of ACE2 gene variations in susceptibility to COVID-19 in Indian populations (11,12). However, little is known regarding the genetic structure of TMPRSS2 haplotypes among South Asian populations, a detailed analysis of the sequence data of TMPRSS2 gene from world populations may unveil its haplotype sharing, which may help understand the role of TMPRSS2 in disease susceptibility globally. Given the relevance of the TMPRSS2 gene in the SARS-CoV-2 infection process, COVID-19 infection and severity pattern may be directly linked to elevated TMPRSS2 gene expression, resulting in varying disease susceptibility outcomes in various communities globally. However, the role of TMPRSS2 polymorphism for disease susceptibility in the Indian populations is largely unexplored and this needs to be examined. Therefore, in the current study, we analysed the haplotype structure of TMPRSS2 focusing on South Asia and its genetic markers that could be responsible for changes in the gene’s expression in the lungs tissue and, correlate it with epidemiological data on COVID-19 for any existing association among Indian population.

## 2. Material and Methods

The TMPRSS2 gene haplotype analysis for various world populations was done using NGS data from (13). PLINK 1.9 was used to extract sequences from the dataset for different populations (14). After excluding samples from Sahul and Africa, as well relatives up till second-degree, a total of 393 samples and 795 SNPs were observed and were used further for study **(Supplementary Table 1 and 2)**. The plink file was converted to fasta (ped to IUPAC) by a customized script (15). For the purpose of phasing, Fst calculation, Population-wise genetic distances calculation, and generation of Network and Arlequin input file, DNAsp was used (16). MEGA X was used to construct an Fst based Neighbour-joining tree (17). To calculate Nei’s genetic and average pairwise distance, Arlequin 3.5 was used and plotted on a graph by R V3.1 (18,19). Network v5 and network publisher were employed to draw the median-joining network while total and prevalent haplotypes in TMPRSS2 gene for each population were calculated using XML file generated through Arlequin 3.5 (18,20).

For the association study, we searched for the studies on TMPRSS2 variants reported in the literature elsewhere in relation to COVID-19 susceptibility (4,21–41). We obtained a total of 5 SNPs (rs2070788, rs734056, rs12329760, rs2276205, and rs3787950) was observed in our data and studied subsequently in detail. Data from the Estonian Biocentre (42–45), data from phase 3 of the 1,000 Genomes Project (46), and our new genotyped samples from several Indian states were used to calculate the frequency of each of these SNPs among various Indian populations using plink 1.9. State-wise frequency maps for rs2070788 and COVID-19 CFR among the Indian population were made by https://www.datawrapper.de/. and worldwide spatial distribution of rs2070788 was generated from the *PGG.SNV* toolkit using 1000 genome samples (47). The regression plots for statewise allele frequency Vs the CFR were constructed using https://www.graphpad.com/quickcalcs/linear1/ and further validated by the Microsoft excel regression calculations. We also performed Pearson’s correlation coefficient test (48) at a 95 percent confidence interval and 1,000 bootstrapping (2,000,000 seeds) for a two-tailed significance test to verify our results by using, SPSS (ver 26). The LD map and aggregate frequency of haplotypes carrying rs2070788 (G allele) were calculated for each of the populations by Haploview (49).

## 3. Result and Discussion

TMPRSS2 is a serine protease enzyme that is encoded in humans by the TMPRSS2 gene that is located on chromosome 21q22.3. (50). This protein aids in virus entry into host cells, such as the influenza virus, and human coronaviruses such as HCoV-229E, MERS-CoV, SARS-CoV, and SARS-CoV-2 by proteolytically cleaving and then activating the viral envelope glycoproteins (51), and thus can be inhibited by TMPRSS2 inhibitor (1). Genetic variation in this gene may account for differential vulnerability for COVID-19 disease among diverse populations, therefore, in the present study with our major focus being on South Asia.

We analyzed TMPRSS2 gene sequence data among world populations by haplotype-based approach for comparison among the various groups. *F*st based neighbour Joining (NJ) tree showed the clustering of South Asians with the West Eurasian populations (Caucasus, West Asia, Europe, and Central Asia) **(Figure 1A)**. Similarly, the Average Pairwise differences analysis showed smaller diversity and genetic distance between populations, among East and West Eurasians, while greater diversity and genetic distance was observed between East and West Eurasian populations. The lowest diversity was found within West Asia & the American population **(Figure 1B)**. A median-joining (MJ) network analysis of the TMPRSS2 gene revealed that there are 499 haplotypes throughout this gene among the examined populations, with prevalent haplotypes (Hap 34, Hap 48, Hap 75, Hap 98, and Hap 260), each having ≥10 individuals. Haplotypes 48 and 75 were found to be more common in Europe, while haplotypes 98 and 260 were observed to be more common in Siberia. Haplotype 34 was frequent in Southeast Asia, followed by Central Asia **(Supplementary Table 3A and Supplementary Figure 1)**. Altogether, South Asian populations carry 47 haplotypes, among which 6 are shared (Hap_34, Hap_48, Hap_78, Hap_112, Hap_219, and Hap_260) with other continental populations while the rest are unique to South Asia. Among the shared haplotypes, five are shared with the West Eurasian populations, whereas only a single haplotype is shared with the East Eurasian populations. **(Figure 1C and Supplementary Table 3B)**. The haplotype sharing, as well as Fst analysis, are consistent with the West Eurasian affiliation of the majority of South Asian TMPRSS2 haplotypes **(Figure 1C and Figure 1A)**. Therefore, the host susceptibility of SARS-CoV-2 for TMPRSS2 gene among South Asians is most likely expected to be similar to West Eurasian rather than that of East Eurasians. In contrast with this, our previous study on the ACE2 gene has shown the strong affinity of South Asian haplotypes with the East Eurasians (11,12). Thus, for the South Asians, ACE2 and TMPRSS2 have an antagonistically genetic relatedness. As a result, it’s worth proposing that the South Asian population’s susceptibility to SARS-CoV-2 will fall somewhere between West and East Eurasian people, which is most likely the cause of the moderate susceptibility.

**Fig 1.**
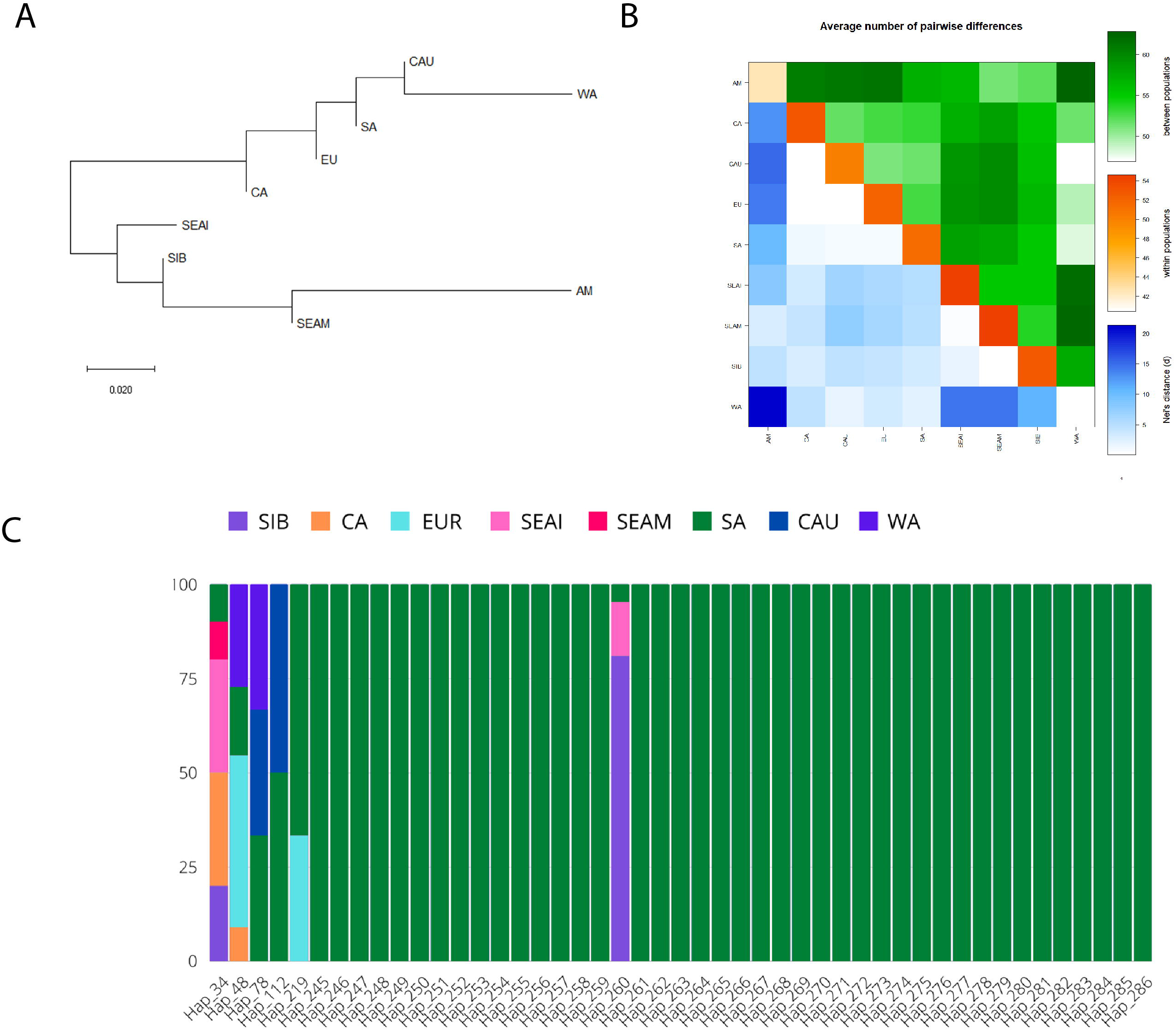
**(A)** Neighbour-Joining (NJ) tree based on Fst distance, showing genetic relationship for TMPRSS2 gene among the studied population. **(B)** Matrix showing average paired variation for TMPRSS2 gene, between the population (green) in the upper triangle, within-population (orange) along diagonal, and Nei’s distance between populations are shown (blue) in the lower triangle. The obtained value for different variables is directly proportional to the color gradient. **(C)** The stacked bar-plot represents 47 haplotypes observed in TMPRSS2 Gene among South Asian populations. Frequency and sharing for each haplotype with South Asia and to other geographic regions are indicated with different coloured bars.

There has not been any association study so far on the TMPRSS2 variants in relation to COVID-19 among Indian Populations. Therefore, we calculated groupwise allele frequencies in Indian populations for all the 5 SNPs (rs2070788, rs734056, rs12329760, rs2276205, and rs3787950) observed in our data. The linear regression analysis was carried out for these SNP’s for spatial frequency in India with COVID-19 CFR among various Indian states **(Supplementary Table 4 A, B and 5)**. The Regression Analysis showed a significant positive correlation for rs2070788 SNP (G allele), between allele frequency and case fatality rate (p < 0.05). Higher CFR was observed where the allele frequency is higher and vice versa **(Figure 2A and B)**. The goodness of fit (R^2^) explained 33.82% of the variation **(Figures 2C)**. Because this is an active pandemic with changing numbers of infected and dead patients, we confirmed our findings at different timelines (latest up to August 2021). The recent data backs up the previous observation with no substantial difference between the outcomes, to further validate our results we performed the Pearson correlation coefficient test which shows a significant positive correlation with r = .582, p = 0.029, thus supporting the previous observation of strong positive association **(Table 1)**.

**FIGURE 2.**
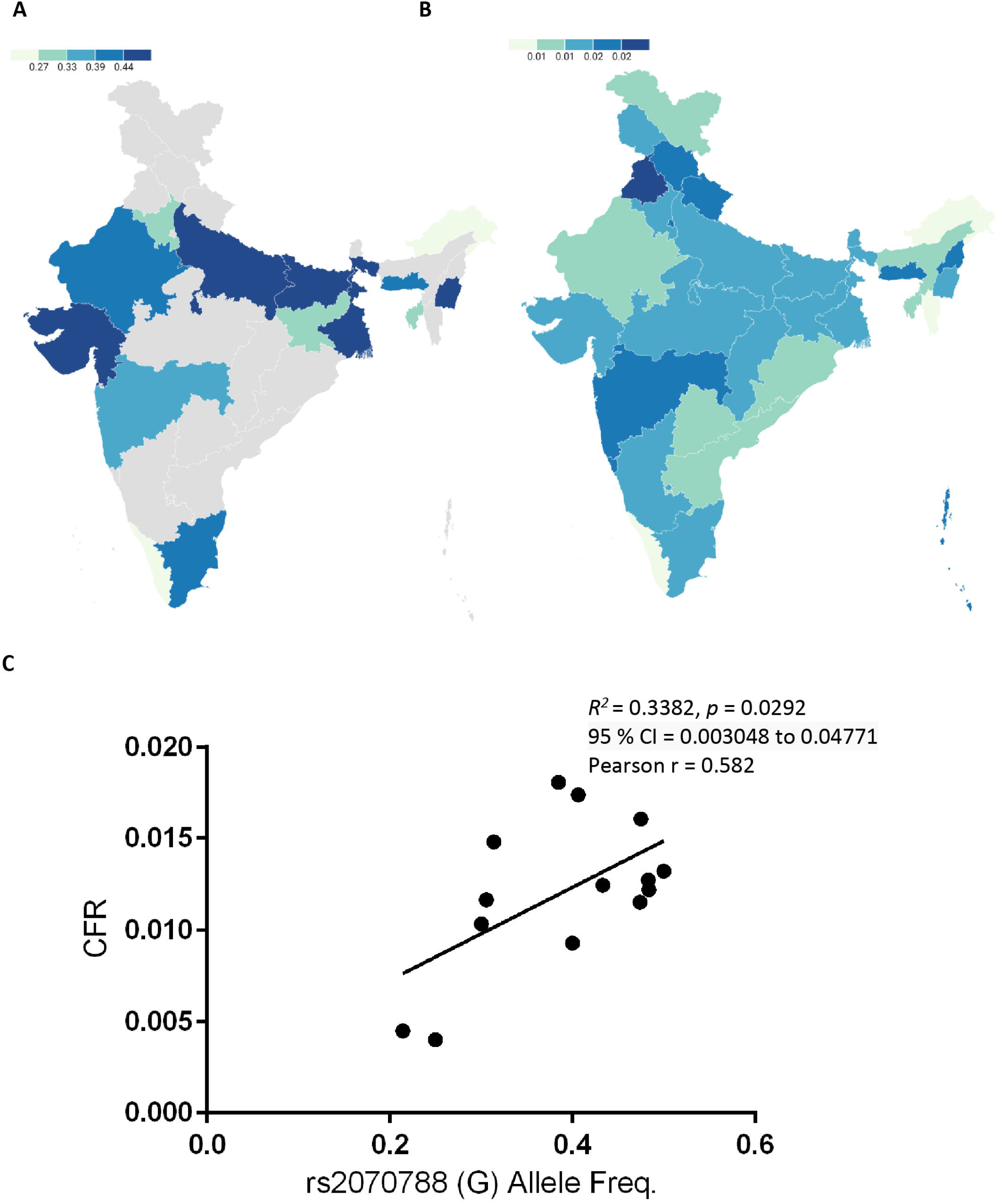
**(A)** frequency map (%) showing the spatial distribution of allele rs2258666 among Indian populations. Grey colour marks the absence of data. **(B)** The Map of state-wise frequency (%) of case-fatality rate (CFR) (updated till 30^th^ August 2021). **(C)** The linear regression analysis graph showing the goodness of fit and Pearson correlation coefficient for the allele frequency vs. CFR.

**TABLE 1.** Outcome of tests conducted for statistical significance at different timelines of the pandemic in India.

| Observation | Linear regression |  | Pearson's correlation |  |
| --- | --- | --- | --- | --- |
| rs2070788 | R square | p-value | r | p-value |
| June 2021_CFR | 0.3382 | 0.0292 | 0.582 | 0.029 |
| July 2021_CFR | 0.3097 | 0.0387 | 0.557 | 0.039 |
| August 2021_CFR | 0.2888 | 0.0475 | 0.537 | 0.047 |

Tmprss2 expression in the lungs was reported to be higher in the rs2070788 GG genotype than those in the AA and AG genotype (52) thus, the G allele may contribute to severe consequences in SARS-COV2 infection in populations with high frequency. We found that G allele frequency in India ranges from 20% to 50%, with the mean frequency of 39%, lowest being in Arunachal Pradesh and highest in Bihar which is in accordance as per data observed which clearly shows Arunachal Pradesh is among those states that show lowest CFR while Bihar and other states are among higher CFR rate **(Supplementary Table 4A and B)**. Thus this may explain the disparity in severity of pandemic among various Indian states **(Figure 2 B)**. Being an androgen-sensitive gene TMPRSS2 is known to mediate sex-related effects and rs2070788 SNP seems to play an important role (53). Higher expression of TMPRSS2 in males might make them more prone to virus fusion and could explain high COVID-19 mortality in males (54,55).

For Linkage disequilibrium (LD) analysis, LD plots were made for each population focussing on rs2070788 and nearby SNPs on that haplotype. LD blocks of various sizes were observed among Central Asians, Caucasians, Europeans, South Asians, Siberians, and West Asians. The highest LD level was found in Americans. **(Supplementary Figure 2)**. We also calculated aggregate haplotypes frequency which are in LD carrying rs2070788 (G allele), in each population presented in **(Supplementary Table 6)**. Considerable levels of variation in haplotype frequency were observed among the populations. The highest haplotype frequency was observed in America (0.654), while the lowest haplotype frequency was recorded in Southeast Asia Island (0.322), these findings are consistent with epidemiological data available on COVID-19 which clearly shows that the American population has the most number of cases and death while Southeast Asians are much below in the list. We also looked for worldwide distribution of rs2070788 (G allele) from 1000 genome data **(Supplementary Table 7 and Supplementary Figure 3)** and found consistent with the previous observation, rs2070788 (G allele) frequency was highest in Americans (0.49), while lowest in African (0.27) and East Asians (0.36) populations, this may explain high fatality in among Americans populations while African and East Asians being least affected. Low severity among East Asians could be due to adaptation at many genes that engage with coronaviruses, also including the SARS-CoV-2, which began 25,000 years back for coronaviruses, or a related virus outbreak in East Asia at that time (56).

## 4. Conclusion

In conclusion for the first time, we have shown closer affinity of South Asians with the West Eurasian populations for TMPRSS2 gene. Hence, hot disease susceptibility in context of TMPRSS2 will be more likely similar to West Eurasian populations. This is in contrast to our prior study on the ACE2 gene, which showed closer genetic affinity of South Asian haplotypes with Easts Eurasians. Thus, for South Asians, ACE2 and TMPRSS2 have an antagonistic genetic relationship. So, it’s worth proposing that the susceptibility of the South Asian population to SARS-CoV-2 will fall somewhere between West and East Eurasian populations, which is most likely the source of the moderate susceptibility. We also found a genetic association between rs2070788 and CFR among various Indian populations. This information could be used as a genetic biomarker to predict susceptible populations, which may be very useful during the epidemic in policymaking and making better resource allocation.

## Author Contributions

GC and RKP conceived and designed this study. RKP, AS, and PPS analysed the data. RKP, AS, PPS, and GC wrote the manuscript. All authors contributed to the article and approved the submitted version

## Acknowledgments

This work is supported by Faculty IOE grant BHU (6031). RKP is supported by the UGC-Non-NET fellowship, AS is supported by UGC-CAS fellowship and PPS is supported by CSIR fellowship.

## Data Availability Statement

All datasets generated for this study are included in the article/Supplementary Material.

---

**Supplementary Figure 1** The median-joining network of TMPRSS2 gene. The circle size determines the number of samples with a certain haplotype. The five most common haplotypes are marked.

**Supplementary Figure 2** LD (linkage disequilibrium) maps of the TMPRSS2 gene, focusing on rs2070788 and its haplotype, in world populations. Shading from white to red indicates the intensity of r2 from 0 to 1. Strong LD is represented by a high percentage (>80) in darker red squares.

**Supplementary Figure 3** The spatial distribution of SNP rs2070788 from 1000 genome data

## Refrences

1. Hoffmann M, Kleine-Weber H, Schroeder S, Krüger N, Herrler T, Erichsen S, et al. SARS-CoV-2 Cell Entry Depends on ACE2 and TMPRSS2 and Is Blocked by a Clinically Proven Protease Inhibitor. Cell. 2020 Apr 16;181(2):271–280.e8.

2. Zhou P, Yang X-L, Wang X-G, Hu B, Zhang L, Zhang W, et al. A pneumonia outbreak associated with a new coronavirus of probable bat origin. Nature. 2020 Mar;579(7798):270–3.

3. Shen LW, Mao HJ, Wu YL, Tanaka Y, Zhang W. TMPRSS2: A potential target for treatment of influenza virus and coronavirus infections. Biochimie. 2017 Nov 1;142:1–10.

4. Mollica V, Rizzo A, Massari F. The pivotal role of TMPRSS2 in coronavirus disease 2019 and prostate cancer. Future Oncol. 2020 Sep 1;16(27):2029–33.

5. Vaarala MH, Porvari KS, Kellokumpu S, Kyllönen AP, Vihko PT. Expression of transmembrane serine protease TMPRSS2 in mouse and human tissues. J Pathol. 2001;193(1):134–40.

6. Webb Hooper M, Nápoles AM, Pérez-Stable EJ. COVID-19 and Racial/Ethnic Disparities. JAMA. 2020 Jun 23;323(24):2466–7.

7. Ejaz H, Alsrhani A, Zafar A, Javed H, Junaid K, Abdalla AE, et al. COVID-19 and comorbidities: Deleterious impact on infected patients. J Infect Public Health. 2020 Dec 1;13(12):1833–9.

8. Sanyaolu A, Okorie C, Marinkovic A, Patidar R, Younis K, Desai P, et al. Comorbidity and its Impact on Patients with COVID-19. SN Compr Clin Med. 2020 Aug 1;2(8):1069–76.

9. Muschitz C, Trummert A, Berent T, Laimer N, Knoblich L, Bodlaj G, et al. Attenuation of COVID-19-induced cytokine storm in a young male patient with severe respiratory and neurological symptoms. Wien Klin Wochenschr [Internet]. 2021 Apr 27 [cited 2021 Aug 28]; Available from: https://doi.org/10.1007/s00508-021-01867-2

10. SeyedAlinaghi S, Mehrtak M, MohsseniPour M, Mirzapour P, Barzegary A, Habibi P, et al. Genetic susceptibility of COVID-19: a systematic review of current evidence. Eur J Med Res. 2021 May 20;26(1):46.

11. Srivastava A, Bandopadhyay A, Das D, Pandey RK, Singh V, Khanam N, et al. Genetic Association of ACE2 rs2285666 Polymorphism With COVID-19 Spatial Distribution in India. Front Genet. 2020;11:1163.

12. Srivastava A, Pandey RK, Singh PP, Kumar P, Rasalkar AA, Tamang R, et al. Most frequent South Asian haplotypes of ACE2 share identity by descent with East Eurasian populations. PLOS ONE. 2020 Sep 16;15(9):e0238255.

13. Pagani L, Lawson DJ, Jagoda E, Mörseburg A, Eriksson A, Mitt M, et al. Genomic analyses inform on migration events during the peopling of Eurasia. Nature. 2016 Oct;538(7624):238–42.

14. Purcell S, Neale B, Todd-Brown K, Thomas L, Ferreira MAR, Bender D, et al. PLINK: A Tool Set for Whole-Genome Association and Population-Based Linkage Analyses. Am J Hum Genet. 2007 Sep 1;81(3):559–75.

15. Sander N, Abel GJ, Bauer R, Schmidt J. Visualising migration flow data with circular plots [Internet]. Vienna Institute of Demography Working Papers; 2014 [cited 2021 Aug 28]. Report No.: 2/2014. Available from: https://www.econstor.eu/handle/10419/97018

16. Rozas J, Ferrer-Mata A, Sánchez-DelBarrio JC, Guirao-Rico S, Librado P, Ramos-Onsins SE, et al. DnaSP 6: DNA Sequence Polymorphism Analysis of Large Data Sets. Mol Biol Evol. 2017 Dec 1;34(12):3299–302.

17. Kumar S, Stecher G, Li M, Knyaz C, Tamura K. MEGA X: Molecular Evolutionary Genetics Analysis across Computing Platforms. Mol Biol Evol. 2018 Jun;35(6):1547–9.

18. Excoffier L, Lischer HEL. Arlequin suite ver 3.5: a new series of programs to perform population genetics analyses under Linux and Windows. Mol Ecol Resour. 2010;10(3):564–7.

19. R: The R Project for Statistical Computing [Internet]. [cited 2021 Aug 28]. Available from: https://www.r-project.org/

20. Bandelt HJ, Forster P, Röhl A. Median-joining networks for inferring intraspecific phylogenies. Mol Biol Evol. 1999 Jan 1;16(1):37–48.

21. Andolfo I, Russo R, Lasorsa VA, Cantalupo S, Rosato BE, Bonfiglio F, et al. Common variants at 21q22.3 locus influence MX1 and TMPRSS2 gene expression and susceptibility to severe COVID-19. iScience. 2021 Apr 23;24(4):102322.

22. Asselta R, Paraboschi EM, Mantovani A, Duga S. ACE2 and TMPRSS2 variants and expression as candidates to sex and country differences in COVID-19 severity in Italy. Aging. 2020 Jun 5;12(11):10087–98.

23. Bhattacharyya C, Das C, Ghosh A, Singh AK, Mukherjee S, Majumder PP, et al. Global Spread of SARS-CoV-2 Subtype with Spike Protein Mutation D614G is Shaped by Human Genomic Variations that Regulate Expression of TMPRSS2 and MX1 Genes [Internet]. 2020 May [cited 2021 Aug 28] p. 2020.05.04.075911. Available from: https://www.biorxiv.org/content/10.1101/2020.05.04.075911v1

24. Darbani B. The Expression and Polymorphism of Entry Machinery for COVID-19 in Human: Juxtaposing Population Groups, Gender, and Different Tissues. Int J Environ Res Public Health. 2020 Jan;17(10):3433.

25. Hou Y, Zhao J, Martin W, Kallianpur A, Chung MK, Jehi L, et al. New insights into genetic susceptibility of COVID-19: an ACE2 and TMPRSS2 polymorphism analysis. BMC Med. 2020 Jul 15;18(1):216.

26. Irham LM, Chou W-H, Calkins MJ, Adikusuma W, Hsieh S-L, Chang W-C. Genetic variants that influence SARS-CoV-2 receptor TMPRSS2 expression among population cohorts from multiple continents. Biochem Biophys Res Commun. 2020 Aug 20;529(2):263–9.

27. Iyer GR, Samajder S, Zubeda S, S DSN, Mali V, PV SK, et al. Infectivity and Progression of COVID-19 Based on Selected Host Candidate Gene Variants. Front Genet. 2020;11:861.

28. Jeon S, Blazyte A, Yoon C, Ryu H, Jeon Y, Bhak Y, et al. Ethnicity-dependent allele frequencies are correlated with COVID-19 case fatality rate [Internet]. Preprints; 2020 Oct [cited 2021 Aug 28]. Available from: https://www.authorea.com/users/367817/articles/487091-ethnicity-dependent-allele-frequencies-are-correlated-with-covid-19-case-fatality-rate?commit=92f9ba974af4c5e0ff312d7dd9994aa1b1589975

29. Kim Y-C, Jeong B-H. Strong Correlation between the Case Fatality Rate of COVID-19 and the rs6598045 Single Nucleotide Polymorphism (SNP) of the Interferon-Induced Transmembrane Protein 3 (IFITM3) Gene at the Population-Level. Genes. 2021 Jan;12(1):42.

30. Latini A, Agolini E, Novelli A, Borgiani P, Giannini R, Gravina P, et al. COVID-19 and Genetic Variants of Protein Involved in the SARS-CoV-2 Entry into the Host Cells. Genes. 2020 Sep;11(9):1010.

31. Paniri A, Hosseini MM, Akhavan-Niaki H. First comprehensive computational analysis of functional consequences of TMPRSS2 SNPs in susceptibility to SARS-CoV-2 among different populations. J Biomol Struct Dyn. 2021 Jul 3;39(10):3576–93.

32. Piva F, Sabanovic B, Cecati M, Giulietti M. Expression and co-expression analyses of TMPRSS2, a key element in COVID-19. Eur J Clin Microbiol Infect Dis. 2021 Feb 1;40(2):451–5.

33. Ragia G, Manolopoulos VG. Assessing COVID-19 susceptibility through analysis of the genetic and epigenetic diversity of ACE2-mediated SARS-CoV-2 entry. Pharmacogenomics. 2020 Dec 1;21(18):1311–29.

34. Senapati S, Kumar S, Singh AK, Banerjee P, Bhagavatula S. Assessment of risk conferred by coding and regulatory variations of TMPRSS2 and CD26 in susceptibility to SARS-CoV-2 infection in human. J Genet. 2020;99:53.

35. Sharma S, Singh I, Haider S, Malik MZ, Ponnusamy K, Rai E. ACE2 Homo-dimerization, Human Genomic variants and Interaction of Host Proteins Explain High Population Specific Differences in Outcomes of COVID19 [Internet]. 2020 Apr [cited 2021 Aug 28] p. 2020.04.24.050534. Available from: https://www.biorxiv.org/content/10.1101/2020.04.24.050534v1

36. Singh H, Choudhari R, Nema V, Khan AA. ACE2 and TMPRSS2 polymorphisms in various diseases with special reference to its impact on COVID-19 disease. Microb Pathog. 2021 Jan 1;150:104621.

37. Strope JD, PharmD CHC, Figg WD. TMPRSS2: Potential Biomarker for COVID-19 Outcomes. J Clin Pharmacol. 2020 May 21;10.1002/jcph.1641.

38. Torre-Fuentes L, Matías-Guiu J, Hernández-Lorenzo L, Montero-Escribano P, Pytel V, Porta-Etessam J, et al. ACE2, TMPRSS2, and Furin variants and SARS-CoV-2 infection in Madrid, Spain. J Med Virol. 2021 Feb;93(2):863–9.

39. Vargas-Alarcón G, Posadas-Sánchez R, Ramírez-Bello J. Variability in genes related to SARS-CoV-2 entry into host cells (ACE2, TMPRSS2, TMPRSS11A, ELANE, and CTSL) and its potential use in association studies. Life Sci. 2020 Nov 1;260:118313.

40. Wang F, Huang S, Gao R, Zhou Y, Lai C, Li Z, et al. Initial whole-genome sequencing and analysis of the host genetic contribution to COVID-19 severity and susceptibility. Cell Discov. 2020 Nov 10;6(1):1–16.

41. Wulandari L, Hamidah B, Pakpahan C, Damayanti NS, Kurniati ND, Adiatmaja CO, et al. Initial study on TMPRSS2 p.Val160Met genetic variant in COVID-19 patients. Hum Genomics. 2021 May 17;15(1):29.

42. Chaubey G, Ayub Q, Rai N, Prakash S, Mushrif-Tripathy V, Mezzavilla M, et al. “Like sugar in milk”: reconstructing the genetic history of the Parsi population. Genome Biol. 2017 Jun 14;18(1):110.

43. Pathak AK, Kadian A, Kushniarevich A, Montinaro F, Mondal M, Ongaro L, et al. The Genetic Ancestry of Modern Indus Valley Populations from Northwest India. Am J Hum Genet. 2018 Dec 6;103(6):918–29.

44. Re3data.Org. Estonian Biocentre Public Data. 2014 [cited 2021 Aug 28]; Available from: http://service.re3data.org/repository/r3d100010986

45. Tätte K, Pagani L, Pathak AK, Kõks S, Ho Duy B, Ho XD, et al. The genetic legacy of continental scale admixture in Indian Austroasiatic speakers. Sci Rep. 2019 Mar 7;9:3818.

46. Durbin RM, Altshuler D, Durbin RM, Abecasis GR, Bentley DR, Chakravarti A, et al. A map of human genome variation from population-scale sequencing. Nature. 2010 Oct;467(7319):1061–73.

47. Zhang C, Gao Y, Ning Z, Lu Y, Zhang X, Liu J, et al. PGG.SNV: understanding the evolutionary and medical implications of human single nucleotide variations in diverse populations. Genome Biol. 2019 Oct 22;20(1):215.

48. Benesty J, Chen J, Huang Y, Cohen I. Pearson Correlation Coefficient. In: Cohen I, Huang Y, Chen J, Benesty J, editors. Noise Reduction in Speech Processing [Internet]. Berlin, Heidelberg: Springer; 2009 [cited 2021 Aug 28]. p. 1–4. (Springer Topics in Signal Processing). Available from: https://doi.org/10.1007/978-3-642-00296-0_5

49. Barrett JC, Fry B, Maller J, Daly MJ. Haploview: analysis and visualization of LD and haplotype maps. Bioinformatics. 2005 Jan 15;21(2):263–5.

50. Paoloni-Giacobino A, Chen H, Peitsch MC, Rossier C, Antonarakis SE. Cloning of the TMPRSS2 Gene, Which Encodes a Novel Serine Protease with Transmembrane, LDLRA, and SRCR Domains and Maps to 21q22.3. Genomics. 1997 Sep 15;44(3):309–20.

51. Huggins DJ. Structural analysis of experimental drugs binding to the SARS-CoV-2 target TMPRSS2. J Mol Graph Model. 2020 Nov 1;100:107710.

52. Cheng Z, Zhou J, To KK-W, Chu H, Li C, Wang D, et al. Identification of TMPRSS2 as a Susceptibility Gene for Severe 2009 Pandemic A(H1N1) Influenza and A(H7N9) Influenza. J Infect Dis. 2015 Oct 15;212(8):1214–21.

53. Alshahawey M, Raslan M, Sabri N. Sex-mediated effects of ACE2 and TMPRSS2 on the incidence and severity of COVID-19; The need for genetic implementation. Curr Res Transl Med. 2020 Nov;68(4):149–50.

54. Lamy P-J, Rébillard X, Vacherot F, de la Taille A. Androgenic hormones and the excess male mortality observed in COVID-19 patients: new convergent data. World J Urol. 2020 Jun 2;1–3.

55. Peckham H, de Gruijter NM, Raine C, Radziszewska A, Ciurtin C, Wedderburn LR, et al. Male sex identified by global COVID-19 meta-analysis as a risk factor for death and ITU admission. Nat Commun. 2020 Dec 9;11(1):6317.

56. Souilmi Y, Lauterbur ME, Tobler R, Huber CD, Johar AS, Moradi SV, et al. An ancient viral epidemic involving host coronavirus interacting genes more than 20,000 years ago in East Asia. Curr Biol. 2021 Aug 23;31(16):3504–3514.e9.

