## Supplementary Figures and Tables for "Genetic association of *TMPRSS2* rs2070788 polymorphism with COVID-19 Case Fatality Rate among Indian populations"

### Supplementary Figure 1

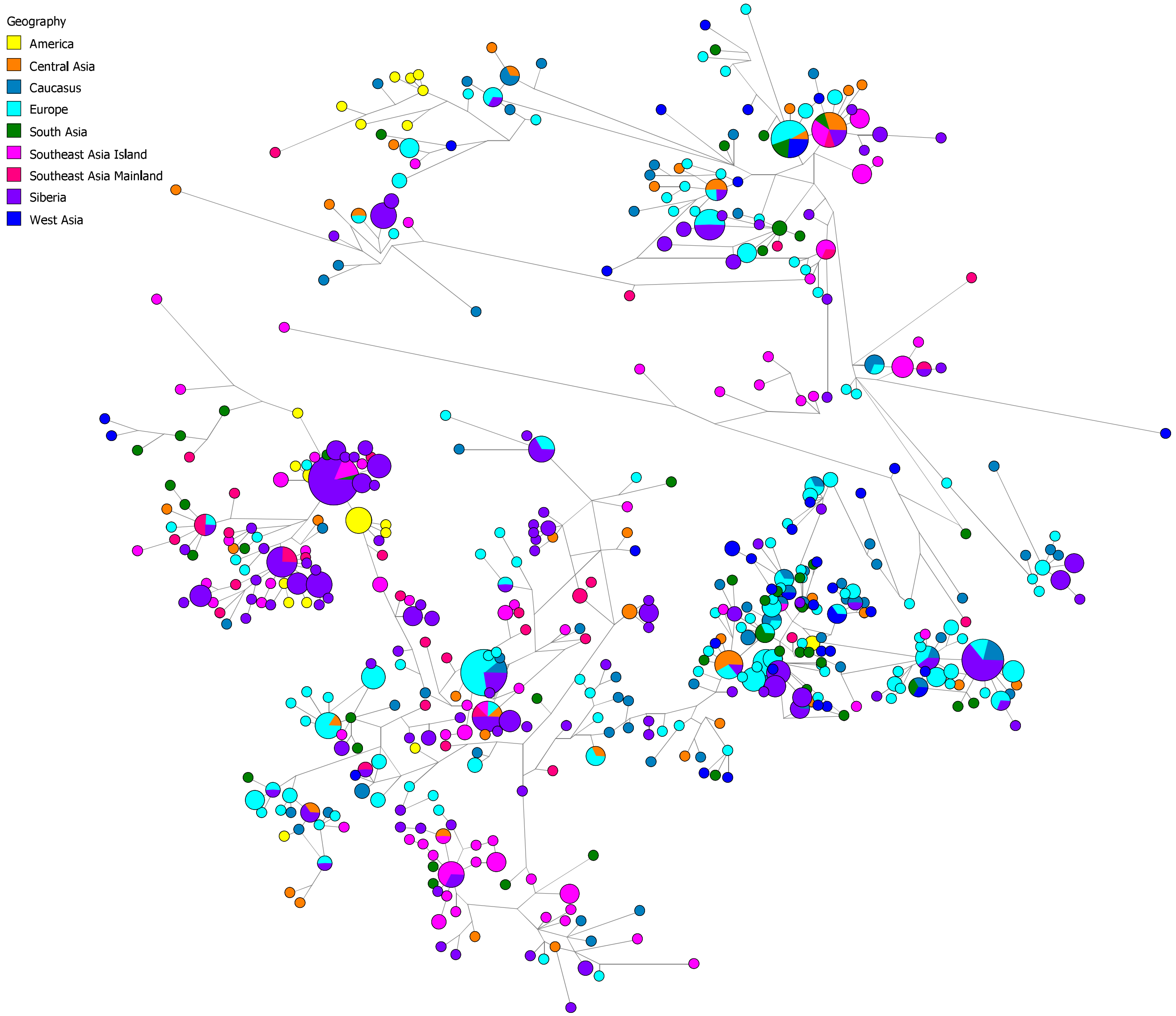

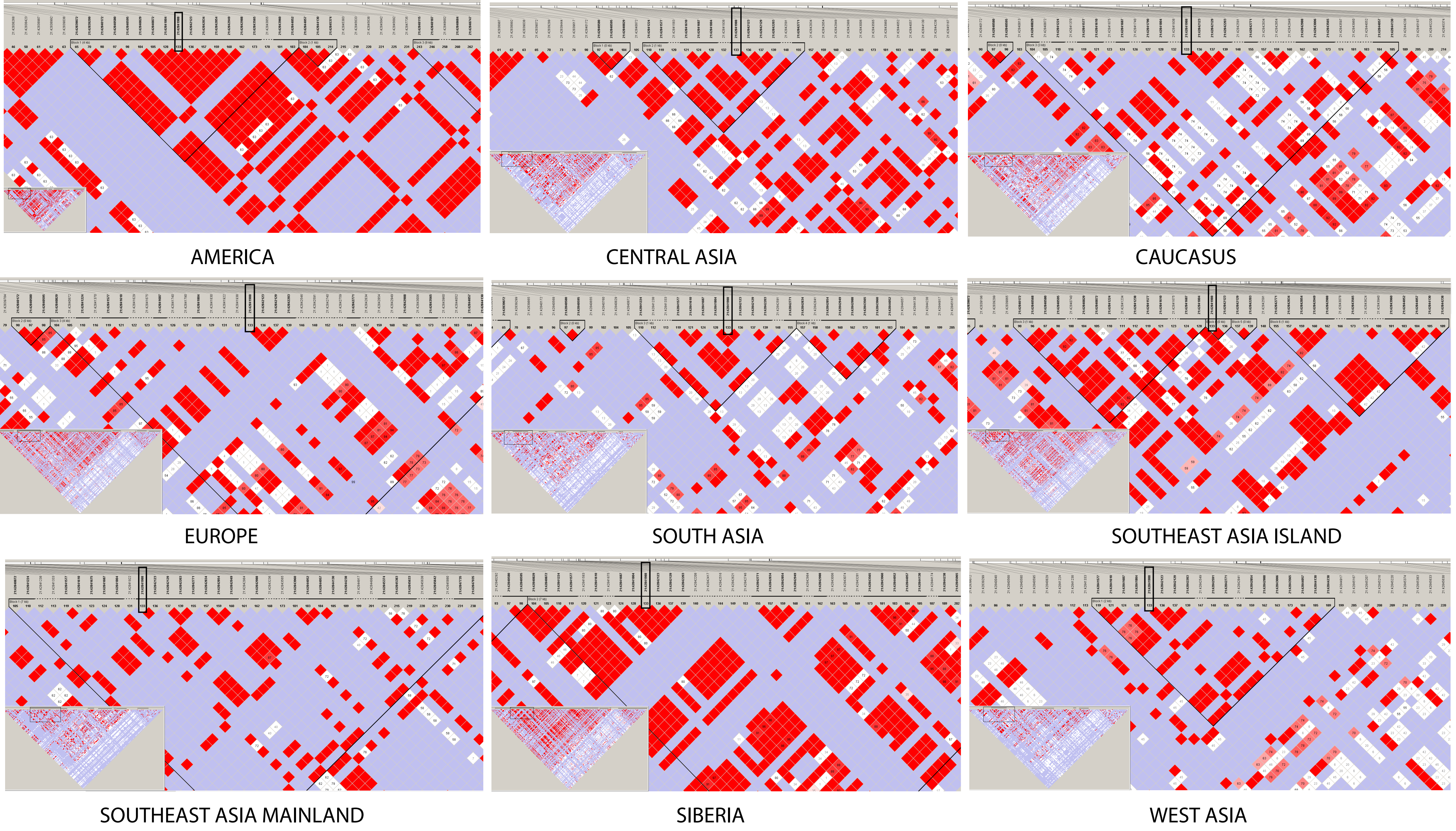

Supplementary Figure 2

Supplementary Figure 3

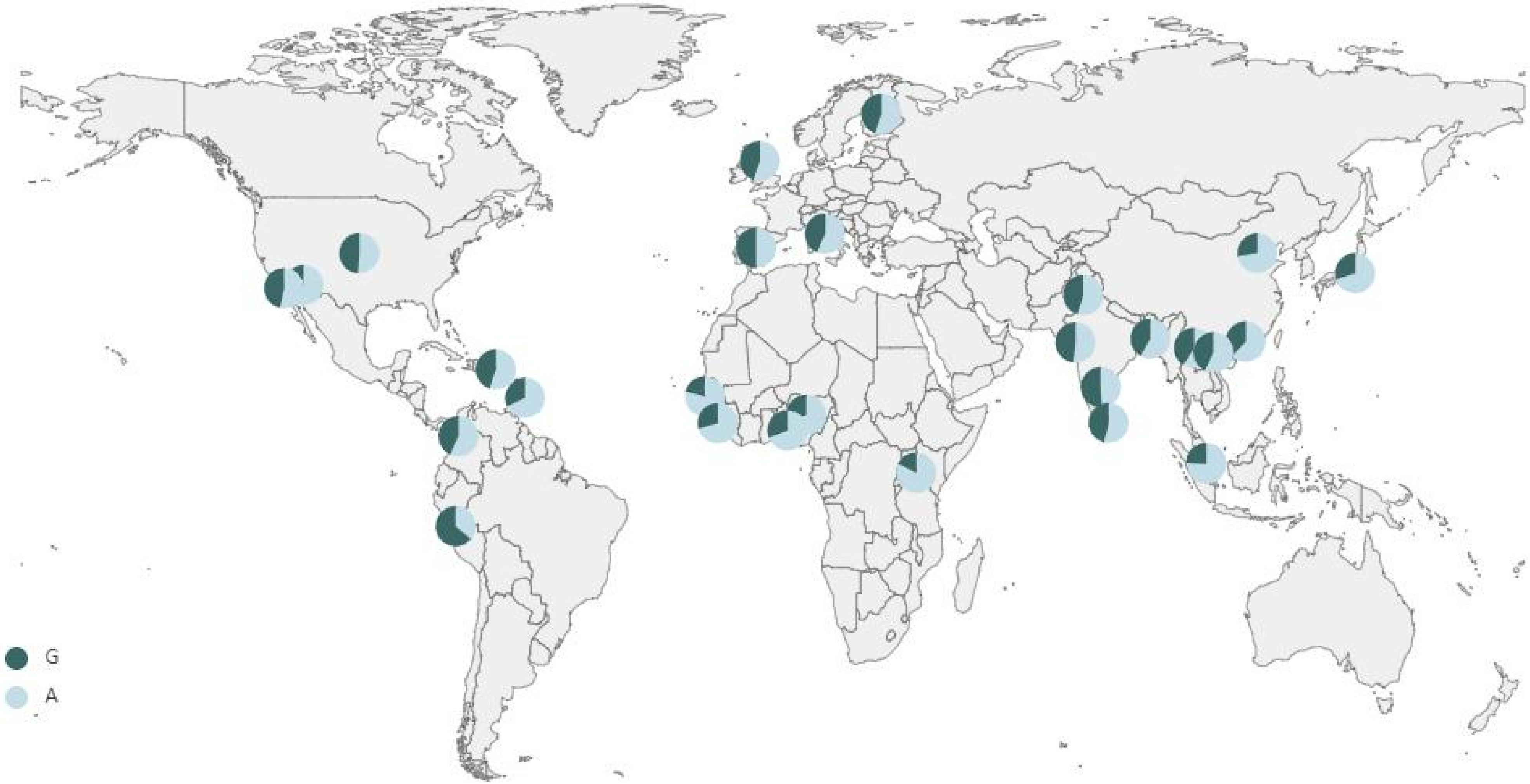

**Supplementary Table 1**. The details of SNPs observed in TMPRSS2 gene. MAF- Minor Allele Frequency, AMR-America, CA-Central Asia, CAU-Caucasus, EU-Europe, SA-South Asia, SEAI-Southeast Asia Island, SEAM-Southeast Asia Mainland SIB-Siberia, WA-West Asia

| S. No. | rs number | Build 37 | Build 38 | Ref Allele | Alternate Allele | Functional Consequence | Total MAF | AMR MAF | CA MAF | CAU MAF | EU MAF | SA MAF | SEAI MAF | SEAM MAF | SIB MAF | WA MAF |
|---|---|---|---|---|---|---|---|---|---|---|---|---|---|---|---|---|
| 1 | rs547758146 | 42836222 | 41464295 | A | C | downstream_gene_variant | 0 | 0 | 0 | 0 | 0.0198 | 0 | 0 | 0 | 0 | 0 |
| 2 | rs1308901935 | 42836290 | 41464363 | C | T | downstream_gene_variant | 0 | 0 | 0 | 0 | 0 | 0 | 0 | 0 | 0 | 0 |
| 3 | rs559811756 | 42836341 | 41464414 | A | G | downstream_gene_variant | 0 | 0 | 0 | 0 | 0 | 0 | 0 | 0 | 0 | 0 |
| 4 | rs8129769 | 42836344 | 41464417 | A | G | downstream_gene_variant | 0 | 0 | 0 | 0 | 0 | 0 | 0 | 0 | 0 | 0 |
| 5 | rs553400003 | 42836389 | 41464462 | A | G | downstream_gene_variant | 0 | 0 | 0 | 0 | 0 | 0 | 0 | 0 | 0 | 0 |
| 6 | rs768729764 | 42836417 | 41464490 | T | C | downstream_gene_variant | 0 | 0 | 0 | 0 | 0 | 0 | 0 | 0 | 0 | 0 |
| 7 | - | 42836476 | 41464549 | A | G | downstream_gene_variant | 0.3462 | 0.3462 | 0.2292 | 0.2179 | 0.2277 | 0.32 | 0.3444 | 0.4167 | 0.4583 | 0.05 |
| 8 | rs76000363 | 42836477 | 41464550 | A | G | downstream_gene_variant | 0.07692 | 0.07692 | 0.08333 | 0.07692 | 0.1683 | 0.06 | 0.02222 | 0.08333 | 0.06019 | 0.025 |
| 9 | - | 42836485 | 41464558 | A | G | 3_prime_UTR_variant | 0 | 0 | 0 | 0 | 0 | 0 | 0 | 0 | 0.00463 | 0 |
| 10 | - | 42836496 | 41464569 | T | C | 3_prime_UTR_variant | 0.3462 | 0.3462 | 0.2292 | 0.2179 | 0.2277 | 0.32 | 0.3333 | 0.4167 | 0.463 | 0.075 |
| 11 | rs929663872 | 42836594 | 41464667 | C | G | 3_prime_UTR_variant | 0 | 0 | 0 | 0 | 0 | 0 | 0 | 0 | 0 | 0 |
| 12 | rs185078457 | 42836613 | 41464686 | T | C | 3_prime_UTR_variant | 0 | 0 | 0.02083 | 0 | 0.009901 | 0 | 0 | 0 | 0 | 0 |
| 13 | - | 42836664 | 41464737 | C | T | 3_prime_UTR_variant | 0.03846 | 0.03846 | 0 | 0 | 0 | 0 | 0 | 0 | 0 | 0 |
| 14 | rs112657409 | 42836672 | 41464745 | T | C | 3_prime_UTR_variant | 0 | 0 | 0 | 0.01282 | 0.00495 | 0 | 0.04444 | 0.08333 | 0.03241 | 0 |
| 15 | rs2838038 | 42836685 | 41464758 | T | A | 3_prime_UTR_variant | 0.07692 | 0.07692 | 0.08333 | 0.07692 | 0.1683 | 0.06 | 0.02222 | 0.08333 | 0.06019 | 0.025 |
| 16 | rs377038375 | 42836701 | 41464774 | C | T | 3_prime_UTR_variant | 0 | 0 | 0 | 0 | 0 | 0 | 0.06667 | 0 | 0 | 0 |
| 17 | - | 42836729 | 41464802 | A | G | 3_prime_UTR_variant | 0.4231 | 0.4231 | 0.1458 | 0.141 | 0.05941 | 0.24 | 0.4444 | 0.4722 | 0.3981 | 0.025 |
| 18 | - | 42836751 | 41464824 | T | A | 3_prime_UTR_variant | 0.3462 | 0.3462 | 0.2292 | 0.2179 | 0.2277 | 0.32 | 0.3444 | 0.4167 | 0.4583 | 0.05 |
| 19 | - | 42836873 | 41464946 | G | A | 3_prime_UTR_variant | 0 | 0 | 0 | 0 | 0 | 0 | 0 | 0 | 0 | 0 |
| 20 | rs117652812 | 42836891 | 41464964 | T | C | 3_prime_UTR_variant | 0 | 0 | 0.04167 | 0.01282 | 0.009901 | 0 | 0.05556 | 0 | 0.02315 | 0 |
| 21 | - | 42836921 | 41464994 | A | G | 3_prime_UTR_variant | 0 | 0 | 0 | 0 | 0.00495 | 0 | 0 | 0 | 0 | 0 |
| 22 | rs17001042 | 42836967 | 41465040 | A | G | 3_prime_UTR_variant | 0 | 0 | 0 | 0 | 0 | 0.02 | 0.07778 | 0.02778 | 0 | 0 |
| 23 | rs572422499 | 42836971 | 41465044 | T | C | 3_prime_UTR_variant | 0 | 0 | 0 | 0.01282 | 0 | 0 | 0 | 0 | 0 | 0 |
| 24 | rs11910678 | 42836986 | 41465059 | C | T | 3_prime_UTR_variant | 0 | 0 | 0 | 0.01282 | 0.00495 | 0 | 0.05556 | 0.08333 | 0.03241 | 0 |
| 25 | rs183418223 | 42837069 | 41465142 | A | G | 3_prime_UTR_variant | 0 | 0 | 0 | 0 | 0 | 0 | 0 | 0 | 0 | 0 |
| 26 | rs1453389785,COSV100152071 | 42837277 | 41465350 | T | C | 3_prime_UTR_variant | 0 | 0 | 0 | 0 | 0 | 0.02 | 0 | 0 | 0 | 0 |
| 27 | rs377348479 | 42837392 | 41465465 | A | G | 3_prime_UTR_variant | 0 | 0 | 0 | 0 | 0 | 0 | 0.01111 | 0.02778 | 0 | 0 |
| 28 | rs75317729 | 42837502 | 41465575 | C | T | 3_prime_UTR_variant | 0 | 0 | 0 | 0 | 0 | 0 | 0 | 0 | 0 | 0 |
| 29 | rs771633249 | 42837518 | 41465591 | T | C | 3_prime_UTR_variant | 0 | 0 | 0 | 0 | 0 | 0 | 0 | 0 | 0 | 0 |
| 30 | rs370421631 | 42837531 | 41465604 | A | G | 3_prime_UTR_variant | 0 | 0 | 0 | 0 | 0 | 0 | 0.01111 | 0 | 0 | 0 |
| 31 | rs77675406 | 42837534 | 41465607 | A | G | 3_prime_UTR_variant | 0.07692 | 0.07692 | 0.08333 | 0.07692 | 0.1683 | 0.06 | 0.02222 | 0.08333 | 0.06019 | 0.025 |
| 32 | rs542458473 | 42837636 | 41465709 | C | G | 3_prime_UTR_variant | 0 | 0 | 0 | 0 | 0 | 0.04 | 0 | 0 | 0 | 0 |
| 33 | rs12627374 | 42837691 | 41465764 | T | C | 3_prime_UTR_variant | 0 | 0 | 0.0625 | 0.05128 | 0.01485 | 0.12 | 0.3 | 0.05556 | 0.05556 | 0 |
| 34 | rs567103273 | 42837751 | 41465824 | G | A | 3_prime_UTR_variant | 0 | 0 | 0 | 0 | 0 | 0 | 0 | 0 | 0 | 0 |
| 35 | rs372405355 | 42837754 | 41465827 | C | T | 3_prime_UTR_variant | 0 | 0 | 0 | 0 | 0 | 0 | 0.06667 | 0 | 0 | 0 |
| 36 | rs996615575 | 42837756 | 41465829 | C | G | 3_prime_UTR_variant | 0 | 0 | 0 | 0 | 0 | 0 | 0.01111 | 0 | 0 | 0 |
| 37 | rs62217525 | 42837848 | 41465921 | T | C | 3_prime_UTR_variant | 0 | 0 | 0.02083 | 0.07692 | 0.04455 | 0 | 0 | 0 | 0.00463 | 0.2 |
| 38 | rs146564124 | 42837861 | 41465934 | T | C | 3_prime_UTR_variant | 0 | 0 | 0 | 0.01282 | 0.0198 | 0 | 0 | 0 | 0.00463 | 0 |
| 39 | rs780868244 | 42837913 | 41465986 | G | A | 3_prime_UTR_variant | 0 | 0 | 0 | 0 | 0 | 0 | 0.02222 | 0 | 0 | 0 |
| 40 | rs77996454 | 42837928 | 41466001 | A | G | 3_prime_UTR_variant | 0 | 0 | 0 | 0 | 0 | 0 | 0 | 0 | 0 | 0 |
| 41 | rs374981614 | 42838036 | 41466109 | A | G | 3_prime_UTR_variant | 0 | 0 | 0 | 0 | 0 | 0 | 0.06667 | 0 | 0 | 0 |
| 42 | rs118028230 | 42838104 | 41466177 | C | G | intron_variant | 0 | 0 | 0 | 0.01282 | 0.00495 | 0 | 0.04444 | 0.08333 | 0.03241 | 0 |
| 43 | rs369092228 | 42838114 | 41466187 | C | T | intron_variant | 0 | 0 | 0 | 0 | 0 | 0 | 0 | 0 | 0 | 0 |
| 44 | rs73905370,COSV59821189 | 42838138 | 41466211 | T | A | intron_variant | 0.07692 | 0.07692 | 0.08333 | 0.07692 | 0.1683 | 0.06 | 0.02222 | 0.08333 | 0.06019 | 0.025 |
| 45 | rs55896064 | 42838198 | 41466271 | A | G | intron_variant | 0.07692 | 0.07692 | 0.08333 | 0.07692 | 0.1683 | 0.06 | 0.02222 | 0.08333 | 0.06019 | 0.025 |
| 46 | rs57474639 | 42838268 | 41466341 | T | C | intron_variant | 0.07692 | 0.07692 | 0.08333 | 0.07692 | 0.1683 | 0.06 | 0.02222 | 0.08333 | 0.06019 | 0.025 |
| 47 | rs79971314 | 42838269 | 41466342 | A | G | intron_variant | 0 | 0 | 0 | 0 | 0 | 0.02 | 0.06667 | 0.02778 | 0 | 0.05 |
| 48 | rs73372161 | 42838279 | 41466352 | A | G | intron_variant | 0 | 0 | 0 | 0 | 0 | 0.02 | 0 | 0.02778 | 0 | 0.05 |
| 49 | rs370634518 | 42838411 | 41466484 | A | C | intron_variant | 0 | 0 | 0 | 0 | 0 | 0 | 0 | 0 | 0 | 0 |
| 50 | rs61459778,COSV59822080 | 42838423 | 41466496 | G | C | intron_variant | 0.07692 | 0.07692 | 0.08333 | 0.07692 | 0.1683 | 0.08 | 0.1 | 0.1111 | 0.06019 | 0.075 |
| 51 | rs145024812 | 42838456 | 41466529 | A | G | intron_variant | 0 | 0 | 0 | 0 | 0 | 0 | 0 | 0 | 0 | 0 |
| 52 | rs377591443 | 42838514 | 41466587 | C | G | intron_variant | 0 | 0 | 0 | 0 | 0 | 0 | 0.01111 | 0 | 0 | 0 |

| | | | | | | | | | | | | | | | | |
|---|---|---|---|---|---|---|---|---|---|---|---|---|---|---|---|---|
| 53 | rs115975538 | 42838629 | 41466702 | A | C | intron_variant | 0 | 0 | 0 | 0 | 0 | 0 | 0 | 0 | 0 | 0 |
| 54 | rs118133613 | 42838665 | 41466738 | A | G | intron_variant | 0 | 0 | 0 | 0 | 0.00495 | 0 | 0 | 0 | 0 | 0 |
| 55 | rs143595083 | 42838707 | 41466780 | T | C | intron_variant | 0 | 0 | 0 | 0 | 0 | 0 | 0 | 0 | 0 | 0 |
| 56 | rs148038688 | 42838711 | 41466784 | T | C | intron_variant | 0 | 0 | 0 | 0 | 0 | 0 | 0 | 0 | 0 | 0 |
| 57 | rs546913068 | 42838803 | 41466876 | G | C | intron_variant | 0 | 0 | 0 | 0 | 0 | 0 | 0 | 0 | 0.009259 | 0 |
| 58 | - | 42838860 | 41466933 | T | A | intron_variant | 0 | 0 | 0 | 0 | 0 | 0 | 0 | 0.02778 | 0 | 0 |
| 59 | rs534326511 | 42838913 | 41466986 | T | C | intron_variant | 0 | 0 | 0 | 0 | 0 | 0 | 0 | 0 | 0.009259 | 0 |
| 60 | rs1033026879 | 42838937 | 41467010 | A | C | intron_variant | 0 | 0 | 0 | 0 | 0 | 0 | 0.01111 | 0 | 0 | 0 |
| 61 | rs73905371 | 42838987 | 41467060 | G | C | intron_variant | 0.07692 | 0.07692 | 0.08333 | 0.07692 | 0.1683 | 0.06 | 0.02222 | 0.08333 | 0.06019 | 0.025 |
| 62 | rs73372163 | 42838992 | 41467065 | A | G | intron_variant | 0.07692 | 0.07692 | 0.08333 | 0.07692 | 0.1683 | 0.08 | 0.1 | 0.1111 | 0.06019 | 0.075 |
| 63 | rs73372166 | 42839038 | 41467111 | A | G | intron_variant | 0.07692 | 0.07692 | 0.08333 | 0.07692 | 0.1683 | 0.08 | 0.1 | 0.1111 | 0.06019 | 0.075 |
| 64 | rs113562865 | 42839044 | 41467117 | T | C | intron_variant | 0 | 0 | 0 | 0 | 0 | 0.02 | 0 | 0.02778 | 0 | 0.05 |
| 65 | - | 42839072 | 41467145 | G | T | intron_variant | 0.4231 | 0.4231 | 0.1458 | 0.141 | 0.05941 | 0.24 | 0.4556 | 0.4722 | 0.4028 | 0.025 |
| 66 | - | 42839196 | 41467269 | G | A | intron_variant | 0 | 0 | 0 | 0 | 0 | 0 | 0.03333 | 0 | 0 | 0 |
| 67 | rs368452629 | 42839233 | 41467306 | C | T | intron_variant | 0 | 0 | 0 | 0 | 0 | 0 | 0 | 0 | 0 | 0 |
| 68 | rs981602772 | 42839261 | 41467334 | T | A | intron_variant | 0 | 0 | 0 | 0 | 0.00495 | 0 | 0 | 0 | 0 | 0 |
| 69 | rs371583992 | 42839278 | 41467351 | G | T | intron_variant | 0 | 0 | 0 | 0 | 0 | 0 | 0 | 0 | 0 | 0 |
| 70 | - | 42839299 | 41467372 | C | G | intron_variant | 0.4231 | 0.4231 | 0.08333 | 0.07692 | 0.0396 | 0.12 | 0.2 | 0.3333 | 0.3102 | 0.025 |
| 71 | - | 42839361 | 41467434 | G | A | intron_variant | 0 | 0 | 0 | 0 | 0 | 0 | 0 | 0 | 0 | 0 |
| 72 | rs73372168 | 42839362 | 41467435 | A | G | intron_variant | 0 | 0 | 0 | 0 | 0 | 0 | 0 | 0 | 0 | 0 |
| 73 | rs910388295 | 42839444 | 41467517 | G | C | intron_variant | 0 | 0 | 0.02083 | 0 | 0.009901 | 0 | 0 | 0 | 0.00463 | 0 |
| 74 | rs192955773 | 42839545 | 41467618 | C | T | intron_variant | 0 | 0 | 0 | 0 | 0 | 0 | 0 | 0 | 0 | 0 |
| 75 | rs961773771 | 42839588 | 41467661 | G | C | intron_variant | 0 | 0 | 0 | 0 | 0 | 0 | 0 | 0 | 0 | 0 |
| 76 | rs1429823013 | 42839617 | 41467690 | G | C | intron_variant | 0 | 0 | 0.02083 | 0 | 0 | 0 | 0 | 0 | 0 | 0 |
| 77 | rs183984610 | 42839626 | 41467699 | A | G | intron_variant | 0 | 0 | 0 | 0 | 0 | 0 | 0 | 0 | 0 | 0 |
| 78 | rs142750000 | 42839661 | 41467734 | T | C | splice_region_variant,synonymous_varian | 0 | 0 | 0 | 0 | 0.00495 | 0 | 0 | 0 | 0 | 0 |
| 79 | rs374377315 | 42839784 | 41467857 | T | C | synonymous_variant | 0 | 0 | 0 | 0 | 0.00495 | 0 | 0 | 0 | 0 | 0 |
| 80 | rs147359020 | 42839865 | 41467938 | A | G | intron_variant | 0 | 0 | 0 | 0 | 0 | 0.04 | 0.01111 | 0 | 0.00463 | 0 |
| 81 | rs1016428148 | 42839900 | 41467973 | G | C | intron_variant | 0 | 0 | 0 | 0 | 0 | 0 | 0 | 0 | 0 | 0 |
| 82 | rs139015396 | 42839982 | 41468055 | T | C | intron_variant | 0 | 0 | 0 | 0 | 0 | 0 | 0 | 0 | 0 | 0 |
| 83 | rs149855493 | 42839993 | 41468066 | C | T | intron_variant | 0 | 0 | 0 | 0 | 0 | 0 | 0 | 0 | 0 | 0 |
| 84 | rs1014364281 | 42839999 | 41468072 | G | C | intron_variant | 0 | 0 | 0 | 0 | 0 | 0 | 0 | 0 | 0 | 0 |
| 85 | rs144823388 | 42840029 | 41468102 | T | A | intron_variant | 0 | 0 | 0 | 0 | 0 | 0 | 0 | 0 | 0 | 0 |
| 86 | rs191394761 | 42840048 | 41468121 | T | C | intron_variant | 0 | 0 | 0 | 0 | 0 | 0 | 0 | 0 | 0 | 0.05 |
| 87 | rs76315847 | 42840130 | 41468203 | A | G | intron_variant | 0 | 0 | 0 | 0 | 0 | 0 | 0 | 0 | 0 | 0 |
| 88 | rs146605032 | 42840168 | 41468241 | A | G | intron_variant | 0 | 0 | 0 | 0 | 0 | 0 | 0 | 0 | 0 | 0 |
| 89 | rs1017698498 | 42840171 | 41468244 | T | C | intron_variant | 0 | 0 | 0 | 0.01282 | 0 | 0 | 0 | 0 | 0 | 0 |
| 90 | rs743542 | 42840172 | 41468245 | A | G | intron_variant | 0.2692 | 0.2692 | 0.2083 | 0.1154 | 0.1287 | 0.18 | 0.3889 | 0.25 | 0.213 | 0 |
| 91 | rs190618812 | 42840187 | 41468260 | A | G | intron_variant | 0 | 0 | 0 | 0 | 0 | 0 | 0 | 0.02778 | 0.00463 | 0 |
| 92 | - | 42840267 | 41468340 | A | G | intron_variant | 0 | 0 | 0 | 0 | 0 | 0 | 0 | 0 | 0.00463 | 0 |
| 93 | rs572963642,COSV59821705 | 42840292 | 41468365 | C | G | intron_variant | 0 | 0 | 0 | 0 | 0 | 0 | 0 | 0 | 0.009259 | 0 |
| 94 | rs148125094 | 42840394 | 41468467 | T | C | missense_variant | 0 | 0 | 0 | 0 | 0 | 0 | 0 | 0 | 0 | 0 |
| 95 | rs375874663 | 42840504 | 41468577 | C | T | intron_variant | 0 | 0 | 0 | 0 | 0 | 0 | 0 | 0 | 0 | 0 |
| 96 | rs73372170 | 42840568 | 41468641 | G | C | intron_variant | 0 | 0 | 0 | 0 | 0 | 0.02 | 0.06667 | 0.02778 | 0 | 0.05 |
| 97 | COSV59827297 | 42840580 | 41468653 | T | C | intron_variant | 0.07692 | 0.07692 | 0.3542 | 0.2308 | 0.3218 | 0.34 | 0.3444 | 0.25 | 0.3843 | 0.025 |
| 98 | - | 42840595 | 41468668 | G | C | intron_variant | 0.07692 | 0.07692 | 0.3542 | 0.2308 | 0.3218 | 0.34 | 0.3444 | 0.25 | 0.3843 | 0.025 |
| 99 | rs143672898 | 42840655 | 41468728 | T | C | intron_variant | 0 | 0 | 0 | 0 | 0 | 0.02 | 0 | 0 | 0 | 0 |
| 100 | rs748580091 | 42840740 | 41468813 | T | A | intron_variant | 0 | 0 | 0 | 0 | 0 | 0 | 0.01111 | 0 | 0 | 0 |
| 101 | rs146797606 | 42840758 | 41468831 | C | T | intron_variant | 0 | 0 | 0 | 0 | 0 | 0 | 0 | 0 | 0 | 0 |
| 102 | rs573343651 | 42840760 | 41468833 | C | A | intron_variant | 0 | 0 | 0 | 0 | 0 | 0.02 | 0 | 0.02778 | 0 | 0 |
| 103 | rs1443771075 | 42840813 | 41468886 | T | C | intron_variant | 0 | 0 | 0 | 0.01282 | 0 | 0 | 0 | 0 | 0 | 0 |
| 104 | rs7283324 | 42840829 | 41468902 | T | C | intron_variant | 0.3462 | 0.3462 | 0.3125 | 0.2179 | 0.3267 | 0.24 | 0.4667 | 0.4167 | 0.3056 | 0.075 |
| 105 | - | 42840872 | 41468945 | T | C | intron_variant | 0.4231 | 0.4231 | 0.0625 | 0.02564 | 0.0297 | 0.12 | 0.1667 | 0.3333 | 0.3102 | 0 |

| | | | | | | | | | | | | | | | | |
|---|---|---|---|---|---|---|---|---|---|---|---|---|---|---|---|---|
| 106 | rs150382508 | 42841054 | 41469127 | C | A | intron_variant | 0 | 0 | 0 | 0 | 0 | 0 | 0 | 0 | 0 | 0 |
| 107 | rs115774576 | 42841073 | 41469146 | A | T | intron_variant | 0 | 0 | 0 | 0 | 0 | 0 | 0 | 0 | 0 | 0 |
| 108 | rs114549926 | 42841128 | 41469201 | A | G | intron_variant | 0 | 0 | 0 | 0 | 0 | 0 | 0 | 0 | 0 | 0 |
| 109 | rs376403654 | 42841168 | 41469241 | A | G | intron_variant | 0 | 0 | 0 | 0 | 0 | 0 | 0 | 0 | 0 | 0 |
| 110 | rs7279603,COSV59826519 | 42841224 | 41469297 | C | T | intron_variant | 0 | 0 | 0.3333 | 0.3077 | 0.2475 | 0.22 | 0.2889 | 0.2222 | 0.1991 | 0.375 |
| 111 | rs971603670 | 42841234 | 41469307 | C | G | intron_variant | 0 | 0 | 0 | 0 | 0 | 0 | 0.03333 | 0 | 0 | 0 |
| 112 | rs7278739 | 42841238 | 41469311 | A | G | intron_variant | 0 | 0 | 0 | 0 | 0 | 0.02 | 0.07778 | 0.02778 | 0 | 0.05 |
| 113 | rs7278627 | 42841333 | 41469406 | A | C | intron_variant | 0 | 0 | 0 | 0 | 0 | 0.02 | 0 | 0.02778 | 0 | 0.05 |
| 114 | rs149109132 | 42841334 | 41469407 | A | G | intron_variant | 0 | 0 | 0 | 0 | 0 | 0 | 0 | 0 | 0 | 0 |
| 115 | rs374608719 | 42841359 | 41469432 | T | C | intron_variant | 0 | 0 | 0 | 0 | 0 | 0 | 0 | 0 | 0 | 0 |
| 116 | rs185416185 | 42841370 | 41469443 | T | C | intron_variant | 0 | 0 | 0 | 0.01282 | 0.009901 | 0 | 0 | 0 | 0 | 0 |
| 117 | rs146052428 | 42841445 | 41469518 | T | C | intron_variant | 0 | 0 | 0 | 0 | 0 | 0 | 0 | 0 | 0 | 0 |
| 118 | rs370663590 | 42841554 | 41469627 | G | C | intron_variant | 0 | 0 | 0 | 0 | 0 | 0 | 0 | 0 | 0 | 0 |
| 119 | rs2070793 | 42841577 | 41469650 | G | A | intron_variant | 0 | 0 | 0.3333 | 0.3077 | 0.2475 | 0.22 | 0.2889 | 0.2222 | 0.1991 | 0.375 |
| 120 | rs117898838 | 42841583 | 41469656 | A | G | intron_variant | 0 | 0 | 0.02083 | 0 | 0 | 0 | 0 | 0 | 0.00463 | 0 |
| 121 | rs2070792 | 42841610 | 41469683 | A | G | intron_variant | 0 | 0 | 0.3333 | 0.3077 | 0.2475 | 0.22 | 0.2889 | 0.2222 | 0.1991 | 0.375 |
| 122 | - | 42841629 | 41469702 | C | A | intron_variant | 0 | 0 | 0 | 0 | 0.00495 | 0 | 0 | 0 | 0 | 0 |
| 123 | rs2070791 | 42841675 | 41469748 | G | C | intron_variant | 0 | 0 | 0 | 0.01282 | 0.00495 | 0 | 0.04444 | 0.08333 | 0.03241 | 0 |
| 124 | rs2070790 | 42841687 | 41469760 | C | G | intron_variant | 0 | 0 | 0.3333 | 0.3077 | 0.2475 | 0.2 | 0.2889 | 0.1944 | 0.1991 | 0.325 |
| 125 | rs373678323 | 42841694 | 41469767 | A | G | intron_variant | 0 | 0 | 0 | 0 | 0 | 0 | 0 | 0 | 0 | 0 |
| 126 | rs142769034 | 42841740 | 41469813 | T | C | intron_variant | 0 | 0 | 0 | 0.01282 | 0.009901 | 0 | 0 | 0 | 0 | 0 |
| 127 | rs111572592 | 42841790 | 41469863 | A | G | intron_variant | 0 | 0 | 0 | 0.05128 | 0.00495 | 0 | 0 | 0 | 0 | 0 |
| 128 | rs2070789 | 42841804 | 41469877 | T | C | intron_variant | 0.3462 | 0.3462 | 0.3125 | 0.2179 | 0.3267 | 0.24 | 0.4667 | 0.4167 | 0.3056 | 0.075 |
| 129 | rs1461069953 | 42841830 | 41469903 | C | T | intron_variant | 0 | 0 | 0 | 0 | 0.00495 | 0 | 0 | 0 | 0 | 0 |
| 130 | rs141603473 | 42841910 | 41469983 | A | C | intron_variant | 0 | 0 | 0 | 0 | 0 | 0 | 0 | 0 | 0 | 0 |
| 131 | rs542946711 | 42841922 | 41469995 | G | A | intron_variant | 0 | 0 | 0 | 0 | 0.009901 | 0 | 0 | 0.02778 | 0 | 0 |
| 132 | rs113564116 | 42841938 | 41470011 | C | T | intron_variant | 0 | 0 | 0.02083 | 0.01282 | 0.0297 | 0 | 0 | 0 | 0 | 0 |
| 133 | - | 42841988 | 41470061 | G | A | intron_variant | 0.3462 | 0.3462 | 0.3542 | 0.4744 | 0.4257 | 0.44 | 0.3222 | 0.3889 | 0.4954 | 0.4 |
| 134 | rs139432971 | 42842046 | 41470119 | G | C | intron_variant | 0 | 0 | 0 | 0 | 0 | 0 | 0 | 0 | 0 | 0 |
| 135 | rs189832305 | 42842079 | 41470152 | T | C | intron_variant | 0 | 0 | 0 | 0 | 0 | 0 | 0 | 0 | 0 | 0 |
| 136 | - | 42842123 | 41470196 | A | C | intron_variant | 0.3462 | 0.3462 | 0.3542 | 0.4744 | 0.4257 | 0.44 | 0.2444 | 0.3889 | 0.4954 | 0.4 |
| 137 | rs2070787 | 42842129 | 41470202 | G | T | intron_variant | 0 | 0 | 0.3333 | 0.3077 | 0.2475 | 0.2 | 0.2778 | 0.1944 | 0.1991 | 0.325 |
| 138 | rs376499029 | 42842152 | 41470225 | C | T | intron_variant | 0 | 0 | 0 | 0 | 0 | 0 | 0 | 0 | 0 | 0 |
| 139 | rs2070786 | 42842203 | 41470276 | C | T | intron_variant | 0 | 0 | 0.3333 | 0.3077 | 0.2475 | 0.2 | 0.2889 | 0.1944 | 0.1991 | 0.325 |
| 140 | rs758426874 | 42842208 | 41470281 | A | G | intron_variant | 0 | 0 | 0 | 0 | 0 | 0 | 0 | 0 | 0.009259 | 0 |
| 141 | - | 42842417 | 41470490 | G | A | intron_variant | 0 | 0 | 0 | 0 | 0 | 0 | 0 | 0 | 0.00463 | 0 |
| 142 | rs547168890 | 42842451 | 41470524 | A | G | intron_variant | 0 | 0 | 0 | 0 | 0 | 0 | 0 | 0 | 0 | 0 |
| 143 | rs373469999 | 42842492 | 41470565 | C | G | intron_variant | 0 | 0 | 0 | 0 | 0 | 0 | 0 | 0 | 0 | 0 |
| 144 | rs1406734513 | 42842516 | 41470589 | A | C | intron_variant | 0 | 0 | 0 | 0 | 0 | 0 | 0 | 0 | 0.00463 | 0 |
| 145 | rs78503214 | 42842543 | 41470616 | T | C | intron_variant | 0 | 0 | 0 | 0 | 0 | 0 | 0 | 0 | 0 | 0 |
| 146 | - | 42842546 | 41470619 | T | C | intron_variant | 0 | 0 | 0 | 0 | 0.00495 | 0 | 0 | 0 | 0 | 0 |
| 147 | rs749103473 | 42842549 | 41470622 | A | T | intron_variant | 0 | 0 | 0 | 0 | 0 | 0 | 0 | 0 | 0 | 0.025 |
| 148 | rs61735794,COSV59824481 | 42842591 | 41470664 | T | C | synonymous_variant | 0 | 0 | 0.02083 | 0.01282 | 0.02475 | 0.02 | 0.01111 | 0 | 0 | 0.05 |
| 149 | - | 42842621 | 41470694 | C | T | synonymous_variant | 0 | 0 | 0 | 0 | 0 | 0 | 0 | 0 | 0.00463 | 0 |
| 150 | rs61735795 | 42842623 | 41470696 | A | G | missense_variant | 0 | 0 | 0 | 0 | 0 | 0 | 0 | 0 | 0 | 0 |
| 151 | rs143712818 | 42842705 | 41470778 | T | G | intron_variant | 0 | 0 | 0 | 0 | 0 | 0 | 0 | 0 | 0 | 0 |
| 152 | rs573482152 | 42842740 | 41470813 | A | T | intron_variant | 0 | 0 | 0 | 0 | 0.00495 | 0 | 0 | 0 | 0 | 0 |
| 153 | rs1160357228 | 42842746 | 41470819 | T | C | intron_variant | 0 | 0 | 0 | 0 | 0 | 0 | 0 | 0 | 0.00463 | 0 |
| 154 | - | 42842759 | 41470832 | A | C | intron_variant | 0 | 0 | 0 | 0 | 0.00495 | 0 | 0 | 0 | 0 | 0 |
| 155 | rs28524972 | 42842771 | 41470844 | G | C | intron_variant | 0 | 0 | 0.3333 | 0.2949 | 0.2376 | 0.2 | 0.2778 | 0.1944 | 0.1991 | 0.325 |
| 156 | - | 42842823 | 41470896 | A | C | intron_variant | 0 | 0 | 0 | 0 | 0 | 0 | 0 | 0 | 0 | 0 |
| 157 | - | 42842834 | 41470907 | T | C | intron_variant | 0.4231 | 0.4231 | 0.0625 | 0.02564 | 0.0297 | 0.12 | 0.2 | 0.3333 | 0.3102 | 0 |
| 158 | rs544946928 | 42842841 | 41470914 | T | C | intron_variant | 0 | 0 | 0 | 0 | 0 | 0.02 | 0 | 0 | 0 | 0.025 |

| | | | | | | | | | | | | | | | | |
|---|---|---|---|---|---|---|---|---|---|---|---|---|---|---|---|---|
| 159 | - | 42842854 | 41470927 | C | A | intron_variant | 0.4231 | 0.4231 | 0.0625 | 0.02564 | 0.0297 | 0.14 | 0.2 | 0.3611 | 0.3102 | 0.05 |
| 160 | - | 42842949 | 41471022 | T | C | intron_variant | 0.4231 | 0.4231 | 0.0625 | 0.02564 | 0.0297 | 0.12 | 0.2 | 0.3333 | 0.3102 | 0 |
| 161 | rs1000072663 | 42842984 | 41471057 | G | A | intron_variant | 0 | 0 | 0 | 0 | 0 | 0 | 0 | 0.02778 | 0.00463 | 0 |
| 162 | rs9305744 | 42842988 | 41471061 | A | G | intron_variant | 0.3462 | 0.3462 | 0.3125 | 0.2308 | 0.3366 | 0.24 | 0.4667 | 0.4167 | 0.3056 | 0.075 |
| 163 | rs35050484 | 42843006 | 41471079 | A | G | intron_variant | 0 | 0 | 0.0625 | 0.0641 | 0.0297 | 0 | 0 | 0 | 0.00463 | 0.05 |
| 164 | rs372665499 | 42843031 | 41471104 | T | C | intron_variant | 0 | 0 | 0 | 0 | 0 | 0 | 0 | 0 | 0 | 0 |
| 165 | rs1198022001 | 42843074 | 41471147 | G | C | intron_variant | 0 | 0 | 0 | 0 | 0 | 0 | 0 | 0 | 0.00463 | 0 |
| 166 | rs375719922,COSV59824983 | 42843078 | 41471151 | C | T | intron_variant | 0 | 0 | 0 | 0 | 0 | 0 | 0.02222 | 0 | 0 | 0 |
| 167 | rs142834250 | 42843107 | 41471180 | G | T | intron_variant | 0 | 0 | 0 | 0 | 0 | 0 | 0 | 0 | 0 | 0 |
| 168 | rs370896169 | 42843236 | 41471309 | C | T | intron_variant | 0 | 0 | 0 | 0 | 0 | 0 | 0 | 0.02778 | 0 | 0 |
| 169 | rs952726028 | 42843281 | 41471354 | C | T | intron_variant | 0 | 0 | 0 | 0 | 0 | 0 | 0 | 0 | 0.00463 | 0 |
| 170 | rs1016288269 | 42843352 | 41471425 | A | G | intron_variant | 0 | 0 | 0 | 0 | 0 | 0 | 0 | 0 | 0 | 0 |
| 171 | rs377410798 | 42843419 | 41471492 | C | G | intron_variant | 0 | 0 | 0 | 0 | 0 | 0 | 0 | 0 | 0 | 0 |
| 172 | rs187052796 | 42843532 | 41471605 | T | C | intron_variant | 0 | 0 | 0 | 0 | 0 | 0 | 0 | 0 | 0 | 0 |
| 173 | rs467375 | 42843565 | 41471638 | A | G | intron_variant | 0.07692 | 0.07692 | 0.2917 | 0.4487 | 0.396 | 0.42 | 0.05556 | 0.02778 | 0.1806 | 0.45 |
| 174 | - | 42843597 | 41471670 | T | C | intron_variant | 0 | 0 | 0 | 0.01282 | 0 | 0 | 0 | 0 | 0 | 0 |
| 175 | rs1015613248 | 42843624 | 41471697 | T | C | intron_variant | 0 | 0 | 0 | 0 | 0 | 0 | 0.01111 | 0 | 0 | 0 |
| 176 | rs116511699 | 42843655 | 41471728 | T | C | intron_variant | 0 | 0 | 0 | 0 | 0 | 0 | 0 | 0 | 0 | 0 |
| 177 | rs370478006 | 42843703 | 41471776 | T | A | intron_variant | 0 | 0 | 0 | 0 | 0 | 0 | 0 | 0 | 0 | 0 |
| 178 | COSV59828007 | 42843855 | 41471928 | C | G | missense_variant | 0.03846 | 0.03846 | 0 | 0 | 0 | 0 | 0 | 0 | 0 | 0 |
| 179 | rs201320664 | 42843935 | 41472008 | C | T | intron_variant | 0 | 0 | 0 | 0 | 0 | 0 | 0 | 0 | 0 | 0 |
| 180 | rs187078345 | 42843940 | 41472013 | A | G | intron_variant | 0 | 0 | 0 | 0 | 0 | 0 | 0.01111 | 0 | 0 | 0 |
| 181 | COSV59823391 | 42843960 | 41472033 | A | G | intron_variant | 0.4231 | 0.4231 | 0.0625 | 0.02564 | 0.0297 | 0.12 | 0.2 | 0.3333 | 0.2917 | 0 |
| 182 | rs937260183 | 42844014 | 41472087 | A | G | intron_variant | 0 | 0 | 0 | 0 | 0 | 0 | 0 | 0 | 0 | 0 |
| 183 | COSV59821944 | 42844052 | 41472125 | T | G | intron_variant | 0.4231 | 0.4231 | 0.0625 | 0.02564 | 0.0297 | 0.12 | 0.2 | 0.3333 | 0.287 | 0 |
| 184 | COSV59824040 | 42844057 | 41472130 | G | A | intron_variant | 0.3462 | 0.3462 | 0.3542 | 0.4744 | 0.4257 | 0.46 | 0.2556 | 0.3611 | 0.4954 | 0.45 |
| 185 | rs7364088 | 42844130 | 41472203 | A | G | intron_variant | 0.3462 | 0.3462 | 0.3333 | 0.2564 | 0.3465 | 0.24 | 0.4667 | 0.4167 | 0.3102 | 0.075 |
| 186 | rs182254633 | 42844168 | 41472241 | G | A | intron_variant | 0 | 0 | 0 | 0 | 0 | 0 | 0 | 0 | 0 | 0 |
| 187 | rs146957681 | 42844174 | 41472247 | T | C | intron_variant | 0 | 0 | 0 | 0 | 0 | 0 | 0 | 0 | 0.009259 | 0 |
| 188 | rs76135088 | 42844202 | 41472275 | G | A | intron_variant | 0 | 0 | 0 | 0 | 0 | 0 | 0 | 0 | 0 | 0 |
| 189 | rs28548447 | 42844238 | 41472311 | A | G | intron_variant | 0 | 0 | 0.3333 | 0.2949 | 0.2475 | 0.22 | 0.2111 | 0.1944 | 0.1991 | 0.325 |
| 190 | rs375524422 | 42844257 | 41472330 | T | C | intron_variant | 0 | 0 | 0 | 0 | 0 | 0 | 0 | 0 | 0 | 0 |
| 191 | rs114844880 | 42844262 | 41472335 | C | T | intron_variant | 0 | 0 | 0 | 0 | 0 | 0 | 0 | 0 | 0 | 0 |
| 192 | rs79065748 | 42844498 | 41472571 | A | C | intron_variant | 0 | 0 | 0 | 0 | 0 | 0 | 0 | 0 | 0 | 0 |
| 193 | rs986882653 | 42844642 | 41472715 | G | C | intron_variant | 0 | 0 | 0 | 0 | 0 | 0 | 0.01111 | 0 | 0 | 0 |
| 194 | rs116606027 | 42844713 | 41472786 | A | G | intron_variant | 0 | 0 | 0 | 0 | 0 | 0 | 0.01111 | 0 | 0 | 0 |
| 195 | rs368644630 | 42844733 | 41472806 | C | T | intron_variant | 0 | 0 | 0 | 0 | 0 | 0 | 0 | 0 | 0 | 0 |
| 196 | rs112980967 | 42844765 | 41472838 | A | G | intron_variant | 0 | 0 | 0 | 0 | 0 | 0 | 0 | 0 | 0 | 0 |
| 197 | rs184164037 | 42844780 | 41472853 | C | T | intron_variant | 0 | 0 | 0 | 0 | 0 | 0 | 0.01111 | 0 | 0 | 0 |
| 198 | - | 42844870 | 41472943 | A | G | intron_variant | 0 | 0 | 0 | 0 | 0 | 0 | 0.01111 | 0 | 0 | 0 |
| 199 | rs73372182 | 42844917 | 41472990 | A | C | intron_variant | 0 | 0 | 0 | 0 | 0 | 0.02 | 0 | 0.02778 | 0 | 0.05 |
| 200 | rs143097165 | 42844925 | 41472998 | A | G | intron_variant | 0 | 0 | 0 | 0 | 0 | 0 | 0.01111 | 0 | 0 | 0 |
| 201 | rs766846682 | 42844994 | 41473067 | T | C | intron_variant | 0 | 0 | 0 | 0 | 0 | 0 | 0 | 0.02778 | 0 | 0 |
| 202 | rs117827300 | 42845065 | 41473138 | A | G | intron_variant | 0 | 0 | 0 | 0 | 0.0198 | 0 | 0 | 0 | 0.06019 | 0 |
| 203 | rs188514624 | 42845101 | 41473174 | C | G | intron_variant | 0 | 0 | 0 | 0 | 0 | 0 | 0.01111 | 0 | 0 | 0 |
| 204 | rs374689537 | 42845160 | 41473233 | T | C | intron_variant | 0 | 0 | 0 | 0 | 0 | 0 | 0.06667 | 0 | 0 | 0 |
| 205 | rs73230068 | 42845167 | 41473240 | C | G | intron_variant | 0 | 0 | 0.02083 | 0.05128 | 0.02475 | 0.02 | 0 | 0 | 0 | 0.05 |
| 206 | rs990674063 | 42845180 | 41473253 | T | C | intron_variant | 0 | 0 | 0 | 0 | 0 | 0 | 0.01111 | 0 | 0 | 0 |
| 207 | rs75168613 | 42845207 | 41473280 | A | G | intron_variant | 0 | 0 | 0 | 0 | 0 | 0 | 0.01111 | 0 | 0 | 0.05 |
| 208 | rs113288437 | 42845216 | 41473289 | G | T | intron_variant | 0 | 0 | 0 | 0 | 0 | 0 | 0.01111 | 0 | 0 | 0.05 |
| 209 | rs140141551,COSV59825540 | 42845220 | 41473293 | A | C | intron_variant | 0 | 0 | 0 | 0.02564 | 0.02475 | 0 | 0 | 0 | 0 | 0.025 |
| 210 | rs774049600 | 42845257 | 41473330 | T | C | synonymous_variant | 0 | 0 | 0 | 0 | 0 | 0 | 0 | 0 | 0 | 0 |
| 211 | rs143291395 | 42845260 | 41473333 | A | G | synonymous_variant | 0 | 0 | 0 | 0 | 0 | 0 | 0 | 0 | 0 | 0 |

| | | | | | | | | | | | | | | | | |
|---|---|---|---|---|---|---|---|---|---|---|---|---|---|---|---|---|
| 212 | rs113436669,COSV59822463,COSV59823505 | 42845278 | 41473351 | A | G | synonymous_variant | 0 | 0 | 0 | 0 | 0 | 0 | 0 | 0 | 0 | 0 |
| 213 | rs2298658 | 42845359 | 41473432 | T | C | synonymous_variant | 0 | 0 | 0 | 0 | 0 | 0 | 0 | 0 | 0.02315 | 0 |
| 214 | rs2298659,COSV59827953 | 42845374 | 41473447 | A | G | synonymous_variant | 0.3462 | 0.3462 | 0.25 | 0.1795 | 0.3168 | 0.16 | 0.1667 | 0.3333 | 0.2454 | 0.075 |
| 215 | rs17854725,COSV59823030 | 42845383 | 41473456 | G | A | synonymous_variant | 0.3462 | 0.3462 | 0.4375 | 0.4744 | 0.4703 | 0.48 | 0.1444 | 0.2222 | 0.3426 | 0.35 |
| 216 | rs756848444 | 42845471 | 41473544 | A | G | intron_variant | 0 | 0 | 0 | 0 | 0.00495 | 0 | 0 | 0 | 0 | 0 |
| 217 | rs377496737 | 42845489 | 41473562 | T | C | intron_variant | 0 | 0 | 0 | 0 | 0.00495 | 0 | 0 | 0 | 0 | 0 |
| 218 | rs113928389 | 42845559 | 41473632 | A | G | intron_variant | 0 | 0 | 0 | 0 | 0 | 0 | 0.01111 | 0 | 0 | 0 |
| 219 | rs2298660 | 42845633 | 41473706 | T | C | intron_variant | 0.3462 | 0.3462 | 0.2083 | 0.1538 | 0.2822 | 0.16 | 0.1667 | 0.3611 | 0.2454 | 0.075 |
| 220 | rs55964536 | 42845638 | 41473711 | T | C | intron_variant | 0.07692 | 0.07692 | 0.2708 | 0.4615 | 0.4158 | 0.42 | 0.04444 | 0.02778 | 0.1852 | 0.4 |
| 221 | rs2298661 | 42845642 | 41473715 | A | C | intron_variant | 0.3462 | 0.3462 | 0.3125 | 0.2436 | 0.3119 | 0.22 | 0.4667 | 0.4167 | 0.3102 | 0.025 |
| 222 | rs1039732893 | 42845655 | 41473728 | A | G | intron_variant | 0 | 0 | 0 | 0 | 0 | 0 | 0.01111 | 0 | 0 | 0 |
| 223 | - | 42845658 | 41473731 | C | G | intron_variant | 0 | 0 | 0 | 0 | 0 | 0 | 0 | 0 | 0 | 0 |
| 224 | rs950562638 | 42845665 | 41473738 | A | G | intron_variant | 0 | 0 | 0 | 0 | 0 | 0 | 0 | 0 | 0.00463 | 0 |
| 225 | rs559830930 | 42845692 | 41473765 | G | A | intron_variant | 0.03846 | 0.03846 | 0 | 0 | 0 | 0 | 0 | 0 | 0 | 0 |
| 226 | rs374850529 | 42845705 | 41473778 | A | G | intron_variant | 0 | 0 | 0 | 0 | 0 | 0 | 0 | 0 | 0 | 0 |
| 227 | rs77511690 | 42845740 | 41473813 | A | G | intron_variant | 0 | 0 | 0 | 0 | 0 | 0 | 0 | 0 | 0 | 0 |
| 228 | rs368994585 | 42845816 | 41473889 | A | G | intron_variant | 0 | 0 | 0 | 0 | 0 | 0 | 0.07778 | 0 | 0 | 0 |
| 229 | rs371053759 | 42845834 | 41473907 | T | G | intron_variant | 0 | 0 | 0 | 0 | 0 | 0 | 0 | 0 | 0 | 0 |
| 230 | - | 42845880 | 41473953 | T | G | intron_variant | 0 | 0 | 0 | 0 | 0 | 0 | 0 | 0.02778 | 0 | 0 |
| 231 | rs3787946 | 42847735 | 41475808 | C | G | intron_variant | 0.3462 | 0.3462 | 0.3125 | 0.2436 | 0.3119 | 0.18 | 0.4778 | 0.4167 | 0.3102 | 0.025 |
| 232 | rs374510753 | 42847763 | 41475836 | T | C | intron_variant | 0 | 0 | 0 | 0 | 0 | 0 | 0.02222 | 0 | 0 | 0 |
| 233 | rs180784757 | 42847764 | 41475837 | A | G | intron_variant | 0 | 0 | 0 | 0 | 0 | 0 | 0 | 0 | 0 | 0 |
| 234 | rs375615614 | 42847773 | 41475846 | A | G | intron_variant | 0 | 0 | 0 | 0 | 0 | 0 | 0 | 0 | 0 | 0 |
| 235 | rs371965055 | 42847798 | 41475871 | T | C | intron_variant | 0 | 0 | 0 | 0 | 0 | 0.02 | 0 | 0 | 0 | 0 |
| 236 | rs117888036 | 42847846 | 41475919 | A | G | intron_variant | 0 | 0 | 0 | 0 | 0 | 0 | 0 | 0 | 0 | 0 |
| 237 | rs368735421 | 42847909 | 41475982 | T | C | intron_variant | 0 | 0 | 0 | 0 | 0 | 0 | 0 | 0 | 0 | 0 |
| 238 | rs66575656 | 42847935 | 41476008 | T | C | intron_variant | 0 | 0 | 0.3542 | 0.2692 | 0.2426 | 0.22 | 0.2222 | 0.1944 | 0.1944 | 0.275 |
| 239 | rs1310846114 | 42848016 | 41476089 | T | C | intron_variant | 0 | 0 | 0 | 0 | 0 | 0 | 0 | 0 | 0.009259 | 0 |
| 240 | rs189425119 | 42848017 | 41476090 | A | G | intron_variant | 0 | 0 | 0 | 0 | 0 | 0 | 0 | 0 | 0 | 0 |
| 241 | - | 42848038 | 41476111 | C | A | intron_variant | 0 | 0 | 0 | 0.01282 | 0 | 0 | 0 | 0 | 0 | 0 |
| 242 | rs80027429 | 42848096 | 41476169 | A | G | intron_variant | 0 | 0 | 0 | 0.03846 | 0.009901 | 0 | 0 | 0 | 0 | 0 |
| 243 | - | 42848115 | 41476188 | G | C | intron_variant | 0.4231 | 0.4231 | 0.0625 | 0.02564 | 0.0297 | 0.12 | 0.1778 | 0.3333 | 0.3102 | 0 |
| 244 | rs367879274 | 42848178 | 41476251 | T | C | intron_variant | 0 | 0 | 0 | 0 | 0 | 0 | 0 | 0 | 0 | 0 |
| 245 | rs8129582,COSV59828343 | 42848183 | 41476256 | A | G | intron_variant | 0 | 0 | 0 | 0 | 0 | 0 | 0 | 0 | 0 | 0 |
| 246 | - | 42848187 | 41476260 | C | T | intron_variant | 0.07692 | 0.07692 | 0.375 | 0.2692 | 0.3416 | 0.36 | 0.2778 | 0.2222 | 0.3796 | 0.075 |
| 247 | rs62217527 | 42848219 | 41476292 | T | C | intron_variant | 0 | 0 | 0.02083 | 0.1026 | 0.08416 | 0.02 | 0 | 0 | 0.009259 | 0.175 |
| 248 | rs112213575 | 42848274 | 41476347 | A | G | intron_variant | 0 | 0 | 0 | 0 | 0 | 0 | 0 | 0 | 0 | 0 |
| 249 | rs527284833 | 42848324 | 41476397 | G | A | intron_variant | 0 | 0 | 0 | 0 | 0 | 0 | 0.01111 | 0 | 0 | 0 |
| 250 | COSV59826599 | 42848363 | 41476436 | A | C | intron_variant | 0 | 0 | 0 | 0 | 0 | 0.02 | 0 | 0 | 0 | 0 |
| 251 | rs113034290 | 42848371 | 41476444 | T | C | intron_variant | 0 | 0 | 0 | 0 | 0 | 0 | 0 | 0 | 0 | 0 |
| 252 | rs111383922 | 42848398 | 41476471 | T | C | intron_variant | 0 | 0 | 0 | 0 | 0 | 0 | 0 | 0 | 0 | 0 |
| 253 | rs75756279,COSV59820906 | 42848457 | 41476530 | T | C | intron_variant | 0 | 0 | 0 | 0.03846 | 0.009901 | 0 | 0 | 0 | 0 | 0 |
| 254 | rs200615061 | 42848494 | 41476567 | C | A | intron_variant | 0 | 0 | 0 | 0.01282 | 0 | 0 | 0 | 0 | 0 | 0 |
| 255 | rs74423429 | 42848579 | 41476652 | A | G | intron_variant | 0 | 0 | 0.04167 | 0.01282 | 0.03465 | 0 | 0 | 0 | 0 | 0 |
| 256 | rs1043973308 | 42848580 | 41476653 | G | A | intron_variant | 0 | 0 | 0 | 0 | 0 | 0 | 0.01111 | 0 | 0 | 0 |
| 257 | rs375891843 | 42848591 | 41476664 | G | T | intron_variant | 0 | 0 | 0 | 0 | 0.00495 | 0 | 0 | 0 | 0 | 0 |
| 258 | - | 42848602 | 41476675 | C | T | intron_variant | 0.03846 | 0.03846 | 0 | 0 | 0 | 0 | 0 | 0 | 0 | 0 |
| 259 | rs1303041038 | 42848640 | 41476713 | A | C | intron_variant | 0 | 0 | 0 | 0 | 0 | 0 | 0 | 0 | 0.00463 | 0 |
| 260 | rs9985159 | 42848684 | 41476757 | T | C | intron_variant | 0.3462 | 0.3462 | 0.3125 | 0.2436 | 0.3119 | 0.24 | 0.4778 | 0.4444 | 0.3102 | 0.075 |
| 261 | rs148136016 | 42848689 | 41476762 | G | A | intron_variant | 0 | 0 | 0 | 0 | 0 | 0 | 0 | 0 | 0 | 0 |
| 262 | rs113506821 | 42848747 | 41476820 | T | C | intron_variant | 0.07692 | 0.07692 | 0.1042 | 0.07692 | 0.08911 | 0.04 | 0 | 0 | 0.1343 | 0 |
| 263 | rs931996263 | 42848765 | 41476838 | C | T | intron_variant | 0 | 0 | 0 | 0 | 0 | 0 | 0.01111 | 0 | 0 | 0 |
| 264 | rs118108663 | 42848819 | 41476892 | T | C | intron_variant | 0 | 0 | 0 | 0 | 0 | 0 | 0 | 0 | 0.03704 | 0 |

| | | | | | | | | | | | | | | | | |
|---|---|---|---|---|---|---|---|---|---|---|---|---|---|---|---|---|
| 265 | rs76973757 | 42848821 | 41476894 | A | G | intron_variant | 0 | 0 | 0 | 0 | 0 | 0 | 0 | 0 | 0 | 0 |
| 266 | - | 42848835 | 41476908 | T | C | intron_variant | 0.07692 | 0.07692 | 0.375 | 0.2692 | 0.3416 | 0.36 | 0.3444 | 0.2222 | 0.3796 | 0.075 |
| 267 | rs1380427461 | 42848924 | 41476997 | C | T | intron_variant | 0 | 0 | 0 | 0 | 0 | 0 | 0 | 0 | 0 | 0.025 |
| 268 | rs375837256 | 42848951 | 41477024 | G | A | intron_variant | 0 | 0 | 0 | 0 | 0 | 0 | 0.03333 | 0 | 0 | 0 |
| 269 | - | 42849089 | 41477162 | C | T | intron_variant | 0 | 0 | 0 | 0 | 0 | 0 | 0 | 0 | 0 | 0.025 |
| 270 | - | 42849097 | 41477170 | G | A | intron_variant | 0.07692 | 0.07692 | 0.3958 | 0.2692 | 0.3515 | 0.36 | 0.3444 | 0.2222 | 0.3796 | 0.075 |
| 271 | rs886737856 | 42849102 | 41477175 | T | C | intron_variant | 0 | 0 | 0 | 0.01282 | 0 | 0 | 0 | 0 | 0 | 0 |
| 272 | - | 42849137 | 41477210 | T | C | intron_variant | 0.07692 | 0.07692 | 0.3958 | 0.2692 | 0.3515 | 0.36 | 0.3444 | 0.2222 | 0.3796 | 0.075 |
| 273 | rs150633108 | 42849163 | 41477236 | T | C | intron_variant | 0 | 0 | 0 | 0 | 0 | 0.04 | 0.01111 | 0 | 0.00463 | 0 |
| 274 | rs58978895 | 42849228 | 41477301 | T | C | intron_variant | 0 | 0 | 0 | 0 | 0 | 0.06 | 0.02222 | 0.02778 | 0 | 0.05 |
| 275 | - | 42849242 | 41477315 | C | T | intron_variant | 0.07692 | 0.07692 | 0.3958 | 0.2821 | 0.3515 | 0.36 | 0.3444 | 0.2222 | 0.3796 | 0.075 |
| 276 | - | 42849258 | 41477331 | C | T | intron_variant | 0.07692 | 0.07692 | 0.3958 | 0.2821 | 0.3515 | 0.36 | 0.3444 | 0.2222 | 0.3796 | 0.075 |
| 277 | rs561179495 | 42849383 | 41477456 | T | C | intron_variant | 0 | 0 | 0 | 0 | 0 | 0 | 0 | 0 | 0 | 0 |
| 278 | rs144576889 | 42849576 | 41477649 | G | A | intron_variant | 0 | 0 | 0 | 0 | 0 | 0 | 0 | 0 | 0 | 0 |
| 279 | rs565720190 | 42849702 | 41477775 | A | G | intron_variant | 0 | 0 | 0 | 0 | 0 | 0.02 | 0 | 0 | 0 | 0 |
| 280 | - | 42849773 | 41477846 | A | C | intron_variant | 0 | 0 | 0 | 0 | 0 | 0 | 0 | 0 | 0.00463 | 0 |
| 281 | - | 42849781 | 41477854 | A | C | intron_variant | 0 | 0 | 0 | 0.01282 | 0.00495 | 0 | 0 | 0 | 0.00463 | 0 |
| 282 | - | 42849783 | 41477856 | A | G | intron_variant | 0 | 0 | 0 | 0 | 0 | 0 | 0 | 0 | 0 | 0 |
| 283 | rs548267325 | 42849802 | 41477875 | A | G | intron_variant | 0 | 0 | 0 | 0 | 0.00495 | 0 | 0 | 0 | 0 | 0 |
| 284 | rs61325328 | 42849821 | 41477894 | G | A | intron_variant | 0 | 0 | 0 | 0 | 0 | 0 | 0 | 0 | 0 | 0 |
| 285 | - | 42849893 | 41477966 | A | G | intron_variant | 0 | 0 | 0 | 0 | 0 | 0 | 0 | 0.02778 | 0 | 0 |
| 286 | rs576215861 | 42849909 | 41477982 | A | C | intron_variant | 0 | 0 | 0 | 0 | 0 | 0 | 0 | 0 | 0 | 0 |
| 287 | rs953081299 | 42849917 | 41477990 | T | C | intron_variant | 0 | 0 | 0 | 0 | 0 | 0 | 0.01111 | 0 | 0 | 0 |
| 288 | rs1259681953 | 42849918 | 41477991 | A | G | intron_variant | 0 | 0 | 0 | 0 | 0 | 0.02 | 0 | 0 | 0 | 0 |
| 289 | - | 42849929 | 41478002 | T | C | intron_variant | 0 | 0 | 0 | 0 | 0 | 0 | 0 | 0 | 0 | 0 |
| 290 | rs115967323 | 42849951 | 41478024 | T | C | intron_variant | 0 | 0 | 0 | 0 | 0 | 0 | 0 | 0 | 0 | 0 |
| 291 | - | 42850038 | 41478111 | C | T | intron_variant | 0 | 0 | 0 | 0 | 0 | 0 | 0 | 0 | 0 | 0.025 |
| 292 | rs9636988 | 42850045 | 41478118 | C | T | intron_variant | 0.2692 | 0.2692 | 0.3333 | 0.2564 | 0.2475 | 0.24 | 0.2222 | 0.1667 | 0.1944 | 0.275 |
| 293 | - | 42850051 | 41478124 | T | G | intron_variant | 0 | 0 | 0 | 0 | 0 | 0 | 0 | 0 | 0 | 0.025 |
| 294 | rs4818239 | 42850075 | 41478148 | C | T | intron_variant | 0.07692 | 0.07692 | 0.2708 | 0.4487 | 0.4158 | 0.4 | 0.04444 | 0.08333 | 0.1759 | 0.325 |
| 295 | rs116170128 | 42850110 | 41478183 | C | A | intron_variant | 0 | 0 | 0.02083 | 0.01282 | 0.03465 | 0.02 | 0.01111 | 0 | 0 | 0.05 |
| 296 | rs543337196 | 42850238 | 41478311 | T | C | intron_variant | 0 | 0 | 0 | 0 | 0 | 0.02 | 0 | 0 | 0 | 0 |
| 297 | rs139816990 | 42850239 | 41478312 | A | G | intron_variant | 0 | 0 | 0 | 0 | 0 | 0 | 0 | 0 | 0 | 0 |
| 298 | rs9983330 | 42850253 | 41478326 | G | A | intron_variant | 0.07692 | 0.07692 | 0.3333 | 0.2692 | 0.3168 | 0.18 | 0.4778 | 0.3889 | 0.3194 | 0.025 |
| 299 | - | 42850281 | 41478354 | T | C | intron_variant | 0 | 0 | 0 | 0 | 0 | 0 | 0 | 0 | 0.00463 | 0 |
| 300 | rs116577479 | 42850370 | 41478443 | A | G | intron_variant | 0 | 0 | 0 | 0 | 0 | 0 | 0 | 0 | 0 | 0 |
| 301 | - | 42850549 | 41478622 | G | A | intron_variant | 0 | 0 | 0 | 0 | 0 | 0 | 0 | 0 | 0.009259 | 0 |
| 302 | rs1339402659 | 42850749 | 41478822 | G | T | intron_variant | 0 | 0 | 0 | 0 | 0 | 0 | 0 | 0.02778 | 0 | 0 |
| 303 | rs374961008 | 42850870 | 41478943 | T | C | intron_variant | 0 | 0 | 0 | 0 | 0 | 0 | 0 | 0 | 0 | 0 |
| 304 | rs73372191 | 42850882 | 41478955 | T | A | intron_variant | 0 | 0 | 0 | 0 | 0 | 0.06 | 0.02222 | 0.02778 | 0 | 0.05 |
| 305 | rs9974995,COSV59826626 | 42850911 | 41478984 | T | C | intron_variant | 0.2692 | 0.2692 | 0.3333 | 0.2692 | 0.2475 | 0.24 | 0.2222 | 0.1667 | 0.1852 | 0.25 |
| 306 | rs766178676 | 42850916 | 41478989 | C | T | intron_variant | 0 | 0 | 0 | 0 | 0 | 0 | 0 | 0 | 0 | 0.025 |
| 307 | rs893281455 | 42850933 | 41479006 | A | C | intron_variant | 0 | 0 | 0 | 0 | 0 | 0 | 0 | 0 | 0 | 0.025 |
| 308 | - | 42850957 | 41479030 | T | C | intron_variant | 0 | 0 | 0 | 0.01282 | 0 | 0 | 0 | 0 | 0 | 0 |
| 309 | rs73372193 | 42850966 | 41479039 | C | A | intron_variant | 0 | 0 | 0 | 0 | 0 | 0.06 | 0.02222 | 0.02778 | 0 | 0.05 |
| 310 | rs9974933,COSV59826632 | 42850977 | 41479050 | G | A | intron_variant | 0.2692 | 0.2692 | 0.3333 | 0.2692 | 0.2475 | 0.24 | 0.2222 | 0.1944 | 0.1852 | 0.25 |
| 311 | rs9975014,COSV59823485 | 42851006 | 41479079 | G | A | intron_variant | 0.2692 | 0.2692 | 0.3333 | 0.2692 | 0.2475 | 0.24 | 0.2222 | 0.1944 | 0.1852 | 0.25 |
| 312 | rs141788162 | 42851134 | 41479207 | A | G | synonymous_variant | 0 | 0 | 0 | 0 | 0.0198 | 0 | 0 | 0 | 0.05556 | 0 |
| 313 | rs201661208 | 42851135 | 41479208 | A | G | missense_variant | 0 | 0 | 0 | 0 | 0 | 0 | 0 | 0 | 0 | 0.025 |
| 314 | rs112247445 | 42851257 | 41479330 | T | C | intron_variant | 0 | 0 | 0 | 0 | 0 | 0 | 0 | 0 | 0 | 0 |
| 315 | rs377385043 | 42851258 | 41479331 | G | A | intron_variant | 0 | 0 | 0 | 0.01282 | 0 | 0 | 0 | 0 | 0 | 0 |
| 316 | rs112985398 | 42851268 | 41479341 | C | T | intron_variant | 0 | 0 | 0 | 0 | 0 | 0 | 0 | 0 | 0 | 0 |
| 317 | rs145292327 | 42851303 | 41479376 | G | A | intron_variant | 0 | 0 | 0 | 0 | 0 | 0 | 0.01111 | 0 | 0 | 0 |

| | | | | | | | | | | | | | | | | |
|---|---|---|---|---|---|---|---|---|---|---|---|---|---|---|---|---|
| 318 | - | 42851310 | 41479383 | A | G | intron_variant | 0 | 0 | 0 | 0.01282 | 0 | 0 | 0 | 0 | 0 | 0 |
| 319 | rs538022064 | 42851370 | 41479443 | C | T | intron_variant | 0 | 0 | 0 | 0 | 0 | 0 | 0 | 0 | 0 | 0 |
| 320 | rs117432315,COSV59822608 | 42851374 | 41479447 | A | G | intron_variant | 0 | 0 | 0.02083 | 0 | 0 | 0 | 0 | 0 | 0 | 0 |
| 321 | rs137871202 | 42851376 | 41479449 | C | T | intron_variant | 0 | 0 | 0 | 0 | 0 | 0 | 0 | 0 | 0 | 0 |
| 322 | rs199575615 | 42851419 | 41479492 | G | A | intron_variant | 0 | 0 | 0 | 0 | 0 | 0 | 0 | 0 | 0.009259 | 0 |
| 323 | rs117941520 | 42851423 | 41479496 | A | G | intron_variant | 0 | 0 | 0 | 0 | 0 | 0.02 | 0 | 0 | 0.00463 | 0 |
| 324 | rs915823 | 42851454 | 41479527 | C | A | intron_variant | 0.07692 | 0.07692 | 0.25 | 0.1795 | 0.2871 | 0.08 | 0.1778 | 0.3333 | 0.2546 | 0.05 |
| 325 | - | 42851575 | 41479648 | G | T | intron_variant | 0 | 0 | 0 | 0.01282 | 0 | 0 | 0 | 0 | 0 | 0 |
| 326 | - | 42851578 | 41479651 | A | C | intron_variant | 0 | 0 | 0.02083 | 0 | 0 | 0 | 0 | 0 | 0 | 0 |
| 327 | - | 42851588 | 41479661 | G | C | intron_variant | 0 | 0 | 0 | 0 | 0 | 0 | 0.03333 | 0 | 0 | 0 |
| 328 | rs184380117 | 42851644 | 41479717 | A | G | intron_variant | 0 | 0 | 0.04167 | 0 | 0 | 0 | 0 | 0 | 0.02778 | 0 |
| 329 | rs553747026 | 42851723 | 41479796 | G | A | intron_variant | 0 | 0 | 0 | 0.01282 | 0 | 0 | 0 | 0 | 0 | 0 |
| 330 | rs1368879178 | 42851727 | 41479800 | G | A | intron_variant | 0 | 0 | 0 | 0 | 0 | 0 | 0 | 0 | 0.01852 | 0 |
| 331 | rs1051561804 | 42851731 | 41479804 | C | T | intron_variant | 0 | 0 | 0 | 0 | 0 | 0 | 0 | 0 | 0 | 0 |
| 332 | rs875393 | 42851815 | 41479888 | A | G | intron_variant | 0 | 0 | 0.1458 | 0.07692 | 0.1139 | 0.04 | 0.1222 | 0.1944 | 0.1944 | 0.025 |
| 333 | rs184764113 | 42851826 | 41479899 | A | G | intron_variant | 0 | 0 | 0 | 0 | 0 | 0 | 0.03333 | 0 | 0 | 0 |
| 334 | rs372876361 | 42851846 | 41479919 | T | G | intron_variant | 0 | 0 | 0 | 0 | 0 | 0 | 0.02222 | 0 | 0 | 0 |
| 335 | - | 42851931 | 41480004 | C | T | intron_variant | 0 | 0 | 0 | 0 | 0 | 0 | 0 | 0 | 0 | 0 |
| 336 | rs149601802 | 42851965 | 41480038 | T | C | intron_variant | 0 | 0 | 0 | 0 | 0 | 0 | 0 | 0 | 0.00463 | 0 |
| 337 | rs967358213 | 42852025 | 41480098 | T | C | intron_variant | 0 | 0 | 0 | 0 | 0 | 0.02 | 0 | 0 | 0 | 0 |
| 338 | rs734055 | 42852170 | 41480243 | A | T | intron_variant | 0 | 0 | 0 | 0 | 0.00495 | 0 | 0 | 0 | 0 | 0 |
| 339 | rs1035659632 | 42852258 | 41480331 | G | T | intron_variant | 0 | 0 | 0 | 0 | 0 | 0 | 0.01111 | 0 | 0 | 0 |
| 340 | rs34561135 | 42852311 | 41480384 | A | G | intron_variant | 0 | 0 | 0.04167 | 0.05128 | 0.08911 | 0.04 | 0 | 0 | 0.02315 | 0.025 |
| 341 | rs373852195 | 42852316 | 41480389 | A | G | intron_variant | 0 | 0 | 0 | 0 | 0.01485 | 0 | 0.06667 | 0 | 0 | 0 |
| 342 | rs734056,COSV59823131 | 42852320 | 41480393 | A | C | intron_variant | 0.07692 | 0.07692 | 0.2708 | 0.4231 | 0.4059 | 0.4 | 0.04444 | 0.05556 | 0.1713 | 0.35 |
| 343 | rs186734573 | 42852365 | 41480438 | T | C | intron_variant | 0 | 0 | 0 | 0 | 0.009901 | 0 | 0 | 0 | 0 | 0 |
| 344 | rs79397218,COSV59825618 | 42852382 | 41480455 | T | C | intron_variant | 0 | 0 | 0 | 0 | 0 | 0 | 0 | 0 | 0 | 0 |
| 345 | rs61735789 | 42852435 | 41480508 | A | G | synonymous_variant | 0 | 0 | 0.02083 | 0.01282 | 0.009901 | 0 | 0 | 0 | 0 | 0 |
| 346 | rs12329760,COSV59820912 | 42852497 | 41480570 | T | C | missense_variant | 0.07692 | 0.07692 | 0.3333 | 0.2821 | 0.3168 | 0.18 | 0.4444 | 0.3056 | 0.3194 | 0.05 |
| 347 | - | 42852545 | 41480618 | A | G | intron_variant | 0 | 0 | 0 | 0 | 0 | 0 | 0.01111 | 0 | 0 | 0 |
| 348 | rs378501 | 42852591 | 41480664 | G | A | intron_variant | 0 | 0 | 0 | 0 | 0 | 0 | 0.05556 | 0.1111 | 0 | 0 |
| 349 | rs1601575263 | 42852673 | 41480746 | A | C | intron_variant | 0 | 0 | 0 | 0 | 0 | 0 | 0 | 0 | 0.00463 | 0 |
| 350 | rs1181456418 | 42852722 | 41480795 | T | C | intron_variant | 0 | 0 | 0 | 0 | 0 | 0 | 0 | 0 | 0.00463 | 0 |
| 351 | rs370654859 | 42852797 | 41480870 | C | T | intron_variant | 0 | 0 | 0 | 0 | 0 | 0 | 0 | 0 | 0 | 0 |
| 352 | - | 42852847 | 41480920 | A | G | intron_variant | 0 | 0 | 0 | 0 | 0 | 0 | 0 | 0 | 0 | 0 |
| 353 | rs866828358 | 42852904 | 41480977 | T | C | intron_variant | 0 | 0 | 0.02083 | 0 | 0 | 0 | 0 | 0 | 0 | 0 |
| 354 | rs935832638 | 42852907 | 41480980 | A | G | intron_variant | 0 | 0 | 0 | 0 | 0 | 0 | 0 | 0 | 0 | 0 |
| 355 | rs112209215 | 42852922 | 41480995 | A | G | intron_variant | 0 | 0 | 0 | 0 | 0 | 0 | 0 | 0 | 0 | 0 |
| 356 | rs75833467,COSV59822764 | 42852952 | 41481025 | A | G | intron_variant | 0 | 0 | 0 | 0.03846 | 0.03465 | 0 | 0 | 0 | 0.00463 | 0.05 |
| 357 | - | 42853026 | 41481099 | C | T | intron_variant | 0.03846 | 0.03846 | 0 | 0 | 0 | 0 | 0 | 0 | 0 | 0 |
| 358 | rs370641046 | 42853045 | 41481118 | T | C | intron_variant | 0 | 0 | 0 | 0 | 0 | 0 | 0 | 0 | 0 | 0.025 |
| 359 | rs4290734 | 42853083 | 41481156 | G | A | intron_variant | 0.07692 | 0.07692 | 0.2708 | 0.4231 | 0.4059 | 0.4 | 0.04444 | 0.05556 | 0.1713 | 0.375 |
| 360 | rs1244375280 | 42853121 | 41481194 | A | G | intron_variant | 0 | 0 | 0 | 0 | 0 | 0 | 0 | 0 | 0 | 0 |
| 361 | rs1569018086 | 42853239 | 41481312 | T | C | intron_variant | 0 | 0 | 0 | 0 | 0 | 0 | 0.01111 | 0 | 0 | 0 |
| 362 | rs149708827 | 42853243 | 41481316 | T | G | intron_variant | 0 | 0 | 0 | 0 | 0 | 0 | 0.04444 | 0 | 0 | 0 |
| 363 | - | 42853301 | 41481374 | A | G | intron_variant | 0 | 0 | 0 | 0 | 0 | 0 | 0 | 0 | 0.00463 | 0 |
| 364 | rs544784789 | 42853341 | 41481414 | G | A | intron_variant | 0 | 0 | 0 | 0 | 0 | 0.02 | 0 | 0 | 0 | 0 |
| 365 | rs1312825450 | 42853470 | 41481543 | C | T | intron_variant | 0 | 0 | 0 | 0 | 0 | 0 | 0 | 0 | 0.00463 | 0 |
| 366 | rs952095916 | 42853500 | 41481573 | A | T | intron_variant | 0 | 0 | 0 | 0 | 0 | 0 | 0.01111 | 0 | 0 | 0 |
| 367 | rs200072801 | 42854102 | 41482175 | G | A | intron_variant | 0 | 0 | 0 | 0 | 0 | 0 | 0 | 0.02778 | 0 | 0 |
| 368 | - | 42854118 | 41482191 | G | T | intron_variant | 0 | 0 | 0 | 0.01282 | 0 | 0 | 0 | 0 | 0 | 0 |
| 369 | rs111620846 | 42854144 | 41482217 | A | C | intron_variant | 0 | 0 | 0 | 0 | 0 | 0 | 0 | 0 | 0 | 0 |
| 370 | rs2276205,COSV59820921 | 42854203 | 41482276 | G | A | intron_variant | 0 | 0 | 0.1875 | 0.1026 | 0.1287 | 0.04 | 0.1222 | 0.1667 | 0.2546 | 0.025 |

| | | | | | | | | | | | | | | | | |
|---|---|---|---|---|---|---|---|---|---|---|---|---|---|---|---|---|
| 371 | rs578103765,COSV59821991 | 42854205 | 41482278 | T | C | intron_variant | 0 | 0 | 0 | 0 | 0 | 0 | 0 | 0 | 0 | 0 |
| 372 | rs144033559 | 42854343 | 41482416 | C | G | intron_variant | 0 | 0 | 0 | 0 | 0 | 0 | 0 | 0 | 0 | 0 |
| 373 | rs118134524 | 42854376 | 41482449 | A | G | intron_variant | 0 | 0 | 0 | 0 | 0 | 0 | 0 | 0.02778 | 0.00463 | 0 |
| 374 | rs527489879 | 42854400 | 41482473 | C | T | intron_variant | 0 | 0 | 0 | 0 | 0 | 0 | 0 | 0.02778 | 0 | 0 |
| 375 | rs930977615 | 42854436 | 41482509 | C | T | intron_variant | 0 | 0 | 0 | 0 | 0 | 0 | 0.01111 | 0 | 0 | 0 |
| 376 | rs112753686 | 42854460 | 41482533 | T | C | intron_variant | 0 | 0 | 0 | 0 | 0 | 0 | 0 | 0 | 0 | 0 |
| 377 | rs900993466 | 42854525 | 41482598 | C | T | intron_variant | 0 | 0 | 0 | 0 | 0 | 0 | 0.01111 | 0 | 0 | 0 |
| 378 | - | 42854637 | 41482710 | C | G | intron_variant | 0 | 0 | 0 | 0 | 0 | 0.02 | 0 | 0 | 0 | 0 |
| 379 | rs34783969 | 42854638 | 41482711 | T | A | intron_variant | 0.07692 | 0.07692 | 0.2708 | 0.4231 | 0.401 | 0.4 | 0.04444 | 0.05556 | 0.1713 | 0.375 |
| 380 | rs980528078 | 42854659 | 41482732 | C | G | intron_variant | 0 | 0 | 0 | 0 | 0 | 0 | 0 | 0 | 0 | 0 |
| 381 | rs1309520200 | 42854715 | 41482788 | T | C | intron_variant | 0 | 0 | 0 | 0 | 0 | 0 | 0.01111 | 0 | 0 | 0 |
| 382 | rs544657781 | 42854830 | 41482903 | C | A | intron_variant | 0 | 0 | 0 | 0 | 0 | 0 | 0 | 0 | 0 | 0 |
| 383 | rs554462605 | 42854914 | 41482987 | A | G | intron_variant | 0 | 0 | 0 | 0 | 0 | 0 | 0 | 0 | 0 | 0 |
| 384 | - | 42854923 | 41482996 | A | C | intron_variant | 0 | 0 | 0 | 0 | 0 | 0 | 0.01111 | 0 | 0 | 0 |
| 385 | rs143818732,COSV59820925 | 42855131 | 41483204 | C | T | intron_variant | 0 | 0 | 0 | 0.07692 | 0.009901 | 0 | 0 | 0 | 0 | 0 |
| 386 | - | 42855215 | 41483288 | G | A | intron_variant | 0 | 0 | 0 | 0 | 0 | 0 | 0 | 0 | 0 | 0 |
| 387 | rs9977234,COSV59822704 | 42855564 | 41483637 | T | G | intron_variant | 0.2692 | 0.2692 | 0.2917 | 0.2436 | 0.2376 | 0.24 | 0.2222 | 0.2222 | 0.1343 | 0.25 |
| 388 | rs9980225 | 42855798 | 41483871 | C | T | intron_variant | 0.07692 | 0.07692 | 0.1042 | 0.1154 | 0.1634 | 0.04 | 0.02222 | 0.08333 | 0.05093 | 0 |
| 389 | rs75430506 | 42855856 | 41483929 | A | G | intron_variant | 0 | 0 | 0.02083 | 0 | 0.00495 | 0 | 0 | 0.05556 | 0.01852 | 0 |
| 390 | rs1230739206 | 42855862 | 41483935 | A | C | intron_variant | 0 | 0 | 0 | 0 | 0.00495 | 0 | 0 | 0 | 0 | 0 |
| 391 | - | 42856048 | 41484121 | A | C | intron_variant | 0.1538 | 0.1538 | 0.3542 | 0.2692 | 0.2673 | 0.36 | 0.4222 | 0.3889 | 0.463 | 0.275 |
| 392 | rs182231135 | 42856065 | 41484138 | C | T | intron_variant | 0.03846 | 0.03846 | 0 | 0 | 0 | 0 | 0 | 0 | 0 | 0 |
| 393 | rs141091900 | 42856099 | 41484172 | C | G | intron_variant | 0 | 0 | 0.04167 | 0 | 0.009901 | 0 | 0 | 0 | 0.01852 | 0 |
| 394 | - | 42856116 | 41484189 | A | G | intron_variant | 0.4231 | 0.4231 | 0.0625 | 0.02564 | 0.0297 | 0.12 | 0.1889 | 0.3889 | 0.3102 | 0 |
| 395 | rs146865760,COSV59826554 | 42856216 | 41484289 | T | C | intron_variant | 0 | 0 | 0 | 0 | 0.01485 | 0 | 0 | 0 | 0 | 0 |
| 396 | rs139222305 | 42856338 | 41484411 | T | C | intron_variant | 0 | 0 | 0 | 0.02564 | 0.009901 | 0 | 0 | 0 | 0 | 0.025 |
| 397 | rs372976235 | 42856379 | 41484452 | C | G | intron_variant | 0 | 0 | 0 | 0 | 0 | 0 | 0 | 0 | 0 | 0 |
| 398 | rs541215881 | 42856397 | 41484470 | A | G | intron_variant | 0.03846 | 0.03846 | 0 | 0 | 0 | 0 | 0 | 0 | 0 | 0 |
| 399 | rs56066678,COSV59822708 | 42856479 | 41484552 | T | C | intron_variant | 0.2692 | 0.2692 | 0.2917 | 0.2436 | 0.2376 | 0.32 | 0.2556 | 0.2222 | 0.1528 | 0.325 |
| 400 | rs2838039 | 42856544 | 41484617 | C | T | intron_variant | 0.07692 | 0.07692 | 0.375 | 0.3077 | 0.3317 | 0.16 | 0.4444 | 0.3333 | 0.3657 | 0.05 |
| 401 | rs147054075 | 42856619 | 41484692 | A | G | intron_variant | 0 | 0 | 0 | 0 | 0 | 0.02 | 0 | 0 | 0 | 0 |
| 402 | rs549990870 | 42856711 | 41484784 | G | A | intron_variant | 0 | 0 | 0 | 0.01282 | 0 | 0 | 0 | 0 | 0 | 0 |
| 403 | rs138164084 | 42856742 | 41484815 | T | G | intron_variant | 0 | 0 | 0 | 0 | 0 | 0 | 0 | 0 | 0 | 0 |
| 404 | - | 42856756 | 41484829 | G | A | intron_variant | 0.4231 | 0.4231 | 0.0625 | 0.02564 | 0.0297 | 0.12 | 0.1889 | 0.3889 | 0.3102 | 0 |
| 405 | - | 42856774 | 41484847 | T | C | intron_variant | 0 | 0 | 0 | 0 | 0 | 0 | 0 | 0 | 0 | 0 |
| 406 | rs79243099 | 42856901 | 41484974 | A | C | intron_variant | 0 | 0 | 0 | 0 | 0 | 0 | 0 | 0 | 0 | 0 |
| 407 | - | 42856949 | 41485022 | T | C | intron_variant | 0.4231 | 0.4231 | 0.0625 | 0.02564 | 0.0297 | 0.12 | 0.1889 | 0.3889 | 0.3102 | 0 |
| 408 | rs149551068 | 42856975 | 41485048 | A | C | intron_variant | 0 | 0 | 0 | 0 | 0 | 0 | 0 | 0 | 0 | 0 |
| 409 | rs762602293 | 42857262 | 41485335 | G | T | intron_variant | 0 | 0 | 0 | 0.01282 | 0 | 0 | 0 | 0 | 0 | 0 |
| 410 | - | 42857302 | 41485375 | T | C | intron_variant | 0.4231 | 0.4231 | 0.0625 | 0.02564 | 0.0297 | 0.18 | 0.2111 | 0.4167 | 0.3056 | 0.05 |
| 411 | rs9983252 | 42857322 | 41485395 | G | C | intron_variant | 0.07692 | 0.07692 | 0.375 | 0.3077 | 0.3317 | 0.16 | 0.4444 | 0.3333 | 0.3704 | 0.05 |
| 412 | - | 42857444 | 41485517 | C | T | intron_variant | 0.4231 | 0.4231 | 0.0625 | 0.02564 | 0.0297 | 0.18 | 0.2222 | 0.4167 | 0.3056 | 0.05 |
| 413 | rs4818242 | 42857557 | 41485630 | T | C | intron_variant | 0.1923 | 0.1923 | 0 | 0 | 0 | 0 | 0 | 0 | 0.00463 | 0 |
| 414 | rs11702475,COSV59824994 | 42857568 | 41485641 | T | C | intron_variant | 0.07692 | 0.07692 | 0.2708 | 0.4231 | 0.396 | 0.42 | 0.04444 | 0.05556 | 0.1713 | 0.4 |
| 415 | - | 42857642 | 41485715 | T | C | intron_variant | 0 | 0 | 0 | 0 | 0 | 0 | 0.02222 | 0 | 0 | 0 |
| 416 | rs2257202,COSV59822713 | 42857715 | 41485788 | C | T | intron_variant | 0.2692 | 0.2692 | 0.2917 | 0.2436 | 0.2426 | 0.24 | 0.2222 | 0.1944 | 0.1528 | 0.25 |
| 417 | rs746347660 | 42857750 | 41485823 | A | G | intron_variant | 0 | 0 | 0 | 0 | 0 | 0 | 0 | 0 | 0 | 0.05 |
| 418 | rs118108194 | 42857755 | 41485828 | G | A | intron_variant | 0 | 0 | 0 | 0 | 0 | 0 | 0 | 0 | 0 | 0 |
| 419 | rs1043408573 | 42857809 | 41485882 | T | C | intron_variant | 0 | 0 | 0.02083 | 0 | 0 | 0 | 0 | 0 | 0 | 0 |
| 420 | rs752423992 | 42857830 | 41485903 | T | C | intron_variant | 0 | 0 | 0 | 0 | 0.00495 | 0 | 0 | 0 | 0 | 0 |
| 421 | rs1262043781 | 42857893 | 41485966 | G | A | intron_variant | 0 | 0 | 0.02083 | 0 | 0 | 0 | 0 | 0 | 0 | 0 |
| 422 | rs62217531 | 42857901 | 41485974 | T | C | intron_variant | 0.07692 | 0.07692 | 0.2292 | 0.3974 | 0.3861 | 0.46 | 0.06667 | 0.08333 | 0.1667 | 0.375 |
| 423 | rs1601579713 | 42857940 | 41486013 | T | G | intron_variant | 0 | 0 | 0 | 0 | 0.00495 | 0 | 0 | 0 | 0 | 0 |

| | | | | | | | | | | | | | | | | |
|---|---|---|---|---|---|---|---|---|---|---|---|---|---|---|---|---|
| 424 | rs144052153 | 42857954 | 41486027 | G | T | intron_variant | 0 | 0 | 0 | 0 | 0 | 0 | 0 | 0 | 0.02315 | 0 |
| 425 | rs2298857,COSV59822719 | 42857981 | 41486054 | A | G | intron_variant | 0.2692 | 0.2692 | 0.2917 | 0.2436 | 0.2426 | 0.3 | 0.2444 | 0.2222 | 0.1528 | 0.35 |
| 426 | - | 42857985 | 41486058 | C | T | intron_variant | 0 | 0 | 0 | 0 | 0 | 0 | 0 | 0 | 0 | 0 |
| 427 | - | 42858054 | 41486127 | C | G | intron_variant | 0 | 0 | 0 | 0 | 0 | 0 | 0 | 0 | 0 | 0 |
| 428 | rs572593241 | 42858080 | 41486153 | T | C | intron_variant | 0 | 0 | 0 | 0 | 0 | 0 | 0 | 0 | 0 | 0.025 |
| 429 | rs182791541 | 42858096 | 41486169 | T | A | intron_variant | 0 | 0 | 0 | 0 | 0 | 0 | 0 | 0 | 0 | 0.025 |
| 430 | rs539934676 | 42858142 | 41486215 | A | G | intron_variant | 0 | 0 | 0 | 0 | 0 | 0 | 0 | 0 | 0 | 0.025 |
| 431 | - | 42858367 | 41486440 | T | C | intron_variant | 0.3077 | 0.3077 | 0.2917 | 0.4231 | 0.4109 | 0.42 | 0.2889 | 0.4167 | 0.4491 | 0.375 |
| 432 | rs752183643 | 42858375 | 41486448 | T | C | intron_variant | 0 | 0 | 0 | 0 | 0 | 0 | 0 | 0 | 0 | 0 |
| 433 | rs564834372 | 42858408 | 41486481 | C | G | intron_variant | 0 | 0 | 0 | 0 | 0.00495 | 0 | 0 | 0 | 0 | 0 |
| 434 | rs139467735 | 42858441 | 41486514 | T | G | intron_variant | 0 | 0 | 0 | 0 | 0 | 0 | 0 | 0 | 0 | 0.025 |
| 435 | rs546775580 | 42858460 | 41486533 | T | C | intron_variant | 0 | 0 | 0 | 0 | 0 | 0 | 0.01111 | 0 | 0 | 0 |
| 436 | rs182336392 | 42858499 | 41486572 | G | A | intron_variant | 0 | 0 | 0 | 0 | 0.00495 | 0 | 0 | 0 | 0 | 0 |
| 437 | rs149676870 | 42858520 | 41486593 | T | C | intron_variant | 0.2692 | 0.2692 | 0 | 0 | 0 | 0 | 0 | 0 | 0 | 0 |
| 438 | - | 42858549 | 41486622 | A | G | intron_variant | 0 | 0 | 0 | 0 | 0 | 0 | 0.01111 | 0 | 0 | 0 |
| 439 | rs779138284 | 42858594 | 41486667 | T | C | intron_variant | 0 | 0 | 0 | 0 | 0.00495 | 0 | 0 | 0 | 0 | 0 |
| 440 | rs193122175 | 42858697 | 41486770 | T | C | intron_variant | 0 | 0 | 0 | 0 | 0 | 0 | 0 | 0 | 0 | 0 |
| 441 | rs364289,COSV59822727 | 42858865 | 41486938 | A | G | intron_variant | 0.2692 | 0.2692 | 0.2917 | 0.2308 | 0.2376 | 0.3 | 0.2778 | 0.3056 | 0.1528 | 0.35 |
| 442 | - | 42858868 | 41486941 | G | A | intron_variant | 0.03846 | 0.03846 | 0 | 0 | 0 | 0 | 0 | 0 | 0 | 0 |
| 443 | rs370363307 | 42858874 | 41486947 | T | A | intron_variant | 0 | 0 | 0 | 0 | 0 | 0 | 0.03333 | 0.02778 | 0.009259 | 0 |
| 444 | rs149226336 | 42858936 | 41487009 | G | C | intron_variant | 0 | 0 | 0 | 0.01282 | 0.00495 | 0 | 0.03333 | 0.02778 | 0.03241 | 0 |
| 445 | rs903787356 | 42858958 | 41487031 | C | T | intron_variant | 0 | 0 | 0.02083 | 0 | 0 | 0 | 0 | 0 | 0 | 0 |
| 446 | - | 42858990 | 41487063 | C | G | intron_variant | 0 | 0 | 0 | 0 | 0 | 0 | 0.02222 | 0 | 0 | 0 |
| 447 | rs79468500,COSV59824518 | 42858995 | 41487068 | G | T | intron_variant | 0 | 0 | 0.02083 | 0.01282 | 0.03465 | 0.02 | 0.01111 | 0 | 0 | 0.025 |
| 448 | rs80275470 | 42859054 | 41487127 | A | G | intron_variant | 0 | 0 | 0 | 0 | 0.0198 | 0 | 0 | 0 | 0 | 0 |
| 449 | rs365025,COSV59827729 | 42859067 | 41487140 | C | G | intron_variant | 0.3462 | 0.3462 | 0.4792 | 0.3846 | 0.3713 | 0.3 | 0.3222 | 0.3611 | 0.3194 | 0.1 |
| 450 | rs193067129 | 42859193 | 41487266 | A | G | intron_variant | 0 | 0 | 0 | 0 | 0 | 0 | 0 | 0 | 0 | 0 |
| 451 | rs365724 | 42859222 | 41487295 | C | G | intron_variant | 0.3462 | 0.3462 | 0.4792 | 0.3718 | 0.3713 | 0.3 | 0.3222 | 0.3611 | 0.3194 | 0.1 |
| 452 | rs188508398 | 42859319 | 41487392 | C | G | intron_variant | 0 | 0 | 0 | 0.01282 | 0 | 0 | 0 | 0 | 0 | 0 |
| 453 | rs372400372 | 42859325 | 41487398 | G | A | intron_variant | 0 | 0 | 0 | 0 | 0 | 0 | 0 | 0 | 0 | 0 |
| 454 | - | 42859604 | 41487677 | G | A | intron_variant | 0.4231 | 0.4231 | 0.0625 | 0.02564 | 0.0297 | 0.12 | 0.1667 | 0.3333 | 0.287 | 0 |
| 455 | - | 42859670 | 41487743 | T | C | intron_variant | 0.4231 | 0.4231 | 0.0625 | 0.02564 | 0.0297 | 0.12 | 0.2222 | 0.3333 | 0.287 | 0 |
| 456 | rs375760,COSV59824141 | 42859686 | 41487759 | T | G | intron_variant | 0.2692 | 0.2692 | 0.2917 | 0.2308 | 0.2376 | 0.26 | 0.2444 | 0.2778 | 0.1528 | 0.25 |
| 457 | rs147063811 | 42859733 | 41487806 | T | A | intron_variant | 0 | 0 | 0 | 0 | 0 | 0 | 0 | 0 | 0 | 0 |
| 458 | rs415918 | 42859876 | 41487949 | T | C | intron_variant | 0.3462 | 0.3462 | 0.4792 | 0.3718 | 0.3713 | 0.3 | 0.3222 | 0.3611 | 0.3194 | 0.1 |
| 459 | rs141888586 | 42859971 | 41488044 | T | C | intron_variant | 0 | 0 | 0 | 0 | 0 | 0 | 0 | 0 | 0 | 0 |
| 460 | - | 42859999 | 41488072 | T | C | intron_variant | 0 | 0 | 0 | 0 | 0 | 0.02 | 0 | 0 | 0 | 0 |
| 461 | rs528451616 | 42860109 | 41488182 | A | G | intron_variant | 0 | 0 | 0 | 0 | 0 | 0 | 0 | 0 | 0 | 0 |
| 462 | rs56136037,COSV59822959 | 42860136 | 41488209 | T | G | intron_variant | 0 | 0 | 0.04167 | 0 | 0.05446 | 0.02 | 0 | 0 | 0.00463 | 0 |
| 463 | rs761668242 | 42860295 | 41488368 | T | C | intron_variant | 0 | 0 | 0 | 0 | 0 | 0 | 0 | 0.02778 | 0 | 0 |
| 464 | rs371531071 | 42860299 | 41488372 | T | C | intron_variant | 0 | 0 | 0 | 0 | 0 | 0 | 0.02222 | 0 | 0 | 0 |
| 465 | rs422471,COSV59821998 | 42860307 | 41488380 | T | C | intron_variant | 0.3462 | 0.3462 | 0.4792 | 0.3718 | 0.3762 | 0.3 | 0.3222 | 0.3611 | 0.3194 | 0.1 |
| 466 | COSV59827512 | 42860336 | 41488409 | T | C | missense_variant | 0 | 0 | 0 | 0 | 0.00495 | 0 | 0 | 0 | 0 | 0 |
| 467 | rs386416,COSV59823496 | 42860485 | 41488558 | C | G | intron_variant | 0.3462 | 0.3462 | 0.4792 | 0.3718 | 0.3713 | 0.3 | 0.3222 | 0.3611 | 0.3194 | 0.1 |
| 468 | rs187290362 | 42860488 | 41488561 | G | A | intron_variant | 0 | 0 | 0 | 0 | 0 | 0 | 0 | 0 | 0 | 0.05 |
| 469 | rs3819138,COSV59826644 | 42860494 | 41488567 | G | C | intron_variant | 0.07692 | 0.07692 | 0.1042 | 0.1154 | 0.1683 | 0.02 | 0.03333 | 0.05556 | 0.05093 | 0 |
| 470 | rs536039173 | 42860514 | 41488587 | T | G | intron_variant | 0 | 0 | 0.02083 | 0.01282 | 0.00495 | 0 | 0 | 0 | 0 | 0 |
| 471 | rs555995855 | 42860520 | 41488593 | C | T | intron_variant | 0 | 0 | 0 | 0 | 0 | 0.06 | 0 | 0 | 0 | 0 |
| 472 | rs117696554,COSV59824532 | 42860532 | 41488605 | A | G | intron_variant | 0 | 0 | 0.02083 | 0.01282 | 0.02475 | 0.02 | 0.01111 | 0 | 0 | 0.025 |
| 473 | rs3787947 | 42860593 | 41488666 | T | C | intron_variant | 0.07692 | 0.07692 | 0.4167 | 0.3333 | 0.3366 | 0.18 | 0.4222 | 0.2778 | 0.3611 | 0.1 |
| 474 | rs910522814 | 42860603 | 41488676 | T | C | intron_variant | 0 | 0 | 0 | 0.01282 | 0 | 0 | 0 | 0 | 0 | 0 |
| 475 | - | 42860648 | 41488721 | C | T | intron_variant | 0 | 0 | 0 | 0 | 0 | 0 | 0 | 0 | 0 | 0.025 |
| 476 | rs139305247 | 42860655 | 41488728 | T | C | intron_variant | 0.07692 | 0.07692 | 0.04167 | 0.01282 | 0.03465 | 0 | 0 | 0 | 0.01389 | 0 |

| | | | | | | | | | | | | | | | | |
|---|---|---|---|---|---|---|---|---|---|---|---|---|---|---|---|---|
| 477 | rs1210853029 | 42860713 | 41488786 | T | C | intron_variant | 0 | 0 | 0 | 0 | 0.00495 | 0 | 0 | 0 | 0 | 0 |
| 478 | rs423596,COSV59825578 | 42860809 | 41488882 | T | C | intron_variant | 0 | 0 | 0.1042 | 0.0641 | 0.05446 | 0.14 | 0.2333 | 0.25 | 0.125 | 0.075 |
| 479 | - | 42860838 | 41488911 | T | G | intron_variant | 0 | 0 | 0 | 0 | 0.00495 | 0 | 0 | 0 | 0 | 0 |
| 480 | rs145996484 | 42860853 | 41488926 | G | C | intron_variant | 0 | 0 | 0 | 0 | 0 | 0 | 0 | 0 | 0 | 0 |
| 481 | rs1230361255 | 42860887 | 41488960 | A | G | intron_variant | 0 | 0 | 0 | 0 | 0 | 0 | 0 | 0 | 0.009259 | 0 |
| 482 | rs532082574 | 42861129 | 41489202 | G | C | intron_variant | 0 | 0 | 0 | 0 | 0 | 0 | 0 | 0 | 0 | 0 |
| 483 | - | 42861143 | 41489216 | T | C | intron_variant | 0 | 0 | 0 | 0 | 0 | 0 | 0 | 0 | 0 | 0 |
| 484 | rs2298664 | 42861181 | 41489254 | C | G | intron_variant | 0.07692 | 0.07692 | 0.4167 | 0.3462 | 0.3416 | 0.18 | 0.4556 | 0.3056 | 0.3935 | 0.1 |
| 485 | rs1191882308 | 42861203 | 41489276 | A | G | intron_variant | 0 | 0 | 0 | 0 | 0 | 0 | 0.01111 | 0 | 0 | 0 |
| 486 | rs1426886770,COSV59823937 | 42861289 | 41489362 | T | G | intron_variant | 0 | 0 | 0.02083 | 0 | 0 | 0 | 0 | 0 | 0 | 0 |
| 487 | rs429442,COSV59824149 | 42861332 | 41489405 | T | C | intron_variant | 0.2692 | 0.2692 | 0.2917 | 0.2308 | 0.2426 | 0.32 | 0.2667 | 0.3056 | 0.1528 | 0.325 |
| 488 | rs151152524 | 42861368 | 41489441 | C | A | intron_variant | 0 | 0 | 0 | 0 | 0 | 0 | 0 | 0 | 0 | 0 |
| 489 | rs768426172,COSV59824319 | 42861473 | 41489546 | T | C | missense_variant | 0 | 0 | 0 | 0 | 0 | 0 | 0.02222 | 0 | 0 | 0 |
| 490 | rs190265904 | 42861476 | 41489549 | A | G | missense_variant | 0 | 0 | 0 | 0 | 0 | 0 | 0 | 0 | 0.01389 | 0 |
| 491 | rs144948620,COSV59826562 | 42861545 | 41489618 | A | G | intron_variant | 0 | 0 | 0 | 0 | 0.01485 | 0 | 0 | 0 | 0 | 0 |
| 492 | rs371894730 | 42861591 | 41489664 | C | T | intron_variant | 0 | 0 | 0 | 0 | 0 | 0 | 0 | 0 | 0 | 0 |
| 493 | rs373622147 | 42861764 | 41489837 | C | T | intron_variant | 0 | 0 | 0 | 0 | 0 | 0 | 0 | 0 | 0 | 0 |
| 494 | rs141764184 | 42861794 | 41489867 | A | G | intron_variant | 0 | 0 | 0 | 0 | 0 | 0 | 0.08889 | 0 | 0 | 0 |
| 495 | rs199704982 | 42861935 | 41490008 | T | C | intron_variant | 0 | 0 | 0 | 0 | 0 | 0 | 0 | 0 | 0.02315 | 0 |
| 496 | rs192666663 | 42861936 | 41490009 | A | G | intron_variant | 0 | 0 | 0 | 0 | 0 | 0 | 0 | 0.02778 | 0 | 0 |
| 497 | rs147099383 | 42861959 | 41490032 | C | T | intron_variant | 0 | 0 | 0.08333 | 0.07692 | 0.0297 | 0.1 | 0.3333 | 0.08333 | 0.06481 | 0 |
| 498 | rs11911394 | 42862275 | 41490348 | C | T | intron_variant | 0.07692 | 0.07692 | 0.4167 | 0.3333 | 0.3366 | 0.24 | 0.4556 | 0.3056 | 0.3611 | 0.175 |
| 499 | rs189040987 | 42862333 | 41490406 | C | G | intron_variant | 0 | 0 | 0 | 0 | 0 | 0 | 0 | 0 | 0.00463 | 0 |
| 500 | rs561412657 | 42862367 | 41490440 | G | A | intron_variant | 0 | 0 | 0 | 0 | 0.00495 | 0 | 0 | 0 | 0 | 0 |
| 501 | rs192259532 | 42862416 | 41490489 | T | G | intron_variant | 0 | 0 | 0 | 0 | 0 | 0 | 0 | 0 | 0 | 0 |
| 502 | rs1315753045 | 42862485 | 41490558 | G | T | intron_variant | 0 | 0 | 0 | 0 | 0 | 0 | 0.03333 | 0 | 0 | 0 |
| 503 | rs928871 | 42862531 | 41490604 | T | C | intron_variant | 0.07692 | 0.07692 | 0.4167 | 0.3333 | 0.3366 | 0.24 | 0.4778 | 0.3056 | 0.3611 | 0.175 |
| 504 | rs1266169667 | 42862605 | 41490678 | T | C | intron_variant | 0 | 0 | 0 | 0 | 0 | 0 | 0 | 0 | 0 | 0 |
| 505 | - | 42862676 | 41490749 | G | A | intron_variant | 0 | 0 | 0.02083 | 0 | 0 | 0 | 0 | 0 | 0 | 0 |
| 506 | rs370823400 | 42862693 | 41490766 | A | G | intron_variant | 0 | 0 | 0 | 0 | 0 | 0 | 0.06667 | 0 | 0 | 0 |
| 507 | rs116865960 | 42862768 | 41490841 | A | G | intron_variant | 0 | 0 | 0.02083 | 0 | 0.01485 | 0 | 0 | 0 | 0 | 0.025 |
| 508 | rs528452414 | 42862848 | 41490921 | C | A | intron_variant | 0 | 0 | 0 | 0.01282 | 0.009901 | 0 | 0 | 0 | 0 | 0 |
| 509 | rs115596471 | 42862869 | 41490942 | A | G | intron_variant | 0 | 0 | 0 | 0 | 0 | 0 | 0 | 0 | 0 | 0 |
| 510 | rs149173609 | 42862892 | 41490965 | T | C | intron_variant | 0 | 0 | 0 | 0 | 0 | 0 | 0 | 0 | 0.00463 | 0 |
| 511 | rs184365262 | 42862918 | 41490991 | T | C | intron_variant | 0 | 0 | 0 | 0 | 0.00495 | 0 | 0 | 0 | 0 | 0 |
| 512 | - | 42862924 | 41490997 | A | G | intron_variant | 0 | 0 | 0 | 0 | 0.00495 | 0 | 0 | 0 | 0.00463 | 0 |
| 513 | rs570544092,COSV59824452 | 42862927 | 41491000 | T | C | intron_variant | 0 | 0 | 0 | 0 | 0.00495 | 0 | 0 | 0 | 0.00463 | 0 |
| 514 | rs6517669 | 42862936 | 41491009 | G | A | intron_variant | 0.07692 | 0.07692 | 0.4167 | 0.3333 | 0.3366 | 0.24 | 0.4556 | 0.3056 | 0.3611 | 0.175 |
| 515 | rs1404996705 | 42862969 | 41491042 | C | T | intron_variant | 0 | 0 | 0 | 0 | 0.00495 | 0 | 0 | 0 | 0 | 0 |
| 516 | rs187208295 | 42863008 | 41491081 | C | T | intron_variant | 0 | 0 | 0 | 0 | 0 | 0 | 0 | 0 | 0 | 0.025 |
| 517 | rs117656646 | 42863050 | 41491123 | C | T | intron_variant | 0 | 0 | 0 | 0.01282 | 0.0297 | 0.02 | 0 | 0 | 0 | 0.025 |
| 518 | rs143270468 | 42863127 | 41491200 | A | G | intron_variant | 0 | 0 | 0 | 0 | 0.00495 | 0 | 0 | 0 | 0 | 0 |
| 519 | rs148304071 | 42863169 | 41491242 | T | C | intron_variant | 0 | 0 | 0 | 0 | 0 | 0 | 0 | 0 | 0 | 0 |
| 520 | - | 42863178 | 41491251 | A | G | intron_variant | 0.4231 | 0.4231 | 0.0625 | 0.03846 | 0.03465 | 0.12 | 0.1889 | 0.3611 | 0.3148 | 0 |
| 521 | rs140458174 | 42863219 | 41491292 | A | G | intron_variant | 0 | 0 | 0 | 0 | 0 | 0 | 0 | 0 | 0.009259 | 0 |
| 522 | - | 42863238 | 41491311 | G | C | intron_variant | 0 | 0 | 0 | 0 | 0 | 0 | 0 | 0 | 0 | 0.025 |
| 523 | rs374771570 | 42863251 | 41491324 | C | T | intron_variant | 0 | 0 | 0 | 0 | 0 | 0 | 0.01111 | 0 | 0 | 0 |
| 524 | rs866824112 | 42863293 | 41491366 | T | C | intron_variant | 0 | 0 | 0 | 0 | 0 | 0 | 0 | 0 | 0 | 0 |
| 525 | rs145570856 | 42863294 | 41491367 | A | G | intron_variant | 0 | 0 | 0 | 0 | 0 | 0 | 0 | 0 | 0 | 0 |
| 526 | rs35899679 | 42863320 | 41491393 | A | C | intron_variant | 0.07692 | 0.07692 | 0.2292 | 0.3846 | 0.3861 | 0.38 | 0.03333 | 0.05556 | 0.1667 | 0.475 |
| 527 | rs562045100 | 42863321 | 41491394 | A | G | intron_variant | 0 | 0 | 0.02083 | 0 | 0 | 0 | 0 | 0 | 0.02778 | 0 |
| 528 | - | 42863326 | 41491399 | A | G | intron_variant | 0.3462 | 0.3462 | 0.2917 | 0.4231 | 0.4208 | 0.5 | 0.2889 | 0.4167 | 0.4815 | 0.475 |
| 529 | rs35041537 | 42863369 | 41491442 | T | C | intron_variant | 0.07692 | 0.07692 | 0.2292 | 0.3974 | 0.3861 | 0.38 | 0.1 | 0.05556 | 0.1667 | 0.475 |

| | | | | | | | | | | | | | | | | |
|---|---|---|---|---|---|---|---|---|---|---|---|---|---|---|---|---|
| 530 | rs115429336 | 42863461 | 41491534 | A | G | intron_variant | 0 | 0 | 0 | 0 | 0 | 0 | 0 | 0 | 0 | 0 |
| 531 | rs1056841585 | 42863507 | 41491580 | A | T | intron_variant | 0 | 0 | 0 | 0 | 0 | 0 | 0 | 0 | 0.00463 | 0 |
| 532 | - | 42863608 | 41491681 | A | G | intron_variant | 0 | 0 | 0 | 0 | 0 | 0 | 0 | 0 | 0.00463 | 0 |
| 533 | - | 42863650 | 41491723 | C | T | intron_variant | 0 | 0 | 0 | 0 | 0 | 0 | 0 | 0 | 0.00463 | 0 |
| 534 | - | 42863685 | 41491758 | T | C | intron_variant | 0 | 0 | 0 | 0 | 0 | 0 | 0 | 0 | 0 | 0 |
| 535 | rs10154090 | 42863723 | 41491796 | A | T | intron_variant | 0.07692 | 0.07692 | 0.4167 | 0.3205 | 0.3218 | 0.18 | 0.4222 | 0.2778 | 0.338 | 0.1 |
| 536 | rs73357642 | 42863750 | 41491823 | A | G | intron_variant | 0 | 0 | 0 | 0 | 0 | 0.06 | 0.03333 | 0.02778 | 0 | 0.075 |
| 537 | rs391099 | 42863779 | 41491852 | C | T | intron_variant | 0.3077 | 0.3077 | 0.2917 | 0.2436 | 0.2574 | 0.32 | 0.2889 | 0.3056 | 0.1759 | 0.35 |
| 538 | rs73357644 | 42863827 | 41491900 | T | C | intron_variant | 0 | 0 | 0 | 0 | 0 | 0.06 | 0.04444 | 0.02778 | 0 | 0.075 |
| 539 | rs750532727 | 42863870 | 41491943 | C | T | intron_variant | 0 | 0 | 0 | 0 | 0 | 0 | 0 | 0 | 0.01389 | 0 |
| 540 | - | 42863922 | 41491995 | C | G | intron_variant | 0 | 0 | 0 | 0 | 0 | 0 | 0 | 0 | 0.00463 | 0 |
| 541 | - | 42863925 | 41491998 | A | C | intron_variant | 0 | 0 | 0 | 0 | 0 | 0 | 0 | 0 | 0.00463 | 0 |
| 542 | rs142659685 | 42863949 | 41492022 | T | C | intron_variant | 0 | 0 | 0 | 0.01282 | 0.00495 | 0 | 0.03333 | 0.02778 | 0.03241 | 0 |
| 543 | rs9305745 | 42864074 | 41492147 | T | C | intron_variant | 0.07692 | 0.07692 | 0.4167 | 0.3205 | 0.3218 | 0.18 | 0.4222 | 0.2778 | 0.3333 | 0.1 |
| 544 | rs886331524 | 42864155 | 41492228 | T | C | intron_variant | 0 | 0 | 0 | 0 | 0 | 0 | 0 | 0 | 0 | 0 |
| 545 | rs392370 | 42864166 | 41492239 | C | A | intron_variant | 0.2692 | 0.2692 | 0.2917 | 0.2436 | 0.2574 | 0.32 | 0.2889 | 0.3056 | 0.1852 | 0.35 |
| 546 | rs1311837486 | 42864175 | 41492248 | G | A | intron_variant | 0 | 0 | 0.02083 | 0 | 0 | 0 | 0 | 0 | 0 | 0 |
| 547 | rs79617378 | 42864217 | 41492290 | C | A | intron_variant | 0 | 0 | 0 | 0.01282 | 0.009901 | 0 | 0 | 0 | 0.00463 | 0.025 |
| 548 | rs771632181 | 42864269 | 41492342 | G | A | intron_variant | 0 | 0 | 0 | 0 | 0 | 0 | 0 | 0 | 0 | 0 |
| 549 | rs374261644 | 42864398 | 41492471 | C | T | intron_variant | 0 | 0 | 0 | 0 | 0 | 0 | 0 | 0 | 0 | 0 |
| 550 | rs886382727 | 42864457 | 41492530 | G | C | intron_variant | 0 | 0 | 0 | 0 | 0 | 0 | 0.01111 | 0 | 0 | 0 |
| 551 | rs746668285 | 42864515 | 41492588 | C | A | intron_variant | 0 | 0 | 0 | 0 | 0 | 0 | 0 | 0 | 0 | 0 |
| 552 | - | 42864520 | 41492593 | C | A | intron_variant | 0 | 0 | 0 | 0 | 0 | 0 | 0 | 0 | 0 | 0 |
| 553 | rs551122576 | 42864687 | 41492760 | A | G | intron_variant | 0 | 0 | 0 | 0 | 0 | 0 | 0 | 0 | 0 | 0 |
| 554 | rs2838040 | 42864692 | 41492765 | G | A | intron_variant | 0.07692 | 0.07692 | 0.4167 | 0.3205 | 0.3168 | 0.18 | 0.4889 | 0.2778 | 0.3333 | 0.125 |
| 555 | - | 42864743 | 41492816 | A | G | intron_variant | 0.2692 | 0.2692 | 0.3125 | 0.4487 | 0.4257 | 0.5 | 0.2556 | 0.4167 | 0.4907 | 0.475 |
| 556 | rs536023174 | 42864747 | 41492820 | T | C | intron_variant | 0 | 0 | 0 | 0 | 0 | 0.02 | 0 | 0 | 0 | 0 |
| 557 | rs401371 | 42864812 | 41492885 | C | G | intron_variant | 0.1923 | 0.1923 | 0.25 | 0.2179 | 0.2228 | 0.24 | 0.2444 | 0.2778 | 0.1574 | 0.225 |
| 558 | rs144988776 | 42864871 | 41492944 | T | C | intron_variant | 0 | 0 | 0 | 0 | 0.01485 | 0 | 0.06667 | 0 | 0 | 0 |
| 559 | - | 42864927 | 41493000 | A | G | intron_variant | 0 | 0 | 0 | 0 | 0 | 0 | 0 | 0 | 0 | 0.025 |
| 560 | rs373940448 | 42864947 | 41493020 | T | G | intron_variant | 0 | 0 | 0 | 0 | 0 | 0 | 0 | 0 | 0 | 0 |
| 561 | rs147138431 | 42865014 | 41493087 | A | C | intron_variant | 0 | 0 | 0 | 0.01282 | 0.009901 | 0 | 0 | 0 | 0 | 0 |
| 562 | rs368086767 | 42865035 | 41493108 | A | G | intron_variant | 0 | 0 | 0 | 0 | 0 | 0 | 0 | 0 | 0 | 0 |
| 563 | - | 42865047 | 41493120 | C | T | intron_variant | 0.3462 | 0.3462 | 0.1458 | 0.03846 | 0.04455 | 0.12 | 0.1889 | 0.5 | 0.4398 | 0 |
| 564 | - | 42865151 | 41493224 | T | C | intron_variant | 0.2692 | 0.2692 | 0.5 | 0.3718 | 0.3465 | 0.34 | 0.3444 | 0.3611 | 0.3426 | 0.125 |
| 565 | rs375827195 | 42865216 | 41493289 | G | A | intron_variant | 0 | 0 | 0 | 0 | 0 | 0 | 0.01111 | 0 | 0 | 0.05 |
| 566 | rs186185630 | 42865275 | 41493348 | T | C | intron_variant | 0 | 0 | 0 | 0 | 0 | 0 | 0 | 0 | 0 | 0 |
| 567 | - | 42865283 | 41493356 | G | A | intron_variant | 0 | 0 | 0 | 0 | 0.00495 | 0.02 | 0 | 0 | 0 | 0 |
| 568 | - | 42865300 | 41493373 | G | A | intron_variant | 0 | 0 | 0 | 0 | 0.00495 | 0 | 0 | 0 | 0 | 0 |
| 569 | - | 42865324 | 41493397 | G | A | intron_variant | 0.1538 | 0.1538 | 0.3542 | 0.2821 | 0.2871 | 0.46 | 0.4444 | 0.3333 | 0.4815 | 0.35 |
| 570 | rs1180926833 | 42865398 | 41493471 | C | T | intron_variant | 0 | 0 | 0 | 0 | 0 | 0 | 0 | 0 | 0 | 0 |
| 571 | rs377011212 | 42865555 | 41493628 | T | C | intron_variant | 0 | 0 | 0 | 0 | 0 | 0 | 0.02222 | 0 | 0 | 0 |
| 572 | rs566903241 | 42865789 | 41493862 | C | T | intron_variant | 0 | 0 | 0 | 0 | 0 | 0 | 0 | 0 | 0 | 0 |
| 573 | rs549801931 | 42865820 | 41493893 | T | C | intron_variant | 0 | 0 | 0 | 0 | 0 | 0 | 0 | 0 | 0 | 0 |
| 574 | rs1601585975 | 42865828 | 41493901 | G | C | intron_variant | 0 | 0 | 0 | 0 | 0 | 0 | 0 | 0 | 0 | 0 |
| 575 | rs2838041 | 42865906 | 41493979 | C | G | intron_variant | 0 | 0 | 0.08333 | 0.07692 | 0.0198 | 0.04 | 0 | 0 | 0.009259 | 0.05 |
| 576 | rs1000568527 | 42866068 | 41494141 | G | C | intron_variant | 0 | 0 | 0 | 0 | 0 | 0 | 0 | 0 | 0 | 0 |
| 577 | rs376235035 | 42866099 | 41494172 | T | C | intron_variant | 0 | 0 | 0 | 0 | 0.0198 | 0 | 0 | 0 | 0.0463 | 0 |
| 578 | rs2838042 | 42866107 | 41494180 | C | T | intron_variant | 0.07692 | 0.07692 | 0.3333 | 0.2051 | 0.2327 | 0.18 | 0.4444 | 0.25 | 0.1343 | 0.25 |
| 579 | rs923545665 | 42866226 | 41494299 | C | G | intron_variant | 0 | 0 | 0 | 0 | 0 | 0 | 0.01111 | 0 | 0 | 0 |
| 580 | rs373196115 | 42866232 | 41494305 | A | G | intron_variant | 0 | 0 | 0 | 0 | 0 | 0 | 0 | 0.05556 | 0 | 0 |
| 581 | rs750152865 | 42866245 | 41494318 | A | G | intron_variant | 0 | 0 | 0 | 0 | 0 | 0 | 0 | 0 | 0 | 0 |
| 582 | rs3787950,COSV59821982 | 42866296 | 41494369 | C | T | synonymous_variant | 0 | 0 | 0.125 | 0.1538 | 0.06931 | 0.22 | 0.4 | 0.2222 | 0.06481 | 0.175 |

| | | | | | | | | | | | | | | | | |
|---|---|---|---|---|---|---|---|---|---|---|---|---|---|---|---|---|
| 583 | rs61735792,COSV59824543 | 42866332 | 41494405 | A | G | synonymous_variant | 0 | 0 | 0 | 0.01282 | 0.009901 | 0 | 0 | 0 | 0 | 0 |
| 584 | rs201679623,COSV59823003 | 42866388 | 41494461 | C | A | missense_variant | 0 | 0 | 0 | 0.01282 | 0 | 0 | 0 | 0 | 0.00463 | 0 |
| 585 | rs138651919 | 42866399 | 41494472 | A | G | missense_variant | 0 | 0 | 0 | 0 | 0 | 0 | 0 | 0 | 0 | 0 |
| 586 | rs199824558,COSV59820898 | 42866422 | 41494495 | A | G | synonymous_variant | 0 | 0 | 0 | 0 | 0 | 0.02 | 0 | 0 | 0 | 0 |
| 587 | rs201093031 | 42866423 | 41494496 | G | A | missense_variant | 0 | 0 | 0 | 0 | 0 | 0 | 0.01111 | 0 | 0 | 0 |
| 588 | rs61735791,COSV59824252 | 42866439 | 41494512 | T | C | missense_variant | 0 | 0 | 0 | 0 | 0.00495 | 0 | 0 | 0 | 0 | 0.025 |
| 589 | rs141685390 | 42866440 | 41494513 | C | G | synonymous_variant | 0 | 0 | 0 | 0 | 0 | 0 | 0.01111 | 0 | 0 | 0 |
| 590 | rs61735790 | 42866468 | 41494541 | C | T | missense_variant | 0 | 0 | 0 | 0 | 0 | 0 | 0 | 0 | 0 | 0 |
| 591 | rs368773451 | 42866553 | 41494626 | C | T | intron_variant | 0 | 0 | 0 | 0 | 0 | 0 | 0 | 0 | 0 | 0 |
| 592 | rs368625065 | 42866644 | 41494717 | A | G | intron_variant | 0 | 0 | 0 | 0 | 0 | 0 | 0 | 0 | 0 | 0 |
| 593 | rs547701911 | 42866723 | 41494796 | C | T | intron_variant | 0 | 0 | 0 | 0 | 0.00495 | 0 | 0 | 0 | 0 | 0 |
| 594 | - | 42866725 | 41494798 | T | C | intron_variant | 0 | 0 | 0 | 0 | 0 | 0 | 0 | 0 | 0 | 0 |
| 595 | rs374203194 | 42866726 | 41494799 | A | G | intron_variant | 0 | 0 | 0 | 0.01282 | 0 | 0 | 0 | 0 | 0 | 0 |
| 596 | rs569947342 | 42866868 | 41494941 | T | C | intron_variant | 0 | 0 | 0 | 0 | 0 | 0 | 0 | 0 | 0.00463 | 0 |
| 597 | - | 42866884 | 41494957 | A | T | intron_variant | 0.03846 | 0.03846 | 0 | 0 | 0 | 0 | 0 | 0 | 0 | 0 |
| 598 | rs927713682,COSV59825745 | 42866888 | 41494961 | T | C | intron_variant | 0 | 0 | 0 | 0 | 0 | 0 | 0 | 0 | 0.009259 | 0 |
| 599 | - | 42866891 | 41494964 | A | G | intron_variant | 0 | 0 | 0 | 0 | 0 | 0 | 0 | 0.02778 | 0 | 0 |
| 600 | rs1007596550 | 42866974 | 41495047 | A | G | intron_variant | 0 | 0 | 0 | 0 | 0 | 0 | 0.03333 | 0 | 0 | 0 |
| 601 | rs190899605,COSV59822774 | 42867015 | 41495088 | C | T | intron_variant | 0 | 0 | 0.02083 | 0 | 0 | 0.02 | 0 | 0 | 0 | 0 |
| 602 | - | 42867175 | 41495248 | A | T | intron_variant | 0 | 0 | 0 | 0 | 0 | 0 | 0 | 0 | 0.00463 | 0 |
| 603 | - | 42867179 | 41495252 | T | C | intron_variant | 0.03846 | 0.03846 | 0 | 0 | 0 | 0 | 0 | 0 | 0 | 0 |
| 604 | rs1307770905 | 42867237 | 41495310 | G | T | intron_variant | 0 | 0 | 0 | 0 | 0 | 0 | 0 | 0.02778 | 0 | 0 |
| 605 | rs558991814 | 42867326 | 41495399 | C | G | intron_variant | 0 | 0 | 0 | 0 | 0.00495 | 0 | 0 | 0 | 0 | 0 |
| 606 | rs530471976 | 42867424 | 41495497 | T | C | intron_variant | 0 | 0 | 0 | 0 | 0 | 0 | 0 | 0 | 0 | 0 |
| 607 | rs114911304 | 42867425 | 41495498 | A | G | intron_variant | 0 | 0 | 0 | 0 | 0 | 0 | 0 | 0 | 0 | 0 |
| 608 | rs775158277 | 42867499 | 41495572 | C | T | intron_variant | 0 | 0 | 0 | 0.02564 | 0 | 0 | 0 | 0 | 0 | 0.025 |
| 609 | rs58146697,COSV59822780 | 42867667 | 41495740 | C | T | intron_variant | 0 | 0 | 0.02083 | 0.01282 | 0.02475 | 0.06 | 0.01111 | 0.05556 | 0.03704 | 0 |
| 610 | - | 42867719 | 41495792 | T | C | intron_variant | 0 | 0 | 0 | 0 | 0 | 0 | 0.01111 | 0 | 0 | 0 |
| 611 | rs372453344 | 42867735 | 41495808 | T | C | intron_variant | 0 | 0 | 0 | 0 | 0 | 0 | 0 | 0 | 0 | 0 |
| 612 | rs142425263 | 42867756 | 41495829 | A | C | intron_variant | 0 | 0 | 0 | 0 | 0 | 0 | 0 | 0 | 0.01389 | 0 |
| 613 | rs184876485 | 42867990 | 41496063 | G | A | intron_variant | 0 | 0 | 0 | 0.01282 | 0 | 0 | 0 | 0 | 0 | 0 |
| 614 | rs370530303 | 42868060 | 41496133 | T | C | intron_variant | 0 | 0 | 0 | 0 | 0 | 0 | 0.04444 | 0 | 0 | 0 |
| 615 | rs55760462 | 42868206 | 41496279 | C | T | intron_variant | 0.03846 | 0.03846 | 0.2083 | 0.2179 | 0.1881 | 0.06 | 0.04444 | 0.02778 | 0.1296 | 0.15 |
| 616 | - | 42868299 | 41496372 | G | A | intron_variant | 0 | 0 | 0 | 0 | 0 | 0 | 0.01111 | 0 | 0 | 0 |
| 617 | rs144276163 | 42868411 | 41496484 | G | T | intron_variant | 0 | 0 | 0 | 0 | 0 | 0 | 0 | 0 | 0 | 0 |
| 618 | rs914184 | 42868641 | 41496714 | A | G | intron_variant | 0 | 0 | 0.02083 | 0 | 0 | 0 | 0.01111 | 0 | 0.009259 | 0 |
| 619 | rs78217567,COSV59821242 | 42868655 | 41496728 | C | T | intron_variant | 0 | 0 | 0.04167 | 0 | 0.04455 | 0.02 | 0 | 0 | 0.00463 | 0 |
| 620 | - | 42868820 | 41496893 | C | T | intron_variant | 0 | 0 | 0 | 0 | 0 | 0 | 0 | 0 | 0 | 0 |
| 621 | rs34256269 | 42868876 | 41496949 | A | G | intron_variant | 0 | 0 | 0.1667 | 0.1026 | 0.1485 | 0.18 | 0.1111 | 0.05556 | 0.0463 | 0.1 |
| 622 | COSV59828093 | 42868888 | 41496961 | C | T | intron_variant | 0.3462 | 0.3462 | 0.3542 | 0.4487 | 0.495 | 0.34 | 0.4778 | 0.4444 | 0.4398 | 0.4 |
| 623 | - | 42868905 | 41496978 | C | T | intron_variant | 0 | 0 | 0 | 0 | 0 | 0 | 0.01111 | 0 | 0 | 0 |
| 624 | rs573213706 | 42868957 | 41497030 | A | G | intron_variant | 0 | 0 | 0 | 0 | 0.00495 | 0 | 0 | 0 | 0 | 0 |
| 625 | - | 42868992 | 41497065 | C | T | intron_variant | 0 | 0 | 0 | 0 | 0.009901 | 0 | 0 | 0 | 0 | 0 |
| 626 | rs34983238,COSV59821615 | 42868997 | 41497070 | C | A | intron_variant | 0 | 0 | 0.08333 | 0.1538 | 0.07426 | 0.08 | 0 | 0 | 0.01389 | 0.25 |
| 627 | - | 42869028 | 41497101 | G | A | intron_variant | 0 | 0 | 0 | 0 | 0 | 0 | 0 | 0 | 0.00463 | 0 |
| 628 | - | 42869063 | 41497136 | A | G | intron_variant | 0.3462 | 0.3462 | 0.1875 | 0.3462 | 0.3564 | 0.16 | 0.3889 | 0.3889 | 0.3889 | 0.3 |
| 629 | - | 42869080 | 41497153 | C | T | intron_variant | 0 | 0 | 0 | 0 | 0.00495 | 0 | 0 | 0 | 0 | 0 |
| 630 | rs924332623 | 42869283 | 41497356 | A | C | intron_variant | 0 | 0 | 0 | 0 | 0 | 0 | 0 | 0 | 0 | 0 |
| 631 | rs1003030,COSV59820931 | 42869606 | 41497679 | G | A | intron_variant | 0 | 0 | 0.25 | 0.141 | 0.1188 | 0.24 | 0.2333 | 0.2222 | 0.1389 | 0.2 |
| 632 | rs184859933 | 42869637 | 41497710 | T | C | intron_variant | 0 | 0 | 0 | 0 | 0 | 0 | 0 | 0 | 0 | 0 |
| 633 | rs56695953 | 42869735 | 41497808 | A | G | intron_variant | 0.2308 | 0.2308 | 0.02083 | 0.1923 | 0.1386 | 0.04 | 0.03333 | 0.02778 | 0.05556 | 0.175 |
| 634 | rs148901354 | 42869772 | 41497845 | T | C | intron_variant | 0 | 0 | 0 | 0.01282 | 0 | 0 | 0 | 0 | 0 | 0 |
| 635 | rs367885466 | 42869836 | 41497909 | T | C | intron_variant | 0 | 0 | 0 | 0 | 0 | 0 | 0 | 0 | 0 | 0 |

| | | | | | | | | | | | | | | | | |
|---|---|---|---|---|---|---|---|---|---|---|---|---|---|---|---|---|
| 636 | rs181592444 | 42869927 | 41498000 | T | C | intron_variant | 0 | 0 | 0 | 0 | 0 | 0 | 0.01111 | 0.02778 | 0 | 0 |
| 637 | rs115968373 | 42869938 | 41498011 | A | G | intron_variant | 0 | 0 | 0 | 0 | 0 | 0 | 0 | 0 | 0 | 0 |
| 638 | rs186510586 | 42870205 | 41498278 | C | T | intron_variant | 0 | 0 | 0 | 0.01282 | 0 | 0 | 0 | 0 | 0 | 0 |
| 639 | - | 42870211 | 41498284 | C | T | intron_variant | 0 | 0 | 0 | 0 | 0 | 0 | 0 | 0 | 0 | 0 |
| 640 | rs11701576,COSV59820936 | 42870262 | 41498335 | G | A | intron_variant | 0 | 0 | 0.25 | 0.141 | 0.1188 | 0.24 | 0.2333 | 0.2222 | 0.1389 | 0.2 |
| 641 | rs780560155 | 42870392 | 41498465 | A | G | intron_variant | 0 | 0 | 0 | 0.01282 | 0 | 0 | 0 | 0 | 0 | 0 |
| 642 | rs138661151 | 42870421 | 41498494 | A | G | intron_variant | 0 | 0 | 0 | 0 | 0 | 0 | 0 | 0 | 0 | 0 |
| 643 | rs150715708 | 42870450 | 41498523 | T | C | intron_variant | 0 | 0 | 0 | 0.01282 | 0 | 0 | 0 | 0 | 0.00463 | 0 |
| 644 | rs557714443 | 42870451 | 41498524 | A | G | intron_variant | 0 | 0 | 0 | 0 | 0 | 0 | 0.01111 | 0 | 0 | 0 |
| 645 | rs3761373,COSV59820941 | 42870522 | 41498595 | T | C | intron_variant | 0 | 0 | 0.25 | 0.141 | 0.1188 | 0.24 | 0.2333 | 0.2222 | 0.1389 | 0.2 |
| 646 | rs940625478 | 42870890 | 41498963 | G | T | intron_variant | 0 | 0 | 0 | 0 | 0.00495 | 0 | 0 | 0 | 0 | 0 |
| 647 | rs913523407 | 42870902 | 41498975 | A | G | intron_variant | 0 | 0 | 0.02083 | 0 | 0 | 0 | 0 | 0 | 0 | 0 |
| 648 | rs422761,COSV59821427 | 42870993 | 41499066 | A | G | intron_variant | 0.3846 | 0.3846 | 0.08333 | 0.02564 | 0.06931 | 0.26 | 0.2 | 0.3056 | 0.2778 | 0.075 |
| 649 | - | 42871051 | 41499124 | G | A | intron_variant | 0 | 0 | 0 | 0 | 0 | 0 | 0 | 0 | 0 | 0 |
| 650 | rs1233077154 | 42871073 | 41499146 | T | C | intron_variant | 0 | 0 | 0 | 0 | 0 | 0 | 0 | 0 | 0 | 0 |
| 651 | rs939814881 | 42871083 | 41499156 | T | C | intron_variant | 0 | 0 | 0 | 0 | 0.00495 | 0 | 0 | 0 | 0 | 0 |
| 652 | rs389001,COSV59821435 | 42871131 | 41499204 | G | A | intron_variant | 0.3846 | 0.3846 | 0.08333 | 0.02564 | 0.06931 | 0.26 | 0.2 | 0.3056 | 0.2778 | 0.075 |
| 653 | rs931406088 | 42871153 | 41499226 | T | C | intron_variant | 0 | 0 | 0 | 0 | 0 | 0 | 0 | 0 | 0 | 0 |
| 654 | rs534034788 | 42871213 | 41499286 | A | G | intron_variant | 0 | 0 | 0 | 0 | 0 | 0.02 | 0 | 0 | 0 | 0 |
| 655 | rs2838043 | 42871220 | 41499293 | T | C | intron_variant | 0.2308 | 0.2308 | 0.02083 | 0.1923 | 0.1386 | 0.04 | 0.03333 | 0.02778 | 0.05556 | 0.175 |
| 656 | rs187648498 | 42871225 | 41499298 | C | T | intron_variant | 0 | 0 | 0 | 0 | 0 | 0 | 0 | 0 | 0 | 0 |
| 657 | rs372249693 | 42871227 | 41499300 | T | C | intron_variant | 0 | 0 | 0 | 0 | 0 | 0 | 0 | 0 | 0 | 0 |
| 658 | rs149424945 | 42871251 | 41499324 | T | C | intron_variant | 0 | 0 | 0 | 0 | 0 | 0 | 0 | 0.02778 | 0 | 0 |
| 659 | rs3761374,COSV59821249 | 42871273 | 41499346 | C | T | intron_variant | 0 | 0 | 0.1667 | 0.1026 | 0.1485 | 0.18 | 0.1222 | 0.05556 | 0.0463 | 0.1 |
| 660 | rs1418216524 | 42871415 | 41499488 | G | A | intron_variant | 0 | 0 | 0 | 0 | 0 | 0 | 0 | 0 | 0.00463 | 0 |
| 661 | rs190673599 | 42871481 | 41499554 | T | A | intron_variant | 0 | 0 | 0 | 0 | 0 | 0 | 0 | 0 | 0 | 0 |
| 662 | rs183722985 | 42871662 | 41499735 | A | G | intron_variant | 0 | 0 | 0 | 0 | 0 | 0 | 0 | 0 | 0 | 0 |
| 663 | rs573044175 | 42871680 | 41499753 | A | G | intron_variant | 0 | 0 | 0 | 0 | 0 | 0 | 0 | 0 | 0 | 0 |
| 664 | - | 42871776 | 41499849 | C | T | intron_variant | 0.03846 | 0.03846 | 0 | 0 | 0 | 0 | 0 | 0 | 0 | 0 |
| 665 | rs188198121 | 42871792 | 41499865 | A | G | intron_variant | 0 | 0 | 0 | 0 | 0 | 0 | 0 | 0 | 0 | 0 |
| 666 | rs571275590 | 42871867 | 41499940 | G | A | intron_variant | 0 | 0 | 0 | 0 | 0 | 0.02 | 0 | 0 | 0 | 0 |
| 667 | rs78528160 | 42871921 | 41499994 | A | T | intron_variant | 0 | 0 | 0 | 0 | 0 | 0 | 0 | 0 | 0 | 0 |
| 668 | - | 42871941 | 41500014 | G | C | intron_variant | 0.1154 | 0.1154 | 0.1667 | 0.1538 | 0.2178 | 0.14 | 0.3556 | 0.3611 | 0.338 | 0.125 |
| 669 | rs144318842 | 42872059 | 41500132 | A | G | intron_variant | 0 | 0 | 0 | 0 | 0 | 0 | 0 | 0 | 0 | 0 |
| 670 | rs556371948 | 42872119 | 41500192 | T | C | intron_variant | 0 | 0 | 0 | 0 | 0 | 0 | 0 | 0 | 0 | 0.025 |
| 671 | rs374881231 | 42872302 | 41500375 | A | G | intron_variant | 0 | 0 | 0 | 0 | 0 | 0 | 0 | 0 | 0 | 0 |
| 672 | rs148719900 | 42872315 | 41500388 | A | C | intron_variant | 0 | 0 | 0 | 0 | 0 | 0 | 0 | 0 | 0 | 0 |
| 673 | rs142296178 | 42872342 | 41500415 | A | G | intron_variant | 0 | 0 | 0 | 0 | 0 | 0 | 0 | 0 | 0 | 0 |
| 674 | rs527416684 | 42872354 | 41500427 | A | G | intron_variant | 0 | 0 | 0 | 0 | 0 | 0.04 | 0 | 0 | 0 | 0 |
| 675 | - | 42872372 | 41500445 | A | G | intron_variant | 0 | 0 | 0 | 0 | 0 | 0 | 0 | 0 | 0.00463 | 0 |
| 676 | rs986686535 | 42872391 | 41500464 | T | C | intron_variant | 0 | 0 | 0 | 0 | 0 | 0 | 0.01111 | 0 | 0 | 0 |
| 677 | rs151248079 | 42872429 | 41500502 | T | A | intron_variant | 0 | 0 | 0 | 0 | 0 | 0 | 0 | 0 | 0 | 0 |
| 678 | rs115266855 | 42872559 | 41500632 | G | A | intron_variant | 0 | 0 | 0 | 0 | 0 | 0 | 0 | 0 | 0 | 0 |
| 679 | rs114027875 | 42872658 | 41500731 | T | A | intron_variant | 0 | 0 | 0 | 0 | 0 | 0 | 0 | 0 | 0 | 0 |
| 680 | rs2187238 | 42872751 | 41500824 | C | T | intron_variant | 0.2308 | 0.2308 | 0.02083 | 0.1923 | 0.1386 | 0.06 | 0.03333 | 0.02778 | 0.05556 | 0.175 |
| 681 | rs112416131 | 42872793 | 41500866 | A | G | intron_variant | 0 | 0 | 0 | 0 | 0.00495 | 0 | 0 | 0 | 0 | 0 |
| 682 | - | 42872897 | 41500970 | G | C | intron_variant | 0.1154 | 0.1154 | 0.1667 | 0.1538 | 0.2178 | 0.14 | 0.3556 | 0.3611 | 0.338 | 0.125 |
| 683 | rs1343871927 | 42872949 | 41501022 | A | G | intron_variant | 0.03846 | 0.03846 | 0 | 0 | 0 | 0 | 0 | 0 | 0 | 0 |
| 684 | rs73903404 | 42872955 | 41501028 | G | A | intron_variant | 0 | 0 | 0 | 0 | 0 | 0 | 0 | 0 | 0 | 0 |
| 685 | - | 42872956 | 41501029 | C | T | intron_variant | 0 | 0 | 0 | 0 | 0 | 0 | 0 | 0.02778 | 0 | 0 |
| 686 | - | 42873008 | 41501081 | C | T | intron_variant | 0 | 0 | 0 | 0 | 0 | 0 | 0.01111 | 0 | 0 | 0 |
| 687 | rs972527688 | 42873213 | 41501286 | A | T | intron_variant | 0 | 0 | 0 | 0 | 0 | 0 | 0 | 0 | 0 | 0 |
| 688 | rs530009764 | 42873243 | 41501316 | C | T | intron_variant | 0 | 0 | 0 | 0 | 0.00495 | 0 | 0 | 0 | 0 | 0 |

| | | | | | | | | | | | | | | | | |
|---|---|---|---|---|---|---|---|---|---|---|---|---|---|---|---|---|
| 689 | rs371046741 | 42873258 | 41501331 | A | G | intron_variant | 0 | 0 | 0 | 0 | 0 | 0 | 0 | 0 | 0 | 0 |
| 690 | rs147620204 | 42873274 | 41501347 | C | G | intron_variant | 0 | 0 | 0 | 0 | 0 | 0 | 0 | 0 | 0 | 0 |
| 691 | rs552447029 | 42873362 | 41501435 | G | C | intron_variant | 0 | 0 | 0 | 0 | 0 | 0 | 0 | 0 | 0 | 0 |
| 692 | rs1003304387 | 42873400 | 41501473 | G | A | intron_variant | 0 | 0 | 0 | 0 | 0 | 0 | 0 | 0 | 0 | 0 |
| 693 | rs962596107 | 42873604 | 41501677 | C | G | intron_variant | 0 | 0 | 0 | 0 | 0 | 0 | 0 | 0 | 0 | 0 |
| 694 | rs9980693,COSV59821257 | 42873605 | 41501678 | A | G | intron_variant | 0 | 0 | 0.1667 | 0.1026 | 0.1535 | 0.16 | 0.1222 | 0.05556 | 0.0463 | 0.1 |
| 695 | rs73903405 | 42873700 | 41501773 | A | C | intron_variant | 0 | 0 | 0 | 0 | 0 | 0 | 0 | 0 | 0 | 0 |
| 696 | rs145877432 | 42873827 | 41501900 | A | G | intron_variant | 0 | 0 | 0 | 0 | 0 | 0 | 0 | 0 | 0 | 0 |
| 697 | - | 42873984 | 41502057 | C | T | intron_variant | 0 | 0 | 0 | 0 | 0 | 0 | 0 | 0 | 0 | 0 |
| 698 | - | 42874065 | 41502138 | A | G | intron_variant | 0 | 0 | 0.02083 | 0 | 0 | 0 | 0 | 0 | 0 | 0 |
| 699 | rs968067353 | 42874098 | 41502171 | A | G | intron_variant | 0 | 0 | 0 | 0 | 0 | 0 | 0 | 0.02778 | 0 | 0 |
| 700 | rs377403765 | 42874104 | 41502177 | A | G | intron_variant | 0 | 0 | 0 | 0 | 0 | 0 | 0 | 0 | 0 | 0 |
| 701 | rs149270377 | 42874184 | 41502257 | A | G | intron_variant | 0 | 0 | 0 | 0 | 0 | 0 | 0 | 0 | 0 | 0 |
| 702 | rs143460343 | 42874270 | 41502343 | A | G | intron_variant | 0 | 0 | 0 | 0 | 0 | 0 | 0 | 0 | 0 | 0 |
| 703 | rs141583878 | 42874306 | 41502379 | A | G | intron_variant | 0 | 0 | 0 | 0.01282 | 0 | 0 | 0 | 0 | 0 | 0 |
| 704 | rs567302726 | 42874338 | 41502411 | A | G | intron_variant | 0 | 0 | 0 | 0 | 0 | 0 | 0 | 0 | 0 | 0 |
| 705 | - | 42874368 | 41502441 | G | A | intron_variant | 0.03846 | 0.03846 | 0 | 0 | 0.00495 | 0 | 0 | 0 | 0 | 0 |
| 706 | - | 42874384 | 41502457 | G | A | intron_variant | 0 | 0 | 0 | 0 | 0 | 0 | 0 | 0 | 0 | 0 |
| 707 | rs536147878 | 42874513 | 41502586 | A | G | intron_variant | 0 | 0 | 0 | 0 | 0.00495 | 0 | 0 | 0 | 0 | 0 |
| 708 | rs992437148 | 42874535 | 41502608 | T | C | intron_variant | 0 | 0 | 0 | 0 | 0 | 0 | 0 | 0 | 0.00463 | 0 |
| 709 | rs146385718 | 42874559 | 41502632 | C | T | intron_variant | 0 | 0 | 0 | 0 | 0 | 0 | 0 | 0 | 0 | 0 |
| 710 | rs531756387 | 42874831 | 41502904 | T | G | intron_variant | 0 | 0 | 0.02083 | 0 | 0 | 0 | 0.01111 | 0 | 0 | 0 |
| 711 | rs139700775,COSV59826804 | 42874841 | 41502914 | C | T | intron_variant | 0 | 0 | 0 | 0 | 0 | 0 | 0 | 0 | 0.00463 | 0 |
| 712 | - | 42874844 | 41502917 | T | C | intron_variant | 0 | 0 | 0 | 0 | 0 | 0 | 0.01111 | 0 | 0 | 0 |
| 713 | rs9984012,COSV59821265 | 42875004 | 41503077 | T | C | intron_variant | 0 | 0 | 0.1667 | 0.1026 | 0.1535 | 0.16 | 0.1222 | 0.05556 | 0.0463 | 0.1 |
| 714 | rs142914234 | 42875089 | 41503162 | C | T | intron_variant | 0 | 0 | 0 | 0 | 0 | 0 | 0 | 0 | 0 | 0 |
| 715 | rs1308693805 | 42875131 | 41503204 | T | C | intron_variant | 0 | 0 | 0 | 0 | 0 | 0 | 0 | 0 | 0.00463 | 0 |
| 716 | - | 42875190 | 41503263 | A | G | intron_variant | 0 | 0 | 0 | 0 | 0.00495 | 0 | 0 | 0 | 0 | 0 |
| 717 | rs420737,COSV59821448 | 42875404 | 41503477 | G | A | intron_variant | 0.3846 | 0.3846 | 0.08333 | 0.02564 | 0.06931 | 0.26 | 0.2444 | 0.2778 | 0.2778 | 0.075 |
| 718 | rs146120690 | 42875620 | 41503693 | A | G | intron_variant | 0 | 0 | 0 | 0 | 0 | 0 | 0.08889 | 0 | 0 | 0 |
| 719 | rs74749793,COSV59820948 | 42875652 | 41503725 | T | G | intron_variant | 0 | 0 | 0.25 | 0.141 | 0.1188 | 0.24 | 0.1889 | 0.1944 | 0.1389 | 0.2 |
| 720 | rs1033407344 | 42875668 | 41503741 | A | C | intron_variant | 0 | 0 | 0 | 0 | 0 | 0 | 0 | 0 | 0 | 0 |
| 721 | rs373052874 | 42875816 | 41503889 | C | G | intron_variant | 0 | 0 | 0 | 0 | 0 | 0 | 0.05556 | 0 | 0 | 0 |
| 722 | rs763295257 | 42875832 | 41503905 | A | G | intron_variant | 0 | 0 | 0 | 0 | 0.009901 | 0 | 0 | 0 | 0.009259 | 0 |
| 723 | - | 42875847 | 41503920 | T | G | intron_variant | 0 | 0 | 0 | 0 | 0 | 0 | 0 | 0 | 0 | 0 |
| 724 | rs1381741328 | 42875859 | 41503932 | T | C | intron_variant | 0 | 0 | 0 | 0 | 0.00495 | 0 | 0 | 0 | 0 | 0 |
| 725 | - | 42876060 | 41504133 | C | T | intron_variant | 0 | 0 | 0 | 0 | 0 | 0 | 0 | 0 | 0 | 0.025 |
| 726 | rs575313753 | 42876148 | 41504221 | G | A | intron_variant | 0 | 0 | 0 | 0 | 0 | 0 | 0 | 0 | 0 | 0 |
| 727 | - | 42876162 | 41504235 | C | T | intron_variant | 0 | 0 | 0 | 0 | 0 | 0 | 0 | 0 | 0 | 0.025 |
| 728 | rs7277080 | 42876400 | 41504473 | T | C | intron_variant | 0.03846 | 0.03846 | 0.3125 | 0.3846 | 0.3069 | 0.14 | 0.04444 | 0.02778 | 0.1435 | 0.325 |
| 729 | rs756657861 | 42876412 | 41504485 | A | G | intron_variant | 0 | 0 | 0 | 0 | 0.009901 | 0 | 0 | 0 | 0.009259 | 0 |
| 730 | rs395584,COSV59821459 | 42876447 | 41504520 | C | T | intron_variant | 0.3846 | 0.3846 | 0.08333 | 0.02564 | 0.06931 | 0.26 | 0.2444 | 0.2778 | 0.2778 | 0.075 |
| 731 | rs8129713 | 42876598 | 41504671 | C | T | intron_variant | 0.2308 | 0.2308 | 0.02083 | 0.1923 | 0.1337 | 0.06 | 0.03333 | 0.05556 | 0.05556 | 0.175 |
| 732 | - | 42876599 | 41504672 | T | C | intron_variant | 0 | 0 | 0 | 0 | 0 | 0.02 | 0 | 0 | 0 | 0 |
| 733 | - | 42876684 | 41504757 | C | G | intron_variant | 0 | 0 | 0 | 0 | 0 | 0 | 0 | 0 | 0 | 0 |
| 734 | rs1027558002 | 42876698 | 41504771 | A | G | intron_variant | 0 | 0 | 0 | 0 | 0 | 0 | 0 | 0 | 0 | 0 |
| 735 | rs1014761397 | 42876720 | 41504793 | T | C | intron_variant | 0 | 0 | 0 | 0 | 0 | 0 | 0 | 0 | 0 | 0 |
| 736 | - | 42876757 | 41504830 | T | C | intron_variant | 0 | 0 | 0 | 0 | 0 | 0 | 0 | 0 | 0 | 0 |
| 737 | rs374886738 | 42876786 | 41504859 | C | A | intron_variant | 0 | 0 | 0 | 0 | 0 | 0 | 0 | 0 | 0 | 0 |
| 738 | rs368411333 | 42876791 | 41504864 | T | A | intron_variant | 0 | 0 | 0 | 0 | 0 | 0 | 0 | 0 | 0 | 0 |
| 739 | rs73357663 | 42876845 | 41504918 | C | T | intron_variant | 0 | 0 | 0 | 0 | 0 | 0 | 0 | 0 | 0 | 0 |
| 740 | rs1045511841 | 42876921 | 41504994 | A | G | intron_variant | 0 | 0 | 0 | 0 | 0 | 0 | 0 | 0 | 0 | 0.025 |
| 741 | - | 42876949 | 41505022 | T | C | intron_variant | 0 | 0 | 0 | 0 | 0 | 0 | 0 | 0 | 0.00463 | 0 |

| | | | | | | | | | | | | | | | | |
|---|---|---|---|---|---|---|---|---|---|---|---|---|---|---|---|---|
| 742 | rs145728087 | 42877028 | 41505101 | A | C | intron_variant | 0 | 0 | 0 | 0 | 0 | 0 | 0 | 0 | 0 | 0 |
| 743 | rs148499598 | 42877041 | 41505114 | C | T | intron_variant | 0 | 0 | 0 | 0 | 0.009901 | 0 | 0 | 0 | 0.00463 | 0 |
| 744 | - | 42877167 | 41505240 | C | A | intron_variant | 0 | 0 | 0 | 0 | 0 | 0.02 | 0 | 0 | 0 | 0 |
| 745 | rs186418926 | 42877262 | 41505335 | A | G | intron_variant | 0 | 0 | 0.02083 | 0 | 0.009901 | 0 | 0 | 0 | 0 | 0 |
| 746 | rs74605993 | 42877274 | 41505347 | T | C | intron_variant | 0 | 0 | 0 | 0 | 0 | 0 | 0 | 0 | 0 | 0 |
| 747 | - | 42877307 | 41505380 | G | C | intron_variant | 0 | 0 | 0 | 0 | 0 | 0 | 0.01111 | 0 | 0 | 0 |
| 748 | rs75373173,COSV59822963 | 42877382 | 41505455 | C | T | intron_variant | 0 | 0 | 0.04167 | 0.07692 | 0.1584 | 0.04 | 0 | 0 | 0.02315 | 0.025 |
| 749 | rs377054370 | 42877408 | 41505481 | T | C | intron_variant | 0 | 0 | 0 | 0 | 0 | 0 | 0 | 0 | 0 | 0 |
| 750 | - | 42877414 | 41505487 | G | A | intron_variant | 0 | 0 | 0 | 0 | 0 | 0.02 | 0 | 0 | 0 | 0 |
| 751 | rs147630170 | 42877432 | 41505505 | G | C | intron_variant | 0 | 0 | 0 | 0 | 0 | 0 | 0 | 0 | 0 | 0 |
| 752 | - | 42877453 | 41505526 | T | C | intron_variant | 0 | 0 | 0 | 0 | 0 | 0 | 0 | 0 | 0 | 0 |
| 753 | rs73357664 | 42877469 | 41505542 | A | G | intron_variant | 0 | 0 | 0 | 0 | 0 | 0 | 0 | 0 | 0 | 0 |
| 754 | rs548025657 | 42877481 | 41505554 | A | G | intron_variant | 0 | 0 | 0 | 0 | 0 | 0.02 | 0 | 0 | 0 | 0 |
| 755 | - | 42877542 | 41505615 | C | A | intron_variant | 0.1154 | 0.1154 | 0.2917 | 0.2308 | 0.3564 | 0.32 | 0.4667 | 0.4722 | 0.3843 | 0.2 |
| 756 | rs936471151 | 42877574 | 41505647 | A | G | intron_variant | 0 | 0 | 0 | 0 | 0 | 0 | 0.01111 | 0 | 0 | 0 |
| 757 | rs183758144 | 42877588 | 41505661 | A | G | intron_variant | 0 | 0 | 0 | 0 | 0 | 0 | 0 | 0 | 0 | 0 |
| 758 | rs115129572 | 42877673 | 41505746 | T | C | intron_variant | 0 | 0 | 0 | 0 | 0 | 0 | 0 | 0 | 0 | 0 |
| 759 | rs191526916 | 42877747 | 41505820 | T | C | intron_variant | 0 | 0 | 0 | 0 | 0.01485 | 0 | 0 | 0 | 0.00463 | 0 |
| 760 | - | 42878111 | 41506184 | A | G | intron_variant | 0 | 0 | 0 | 0 | 0.00495 | 0 | 0 | 0 | 0 | 0 |
| 761 | rs866793401 | 42878114 | 41506187 | A | T | intron_variant | 0 | 0 | 0 | 0 | 0 | 0 | 0 | 0 | 0 | 0.025 |
| 762 | rs28360562,COSV59821628 | 42878126 | 41506199 | C | A | intron_variant | 0 | 0 | 0.08333 | 0.1667 | 0.07426 | 0.08 | 0 | 0 | 0.01389 | 0.25 |
| 763 | - | 42878274 | 41506347 | C | T | intron_variant | 0 | 0 | 0 | 0 | 0 | 0 | 0 | 0 | 0.00463 | 0 |
| 764 | rs144458055,COSV59820956 | 42878469 | 41506542 | T | G | intron_variant | 0 | 0 | 0 | 0 | 0 | 0 | 0 | 0 | 0 | 0 |
| 765 | - | 42878471 | 41506544 | A | G | intron_variant | 0 | 0 | 0 | 0 | 0 | 0 | 0 | 0 | 0 | 0 |
| 766 | rs151189718 | 42878478 | 41506551 | T | C | intron_variant | 0 | 0 | 0 | 0 | 0 | 0 | 0 | 0 | 0 | 0 |
| 767 | rs746727496 | 42878493 | 41506566 | A | C | intron_variant | 0 | 0 | 0 | 0 | 0 | 0 | 0 | 0 | 0 | 0.025 |
| 768 | rs114641598 | 42878507 | 41506580 | C | T | intron_variant | 0 | 0 | 0 | 0 | 0 | 0 | 0 | 0 | 0 | 0 |
| 769 | | 42878611 | 41506684 | T | C | intron_variant | 0.2308 | 0.2308 | 0.02083 | 0.1923 | 0.1337 | 0.04 | 0.03333 | 0.05556 | 0.05556 | 0.175 |
| 770 | rs55704664 | 42878612 | 41506685 | C | T | intron_variant | 0 | 0 | 0 | 0.01282 | 0.0198 | 0.02 | 0.05556 | 0.05556 | 0.03704 | 0.025 |
| 771 | rs75200570,COSV59820977 | 42878626 | 41506699 | C | G | intron_variant | 0 | 0 | 0 | 0 | 0 | 0 | 0 | 0 | 0 | 0 |
| 772 | rs538465291 | 42878710 | 41506783 | A | C | intron_variant | 0 | 0 | 0 | 0 | 0 | 0 | 0 | 0 | 0 | 0 |
| 773 | rs150314077 | 42878761 | 41506834 | T | C | intron_variant | 0 | 0 | 0 | 0 | 0.00495 | 0 | 0 | 0 | 0 | 0 |
| 774 | rs538226704,COSV59824573 | 42878941 | 41507014 | G | A | intron_variant | 0 | 0 | 0 | 0 | 0 | 0 | 0 | 0 | 0 | 0 |
| 775 | rs149021153 | 42878991 | 41507064 | G | A | intron_variant | 0 | 0 | 0 | 0 | 0 | 0 | 0 | 0 | 0 | 0 |
| 776 | - | 42879072 | 41507145 | C | G | intron_variant | 0 | 0 | 0 | 0 | 0 | 0 | 0 | 0 | 0 | 0 |
| 777 | rs1601595157 | 42879098 | 41507171 | T | C | intron_variant | 0 | 0 | 0 | 0 | 0 | 0 | 0.01111 | 0 | 0 | 0 |
| 778 | rs912134074 | 42879182 | 41507255 | T | G | intron_variant | 0 | 0 | 0 | 0 | 0 | 0 | 0 | 0 | 0 | 0 |
| 779 | rs145900878 | 42879197 | 41507270 | T | A | intron_variant | 0 | 0 | 0 | 0 | 0 | 0 | 0 | 0 | 0 | 0 |
| 780 | rs968128742 | 42879236 | 41507309 | A | C | intron_variant | 0 | 0 | 0 | 0.03846 | 0.009901 | 0 | 0 | 0 | 0 | 0 |
| 781 | rs138498737,COSV59824580 | 42879283 | 41507356 | T | G | intron_variant | 0 | 0 | 0 | 0 | 0 | 0 | 0.01111 | 0 | 0 | 0 |
| 782 | - | 42879356 | 41507429 | A | G | intron_variant | 0 | 0 | 0 | 0 | 0 | 0 | 0 | 0 | 0 | 0 |
| 783 | rs57161767 | 42879403 | 41507476 | T | A | intron_variant | 0.03846 | 0.03846 | 0.2917 | 0.2436 | 0.2475 | 0.06 | 0.04444 | 0.02778 | 0.1296 | 0.1 |
| 784 | rs112467088 | 42879473 | 41507546 | C | T | intron_variant | 0 | 0 | 0 | 0 | 0 | 0.02 | 0 | 0 | 0 | 0 |
| 785 | rs578020496 | 42879604 | 41507677 | A | C | intron_variant | 0 | 0 | 0.1667 | 0.08974 | 0.1535 | 0.16 | 0.1222 | 0.05556 | 0.0463 | 0.1 |
| 786 | rs73230088,COSV59821273 | 42879688 | 41507761 | A | G | intron_variant | 0 | 0 | 0 | 0 | 0 | 0 | 0 | 0.02778 | 0 | 0 |
| 787 | rs573736906 | 42879690 | 41507763 | A | G | intron_variant | 0 | 0 | 0 | 0 | 0 | 0 | 0 | 0.02778 | 0 | 0 |
| 788 | rs1038320368 | 42879705 | 41507778 | T | A | intron_variant | 0 | 0 | 0 | 0 | 0 | 0.02 | 0 | 0 | 0 | 0 |
| 789 | - | 42879709 | 41507782 | C | G | intron_variant | 0 | 0 | 0 | 0 | 0 | 0 | 0 | 0 | 0 | 0 |
| 790 | - | 42879724 | 41507797 | A | G | intron_variant | 0.2308 | 0.2308 | 0.02083 | 0.1923 | 0.1238 | 0.04 | 0.03333 | 0.05556 | 0.05556 | 0.175 |
| 791 | rs8126497 | 42879747 | 41507820 | C | T | intron_variant | 0 | 0 | 0 | 0 | 0 | 0 | 0.01111 | 0 | 0 | 0 |
| 792 | rs371055210 | 42879841 | 41507914 | T | G | intron_variant | 0 | 0 | 0 | 0 | 0 | 0 | 0 | 0 | 0 | 0 |
| 793 | rs1031142674 | 42879909 | 41507982 | A | C | missense_variant | 0.03846 | 0.03846 | 0.3958 | 0.4103 | 0.3267 | 0.14 | 0.04444 | 0.02778 | 0.1435 | 0.35 |
| 794 | rs75603675,COSV59827803 | 42879910 | 41507983 | G | C | missense_variant | 0 | 0 | 0 | 0 | 0.009901 | 0 | 0 | 0 | 0 | 0 |

| | | | | | | | | | | | | | | | | |
|---|---|---|---|---|---|---|---|---|---|---|---|---|---|---|---|---|
| 795 | rs200291871,COSV59827230 | 42880296 | 41508369 | C | G | upstream_gene_variant | 0 | 0 | 0 | 0 | 0 | 0 | 0 | 0 | 0 | 0 |

**Supplementary Table 2:** Geographic region and numbers of samples examined in this study

| Geographic Region | Number of Samples (N) |
|---|---|
| America | 13 |
| Central Asia | 24 |
| Caucasus | 39 |
| Europe | 101 |
| **South Asia:-**<br>*Indo-European*<br>*Dravidian*<br>*Austroasiatic* | 25<br>18<br>4<br>3 |
| SEAM (Southeast Asia Mainland) | 18 |
| SEAI (Southeast Asia Island) | 45 |
| Siberia | 108 |
| West Asia | 20 |

**Supplementary Table 3a.** Distribution of major haplotypes of TMPRSS2 gene among the world population.

| Haplotype No. | AMR | SIB | CA | EUR | SEAI | SEAM | SA | CAU | WA |
|---|---|---|---|---|---|---|---|---|---|
| **Hap_34** | 0 | 2 | 3 | 0 | 3 | 1 | 1 | 0 | 0 |
| **Hap_48** | 0 | 0 | 1 | 5 | 0 | 0 | 2 | 0 | 3 |
| **Hap_75** | 0 | 4 | 0 | 12 | 0 | 0 | 0 | 2 | 0 |
| **Hap_98** | 0 | 9 | 0 | 2 | 0 | 0 | 0 | 3 | 0 |
| **Hap_260** | 0 | 17 | 0 | 0 | 3 | 0 | 1 | 0 | 0 |

**Supplementary Table 3b.** Frequency of haplotypes present in South Asia, among studied groups

| Haplotypes | AMR | SIB | CA | EUR | SEAI | SEAM | SA | CAU | WA | Total | AMR | SIB | CA | EUR | SEAI | SEAM | SA | CAU | WA |
|---|---|---|---|---|---|---|---|---|---|---|---|---|---|---|---|---|---|---|---|
| Hap_34 | 0 | 2 | 3 | 0 | 3 | 1 | 1 | 0 | 0 | 10 | - | 0.200 | 0.300 | - | 0.300 | 0.100 | 0.100 | - | - |
| Hap_48 | 0 | 0 | 1 | 5 | 0 | 0 | 2 | 0 | 3 | 11 | - | - | 0.091 | 0.455 | - | - | 0.182 | - | 0.273 |
| Hap_78 | 0 | 0 | 0 | 0 | 0 | 0 | 1 | 1 | 1 | 3 | - | - | - | - | - | - | 0.333 | 0.333 | 0.333 |
| Hap_112 | 0 | 0 | 0 | 0 | 0 | 0 | 1 | 1 | 0 | 2 | - | - | - | - | - | - | 0.500 | 0.500 | - |
| Hap_219 | 0 | 0 | 0 | 1 | 0 | 0 | 2 | 0 | 0 | 3 | - | - | - | 0.333 | - | - | 0.667 | - | - |
| Hap_245 | 0 | 0 | 0 | 0 | 0 | 0 | 1 | 0 | 0 | 1 | - | - | - | - | - | - | 1 | - | - |
| Hap_246 | 0 | 0 | 0 | 0 | 0 | 0 | 1 | 0 | 0 | 1 | - | - | - | - | - | - | 1 | - | - |
| Hap_247 | 0 | 0 | 0 | 0 | 0 | 0 | 1 | 0 | 0 | 1 | - | - | - | - | - | - | 1 | - | - |
| Hap_248 | 0 | 0 | 0 | 0 | 0 | 0 | 1 | 0 | 0 | 1 | - | - | - | - | - | - | 1 | - | - |
| Hap_249 | 0 | 0 | 0 | 0 | 0 | 0 | 1 | 0 | 0 | 1 | - | - | - | - | - | - | 1 | - | - |
| Hap_250 | 0 | 0 | 0 | 0 | 0 | 0 | 1 | 0 | 0 | 1 | - | - | - | - | - | - | 1 | - | - |
| Hap_251 | 0 | 0 | 0 | 0 | 0 | 0 | 1 | 0 | 0 | 1 | - | - | - | - | - | - | 1 | - | - |
| Hap_252 | 0 | 0 | 0 | 0 | 0 | 0 | 1 | 0 | 0 | 1 | - | - | - | - | - | - | 1 | - | - |
| Hap_253 | 0 | 0 | 0 | 0 | 0 | 0 | 1 | 0 | 0 | 1 | - | - | - | - | - | - | 1 | - | - |
| Hap_254 | 0 | 0 | 0 | 0 | 0 | 0 | 1 | 0 | 0 | 1 | - | - | - | - | - | - | 1 | - | - |
| Hap_255 | 0 | 0 | 0 | 0 | 0 | 0 | 1 | 0 | 0 | 1 | - | - | - | - | - | - | 1 | - | - |
| Hap_256 | 0 | 0 | 0 | 0 | 0 | 0 | 1 | 0 | 0 | 1 | - | - | - | - | - | - | 1 | - | - |
| Hap_257 | 0 | 0 | 0 | 0 | 0 | 0 | 1 | 0 | 0 | 1 | - | - | - | - | - | - | 1 | - | - |
| Hap_258 | 0 | 0 | 0 | 0 | 0 | 0 | 1 | 0 | 0 | 1 | - | - | - | - | - | - | 1 | - | - |
| Hap_259 | 0 | 0 | 0 | 0 | 0 | 0 | 1 | 0 | 0 | 1 | - | - | - | - | - | - | 1 | - | - |
| Hap_260 | 0 | 17 | 0 | 0 | 3 | 0 | 1 | 0 | 0 | 21 | - | 0.810 | - | - | 0.143 | - | 0.048 | - | - |
| Hap_261 | 0 | 0 | 0 | 0 | 0 | 0 | 1 | 0 | 0 | 1 | - | - | - | - | - | - | 1 | - | - |
| Hap_262 | 0 | 0 | 0 | 0 | 0 | 0 | 1 | 0 | 0 | 1 | - | - | - | - | - | - | 1 | - | - |
| Hap_263 | 0 | 0 | 0 | 0 | 0 | 0 | 1 | 0 | 0 | 1 | - | - | - | - | - | - | 1 | - | - |
| Hap_264 | 0 | 0 | 0 | 0 | 0 | 0 | 1 | 0 | 0 | 1 | - | - | - | - | - | - | 1 | - | - |
| Hap_265 | 0 | 0 | 0 | 0 | 0 | 0 | 1 | 0 | 0 | 1 | - | - | - | - | - | - | 1 | - | - |
| Hap_266 | 0 | 0 | 0 | 0 | 0 | 0 | 1 | 0 | 0 | 1 | - | - | - | - | - | - | 1 | - | - |
| Hap_267 | 0 | 0 | 0 | 0 | 0 | 0 | 1 | 0 | 0 | 1 | - | - | - | - | - | - | 1 | - | - |
| Hap_268 | 0 | 0 | 0 | 0 | 0 | 0 | 1 | 0 | 0 | 1 | - | - | - | - | - | - | 1 | - | - |
| Hap_269 | 0 | 0 | 0 | 0 | 0 | 0 | 1 | 0 | 0 | 1 | - | - | - | - | - | - | 1 | - | - |
| Hap_270 | 0 | 0 | 0 | 0 | 0 | 0 | 1 | 0 | 0 | 1 | - | - | - | - | - | - | 1 | - | - |
| Hap_271 | 0 | 0 | 0 | 0 | 0 | 0 | 2 | 0 | 0 | 2 | - | - | - | - | - | - | 1 | - | - |
| Hap_272 | 0 | 0 | 0 | 0 | 0 | 0 | 1 | 0 | 0 | 1 | - | - | - | - | - | - | 1 | - | - |
| Hap_273 | 0 | 0 | 0 | 0 | 0 | 0 | 1 | 0 | 0 | 1 | - | - | - | - | - | - | 1 | - | - |
| Hap_274 | 0 | 0 | 0 | 0 | 0 | 0 | 1 | 0 | 0 | 1 | - | - | - | - | - | - | 1 | - | - |
| Hap_275 | 0 | 0 | 0 | 0 | 0 | 0 | 1 | 0 | 0 | 1 | - | - | - | - | - | - | 1 | - | - |
| Hap_276 | 0 | 0 | 0 | 0 | 0 | 0 | 1 | 0 | 0 | 1 | - | - | - | - | - | - | 1 | - | - |
| Hap_277 | 0 | 0 | 0 | 0 | 0 | 0 | 1 | 0 | 0 | 1 | - | - | - | - | - | - | 1 | - | - |
| Hap_278 | 0 | 0 | 0 | 0 | 0 | 0 | 1 | 0 | 0 | 1 | - | - | - | - | - | - | 1 | - | - |
| Hap_279 | 0 | 0 | 0 | 0 | 0 | 0 | 1 | 0 | 0 | 1 | - | - | - | - | - | - | 1 | - | - |
| Hap_280 | 0 | 0 | 0 | 0 | 0 | 0 | 1 | 0 | 0 | 1 | - | - | - | - | - | - | 1 | - | - |
| Hap_281 | 0 | 0 | 0 | 0 | 0 | 0 | 1 | 0 | 0 | 1 | - | - | - | - | - | - | 1 | - | - |
| Hap_282 | 0 | 0 | 0 | 0 | 0 | 0 | 1 | 0 | 0 | 1 | - | - | - | - | - | - | 1 | - | - |
| Hap_283 | 0 | 0 | 0 | 0 | 0 | 0 | 1 | 0 | 0 | 1 | - | - | - | - | - | - | 1 | - | - |
| Hap_284 | 0 | 0 | 0 | 0 | 0 | 0 | 1 | 0 | 0 | 1 | - | - | - | - | - | - | 1 | - | - |
| Hap_285 | 0 | 0 | 0 | 0 | 0 | 0 | 1 | 0 | 0 | 1 | - | - | - | - | - | - | 1 | - | - |
| Hap_286 | 0 | 0 | 0 | 0 | 0 | 0 | 1 | 0 | 0 | 1 | - | - | - | - | - | - | 1 | - | - |

**Supplementary Table 4a**. Statewise allele frequency of studied SNP for the association

| State | Population Groups | Sanple Size(N) | rs2070788(G) | rs734056(A) | rs12329760(T) | rs2276205(G) | rs3787950(C) | Refrences |
|---|---|---|---|---|---|---|---|---|
| Arunanchal Pradesh | Nyshi_Bom_Mro_Oraon | 28 | 0.2143 | 0.125 | 0.3214 | 0.05357 | 0.1607 | present study |
| Bihar | Brahmin Bihar | 8 | 0.5 | 0.4375 | 0.3125 | 0.125 | 0.0625 | present study |
| Gujarat | Gujarati and Brahmin_Guj | 121 | 0.4835 | 0.5 | 0.1694 | 0.03719 | 0.2521 | 1000 Genome |
| Haryana | Ror and Brahmin_Har | 18 | 0.3056 | 0.3056 | 0.3333 | 0.1471 | 0.1765 | Pathak et al. 2018 |
| Jharkhand | Santhal and Oraon | 51 | 0.3137 | 0.2059 | 0.3431 | 0.01961 | 0.2843 | present study |
| Kerala | Cochin_Jews | 8 | 0.25 | 0.1875 | 0.0625 | 0 | 0 | present study |
| Meghalaya | Garo and Khasi | 16 | 0.4062 | 0 | 0.1562 | 0.09375 | 0.1875 | Tatte et al. 2019 and present study |
| Rajasthan | Gujjar | 15 | 0.4 | 0.3667 | 0.3667 | 0.1 | 0.2 | Pathak et al. 2018 |
| Tamilnadu | Brahmins_TN | 15 | 0.4333 | 0.4667 | 0.3 | 0.06667 | 0.2667 | 1000 Genome and present study |
| Tripura | Tripuri | 20 | 0.3 | 0.05 | 0.175 | 0.1 | 0.025 | present study |
| Uttar Pradesh | Brahmins and Kshatriya | 89 | 0.483 | 0.4494 | 0.2303 | 0.0618 | 0.1818 | present study |
| West Bengal | Bengali and Brahmin_WB | 19 | 0.4737 | 0.4211 | 0.1579 | 0.07895 | 0.2105 | present study |
| Maharashtra | Maratha and Mumbai Jews | 13 | 0.3846 | 0.3077 | 0.3462 | 0.1154 | 0.2308 | Chaubey et al. 2017 and present study |
| Manipur | Manipuri Brahmin | 29 | 0.475 | 0.2895 | 0.25 | 0.1 | 0.2 | present study |

**Supplementary Table 4b**. Statewise epidemiological data of cases, deaths and CFR.

| | (June_2021) | | | (July_2021) | | | (August_2021) | | |
|---|---|---|---|---|---|---|---|---|---|
| **State** | **Cases** | **Death** | **CFR** | **Cases** | **Death** | **CFR** | **Cases** | **Death** | **CFR** |
| Arunanchal Pradesh | 41646 | 197 | 0.004730346 | 30850 | 138 | 0.004473258 | 50070 | 242 | 0.00483323 |
| Bihar | 723695 | 9625 | 0.013299802 | 716296 | 9466 | 0.013215207 | 725192 | 9646 | 0.01330131 |
| Gujarat | 824384 | 10074 | 0.012220033 | 819376 | 9985 | 0.012186103 | 825045 | 10077 | 0.01221388 |
| Haryana | 769417 | 9578 | 0.012448386 | 765096 | 8904 | 0.011637755 | 770079 | 9649 | 0.01252988 |
| Jharkhand | 346533 | 5120 | 0.014774928 | 343065 | 5082 | 0.014813519 | 347392 | 5130 | 0.01476718 |
| Kerala | 3117083 | 15025 | 0.004820212 | 2702823 | 10804 | 0.003997302 | 3552525 | 17747 | 0.0049956 |
| Meghalaya | 56052 | 926 | 0.016520374 | 41100 | 714 | 0.017372263 | 69171 | 1163 | 0.0168134 |
| Rajasthan | 953257 | 8947 | 0.009385717 | 949008 | 8799 | 0.009271787 | 953827 | 8954 | 0.00938745 |
| Tamilnadu | 2528806 | 33606 | 0.013289276 | 2324597 | 28906 | 0.012434844 | 2575308 | 34317 | 0.0133254 |
| Tripura | 72860 | 722 | 0.009909415 | 58521 | 604 | 0.010321081 | 80211 | 770 | 0.00959968 |
| Uttar Pradesh | 1707655 | 22705 | 0.013296011 | 1701668 | 21667 | 0.012732801 | 1708772 | 22773 | 0.01332711 |
| West Bengal | 1515599 | 17970 | 0.011856698 | 1452987 | 16731 | 0.0115149 | 1533803 | 18229 | 0.01188484 |
| Maharashtra | 6189257 | 126560 | 0.020448335 | 5887853 | 106367 | 0.018065499 | 6353328 | 133996 | 0.02109068 |
| Manipur | 81560 | 1340 | 0.016429622 | 58768 | 944 | 0.016063164 | 104324 | 1650 | 0.01581611 |

**Supplementary Table 5.** Results of the linear regression and Pearson correlation coefficient tests performed for statistical significance**.**

| **Observation** | **Linear regression** | | **Pearson's correlation** | |
|---|---|---|---|---|
| **rs2070788** | **R square** | **p-value** | **r** | **p-value** |
| June_CFR | 0.3382 | 0.0292 | 0.582 | 0.029 |
| July_CFR | 0.3097 | 0.0387 | 0.557 | 0.039 |
| August_CFR | 0.2888 | 0.0475 | 0.537 | 0.047 |

| **Observation** | **Linear regression** | | **Pearson's correlation** | |
|---|---|---|---|---|
| **rs734056** | **R square** | **p-value** | **r** | **p-value** |
| June 2021_CFR | 0.01111 | 0.7198 | 0.105 | 0.720 |
| July 2021_CFR | 0.02769 | 0.5697 | 0.166 | 0.570 |
| August 2021_CFR | 0.02627 | 0.5798 | 0.162 | 0.580 |

| **Observation** | **Linear regression** | | **Pearson's correlation** | |
|---|---|---|---|---|
| **rs12329760** | **R square** | **p-value** | **r** | **p-value** |
| June 2021_CFR | 0.05556 | 0.4172 | 0.236 | 0.417 |
| July 2021_CFR | 0.07899 | 0.3304 | 0.281 | 0.330 |
| August 2021_CFR | 0.08088 | 0.3244 | 0.284 | 0.324 |

| **Observation** | **Linear regression** | | **Pearson's correlation** | |
|---|---|---|---|---|
| **rs2276205** | **R square** | **p-value** | **r** | **p-value** |
| June 2021_CFR | 0.1834 | 0.1265 | 0.428 | 0.127 |
| July 2021_CFR | 0.1896 | 0.1197 | 0.435 | 0.120 |
| August 2021_CFR | 0.1839 | 0.1261 | 0.429 | 0.126 |

| **Observation** | **Linear regression** | | **Pearson's correlation** | |
|---|---|---|---|---|
| **rs3787950** | **R square** | **p-value** | **r** | **p-value** |
| June 2021_CFR | 0.2650 | 0.0596 | 0.515 | 0.060 |
| July 2021_CFR | 0.2782 | 0.0526 | 0.527 | 0.053 |
| August 2021_CFR | 0.2802 | 0.0516 | 0.529 | 0.052 |

**Supplementary Table 6:** Aggregate haplotypes frequency carrying rs2070788 (G allele).

| | AMR | CA | CAU | EUR | SA | SEAI | SEAM | SIB | WA |
|---|---|---|---|---|---|---|---|---|---|
| **Haplotype Freq.** | 0.654 | 0.354 | 0.474 | 0.426 | 0.560 | 0.322 | 0.389 | 0.486 | 0.6 |

**Supplementary Table 7** | rs2070788 frequency in 1000 genome data

| Region | rs2070788 | |
|---|---|---|
| | Ref. Allele (G) | Alt. Allele (A) |
| African | 0.2738 | 0.7262 |
| East Asian | 0.3562 | 0.6438 |
| Europe | 0.4642 | 0.5358 |
| South Asian | 0.466 | 0.534 |
| American | 0.494 | 0.506 |
